# Gene expression divergence between locally adapted inland annual and coastal perennial ecotypes of *Mimulus guttatus* across developmental stages

**DOI:** 10.1101/2024.11.04.621855

**Authors:** Jason D. Olsen, Billie A Gould, Yani Chen, David B. Lowry

## Abstract

The action of natural selection across heterogeneous natural landscapes drives local adaptation and the formation of plant ecotypes, the precursors to new species. Plant ecotypes typically differ significantly in morphology, physiology, and development, yet our understanding of their underlying genetic basis remains limited. Despite their importance, studies of the molecular underpinnings of ecotypic divergence through developmental stages are rare. Here, we compared gene expression at different developmental time points between ecologically reproductively isolated coastal perennial and inland annual ecotypes of the yellow monkeyflower, *Mimulus guttatus*. We took two major approaches to understand differences in gene regulation between the ecotypes at the level of gene networks. First, we evaluated expression variation between the ecotypes in candidate molecular pathways. Next, we conducted gene co-expression network analyses to identify new candidate mechanisms driving ecotypic divergence. Overall, we found significant differences in global expression between the ecotypes and across developmental stages. Among the candidate pathways we evaluated, genes in the jasmonic acid pathway were the most significantly enriched for divergent gene expression. This includes the most differentially expressed gene in our analyses, which is a key gene (cytochrome P450 CYP94B1) involved in the degradation of bioactive jasmonic acid. Our gene co-expression network analysis revealed different but complementary insights into the differential regulation of genes between the ecotypes, especially at a more microscopic level of these organisms. Overall, our research further supports the hypothesis that plant hormone pathways play a crucial role in the evolution of plant ecotypes and, consequently, the emergence of new species.

## INTRODUCTION

The evolution of new species generally occurs over long periods of time and involves a combination of geographic isolation, natural selection, and genetic drift (Coyne & Orr 2004). In plants, speciation is thought to often be facilitated through ecogeographic isolation among populations driven by local adaptation (Schemske 2000; Sobel et al. 2010; Sobel 2014; Butlin & Faria 2024). This process leads to the formation of ecotypes, which initially are reproductively isolated by ecological isolating barrier (Lowry 2012; Butlin & Faria 2024). These ecotypes generally have higher fitness in their native habitats, which limits gene flow between habitats and, in turn, facilitates the buildup of greater divergence and reproductive isolating mechanisms (Sobel et al. 2010; Lowry 2012). In this way, ecotypes can be viewed as intermediates in the process of speciation.

Some of the greatest progress in our understanding of the evolution of plant ecotypes comes from studies of the adaptive divergence between coastal and inland plant ecotypes. Indeed, Turesson’s (1922a,b) ecotype concept was largely inspired by the high level of morphological divergence between coastal and inland populations of plants in Sweden. Clausen, Keck, and Hiesey famously followed up on the work of Turesson through multiple monograph-length studies of coastal and inland ecotypes of multiple plant species in California (Clausen et al. 1940, 1948). Since that time, several other researchers have found similar patterns of differentiation between coastal and inland ecotypes worldwide (Boyce 1954; Nagy & Rice 1997; Foster et al. 2007; Verhoeven et al. 2008; Itoh et al. 2024). From these studies, some major generalizations can be made about ecotypic differentiation associated with coastal habitats (Lowry 2012).

Coastal ecotypes are generally more dwarfed in stature, prostrate in their growth form, and flower later than inland ecotypes of the same species. Boyce (1954) argued that much of this growth form difference was due to coastal plant adaptations to oceanic salt spray, and suggested that coastal ecotypes might be better referred to as salt spray ecotypes.

While the concept of plant ecotypes was first formulated by Turesson (1922a,b) over a century ago, our understanding of the genetic, physiological, and developmental mechanisms underlying the evolution of ecotypes remains limited. Some of the largest advances in our understanding of ecotype evolution have come from the coastal ecotype systems, where molecular genetic and genomic tools have recently become available. For example, recent molecular research on the evolution of dune and headland populations of *Senecio latifolius* on the east coast of Australia has revealed many clues about the evolution of plant ecotypes (Roda et al. 2013; James et al. 2021, 2023; Wilkinson et al. 2021; Broad et al. 2024). In *S. latifolius*, the headland ecotype experiences intense winds, and presumably salt spray, which in combination with the availability of steep cliff habitat has driven a dwarfed and prostrate growth architecture. In contrast, the dune ecotype is taller with an upright growth habit, which may reflect a more sheltered habitat where this growth form is advantageous. Recent genetic research on this system suggests a complex genetic basis of the differences in physiology and development between the dune and headland ecotypes. Despite this complexity, there is strong evidence for a role of evolutionary changes in the auxin hormone pathway being involved in the changes in growth form, especially with a large divergence in gravitropism between the ecotypes (Wilkinson et al. 2021; James et al. 2023; Broad et al. 2024).

Given the major role that plant hormone pathways play in shaping the growth form and physiology of plants, it is perhaps not surprising that they may also play a key role in the evolution of plant ecotypes (Voesenek & Blom 1996; VanWallendael et al. 2019; Broad et al. 2024). Recent research has shown that hormone pathways interact with each other through complex crosstalk through which they coordinate development and stress responses (Fujita et al. 2006; Depuydt & Hardtke 2011; Murphy 2015; Aerts et al. 2021; Gasperini & Howe 2024). For example, the jasmonic acid and gibberellin hormone pathways are thought to coordinate plant trade-offs between growth and defense at least partially through the interaction of JASMONATE ZIM-domain (JAZ) and DELLA genes in each of these hormones’ respective signaling pathways (Pieterse et al. 2014; Howe et al. 2018; Guo et al. 2018). While we are still far from understanding the highly complex network of plant hormone interactions, our current knowledge of these pathways provides great utility for gaining new insights into how plant ecotypes evolve through local adaptation to different habitats.

To better understand how hormone and other key pathways have shifted in the evolution of plant ecotypes, we conducted a study of gene expression divergence between coastal and inland ecotypes through developmental stages, from the first set of true leaves to the emergence of floral buds. The focus of our study was on coastal and inland ecotypes in the yellow monkeyflower, *M. guttatus*. Over the past two decades, extensive research has been conducted to understand the evolutionary mechanisms underlying the divergence of inland annual versus coastal perennial ecotypes in this species (Hall et al. 2006, 2010; Lowry et al. 2008, 2009; Lowry & Willis 2010; Twyford & Freidman 2015; Twyford et al. 2015; Gould et al. 2017, 2018; Popovic & Lowry 2020; Kollar et al. 2025). There are large differences in a wide swath of traits between coastal perennial and nearby inland annual ecotypes (Abrams 1951; Hitchcock and Cronquit 1973; Vickery 1978; Lowry et al. 2008). These ecotypes diverge in traits such as growth rate, morphology, phenology, and resistance to herbivory, and are locally adaptative in reciprocal transplant experiments (Hall and Willis 2006; Van Kleunen 2007; Lowry et al. 2008; Hall et al. 2010; Lowry et al. 2019). In general, the inland annual ecotype flowers early to avoid seasonal drought and produces fewer true leaves before developing floral buds. The coastal perennial ecotype flowers later and produces more leaf pairs before budding. Based on the morphological distinctness of the coastal perennial ecotype, some taxonomists have raised it to the species level, naming it the magnificent monkeyflower, *Mimulus grandis* (syn. *Erythranthe grandis*; Nesom 2012). However, given the high level of allele sharing between coastal perennial and inland annual groups of populations (only four fixed nucleotide differences out of >29 million SNPs; Gould et al. 2017), we more cautiously treat each group as an ecotype instead of a species. While we don’t consider the coastal perennial and inland annual ecotypes as distinct species, we do recognize that they are on the speciation continuum, as they have acquired a significant level of ecologically-based reproductive isolation (Lowry et al. 2008; Lowry & Willis 2010).

Prior research on the coastal perennial and inland annual ecotypes suggests that their morphological divergence is largely driven by evolutionary shifts in hormone pathways. This hypothesis first emerged as a result of a prior population genomic outlier study (Gould et al. 2017), which identified genes in the gibberellin hormone pathway as top candidate genes for ecotype divergence. One of those candidate genes, GA20ox2, a key enzyme in the gibberellin biosynthetic pathway, is located in a chromosomal inversion polymorphism on chromosome 8 that is responsible for a substantial portion of the divergence between coastal perennial and inland annual ecotypes (Lowry & Willis 2010; Friedman 2014; Lowry et al. 2019; Blanchard et al. 2024). A recent study found that GA20ox2 is not expressed in leaves; however, it exhibits higher expression in the shoot apices of inland annuals compared to coastal perennials (Kollar et al., 2025). This differential expression could initiate a developmental cascade that impacts the expression of other important pathways in developing leaves, branches, and reproductive structures, which may explain some of the broad phenotypic effects associated with the chromosome 8 inversion. Indeed, the application of gibberellin (GA3) to coastal perennial plants leads to substantial phenotypic changes, converting their growth architecture from prostrate to upright (Lowry et al. 2019) and making them more susceptible to oceanic salt spray in the field (Toll et al. 2024). Beyond its significant morphological and phenological effects, the chromosome 8 inversion also contributes to elevated constitutive and inducible levels of defensive phenylpropanoid glycoside (PPG) in coastal perennials versus inland annual plants (Lowry et al. 2019; Blanchard et al. 2024). This finding suggests the involvement of the jasmonic acid (JA) hormone pathway in the differentiation of coastal and inland plants, as PPG production is known to be stimulated by jasmonic acid. Given the known cross-talk between the gibberellin and jasmonic acid pathways (Fujita et al. 2006; Yang et al. 2012; Wasternack 2017; Züst & Agrawal 2017; Guo et al. 2018), evaluation of expression patterns for gene in those pathways could provide further insights into the mechanisms underlying the divergence of the ecotypes.

In this study, we analyzed patterns of gene expression between coastal perennial and inland annual accessions of *M. guttatus* across development, from the production of the first pair of true leaves to the emergence of floral buds. The primary goal of this work was to understand how coastal perennial and inland annual ecotypes are differentiated at the level of gene networks. To that end, we took two complementary approaches. First, we evaluated how individual genes were differentially regulated between ecotypes and across developmental stages, across all of the major plant hormone pathways. Based on our prior work, we hypothesized that the gibberellin and jasmonic acid pathways would have an elevated level of differential gene expression between the ecotypes. In addition to hormone pathways, we evaluated gene expression in the Salt Overly Sensitive (SOS) pathway, as a few key genes in this pathway were implicated in divergence between the ecotypes in prior work (Gould et al. 2018). Following our analyses of individual genes in candidate pathways, we conducted a whole-genome transcriptional module analysis using a Weighted Gene Co-expression Network Analysis (WGCNA; Langfelder and Horvath 2008). This analysis did not incorporate any prior knowledge about the *M. guttatus* system and was thus effective at generating additional new hypotheses of mechanisms underlying divergence of the ecotypes. Overall, we found that the two approaches to understanding gene expression at the network level provided different, yet complementary, insights into ecotype divergence.

## METHODS

### Plant material

To identify patterns of gene expression divergence between the ecotypes, we selected one coastal perennial (SWB-4) and one inland annual (LMC-17) accession for a detailed study throughout development. The coastal perennial SWB population (N 39.02.159, W 123.41.428) is located near Manchester, CA, while the inland annual population LMC (N 38.51.839, W 123.05.035) is located near Boonville, CA. By focusing on two accessions, we were able to achieve a high level of replication for each developmental stage. Seeds from each accession consisted of self-fertilized families that were two generations inbred in the laboratory before the experiments. To eliminate potential maternal effects, seeds were grown for one additional generation at Michigan State University (MSU, East Lansing, MI) in the four months before the growth chamber experiment, where tissue was collected for gene expression analysis.

### Plant growth conditions

Surface-sterilized *Mimulus guttatus* seeds were placed on 60 mm diameter x 15mm height Petri dishes containing half-strength Linsmaier and Skoog (LS) medium: 1.5% w/v sucrose and 0.25% Phytagel (P8169, Sigma, St. Louis, MO). Seeds were then stratified at 4°C in the dark for 14 days. After cold treatment, plants were germinated at 20°C under a 16 h 100 mE light / 8 h dark cycle for 5 to 14 days in a Percival (Percival Scientific, Perry, IA) chamber. Cotyledon-stage seedlings were transferred to 3.5-inch square pots containing SureMix Perlite (Michigan Grower Products, Galesburg, MI) and grown in a BioChambers FXC-19 flex growth chamber (BioChambers, Winnipeg, Manitoba) in the same conditions as in the Percival chamber. Three key steps were taken to minimize the effects of environmental heterogeneity within the growth chamber. First, the locations of plants were fully randomized across trays within the chamber. Second, the tray position within the chamber was haphazardly rotated daily. Third, the developmental stage at which each plant was sampled was assigned randomly. Leaf pairs and floral buds were collected in a time series (Fig. S1). Leaves were collected at the point in development when they were nearing maximal expansion. The floral buds were collected from plants around the time of anthesis. Only one tissue was collected per plant to avoid induced responses and repeated measures. Each tissue collection was harvested in a fixed time window (14:00 to 14:30) during the simulated daylight period. Tissue was immediately flash frozen on liquid nitrogen and stored in a -80° C freezer. RNA was later extracted from samples using a Spectrum Plant Total RNA Kit (Sigma-Aldrich, St. Louis, MO). We collected 10 biological replicates for each developmental stage x ecotype combination (*N =* 80 total tissue samples). For the inland annual plants, we sampled at the two-leaf, four-leaf, and floral bud stages because plants often flowered after producing two pairs of true leaves. Because the coastal perennial plants flowered at a later developmental stage, we sampled at the two-leaf, four-leaf, six-leaf, eight-leaf, and floral bud stages. All tissue samples were collected before anthesis.

### Library preparation and sequencing, and processing

We prepared genomic libraries using a 3’-Tag-Seq procedure, which restricts sequencing to the 3’ end of mRNAs and dramatically reduces the cost for RNA-seq studies (Meyer et al. 2011; Lohman et al. 2016; Marx et al. 2020). Our protocol for library preparation followed an updated version of the methods of Meyer et al. (2011), which had been adapted for sequencing on the Illumina platform, as described in Weng & Juenger (2022). The detailed protocol of library preparation for our study can be found in the Supplemental Methods. Each sample was assigned a fully randomized position across four pools of 20 samples before library construction. The MSU Genomics Core Facility conducted sequencing in Illumina HiSeq 2500 Rapid Run flow cells (v2) in the 1x100bp conformation.

Sequencing of the four pooled libraries was conducted across three different flow cells, with two pools being sequenced in two separate flow cells and the other pools only being sequenced on a single flow cell each.

Raw Illumina reads were processed following our protocol that is available on GitHub along with code for other analyses detailed below (github.com/lowrylab/Mimulus_expression_across_development). Briefly, raw reads were first examined for quality with FastQC (bioinformatics.babraham.ac.uk/projects/fastqc/), which found a median Phred score of ∼37 across libraries. Adapter sequences were then removed with Cutadapt (Martin 2011). Cutadapt was applied with an adapter mismatch error threshold of 20% of primer length. Primer trimming began with the 5’ end (NNNN followed by 3-5 G’s with a first step to cut 8 bases and a second step to cut the remaining 5’ G’s). Trimming of the 3’ end was done by removing 15bp of the 3’ end of each read to remove the poly-A primer and any readthrough. To retain only high-quality sequences after trimming, any bases on either end with a quality score < 30 were removed, and we required the last two bases of each read to match the poly-A primer while discarding any reads <30bp in length. These parameters align with the requirements for the Tag-Seq methodology to remove both adapter sequences and non-template bases created during library preparation (Meyer et al. 2011). Trimmed sequence files were concatenated and aligned to the *Mimulus guttatus* v2.0 reference genome (Hellsten et al. 2013; phytozome-next.jgi.doe.gov) using the memory-efficient Burrows-Wheeler Aligner (BWA-MEM; Li 2013), as in Marx et al. (2020) and Weng & Juenger (2022). The v2.0 reference genome was selected for alignment because genes are well annoatated and it has been used for many other recent studies. Aligned reads were filtered to include only high-quality, primary alignments using Samtools 1.2 (Li et al. 2009). The total number of aligned reads retained for each sample was calculated with Samtools. Counting the number of reads aligned per gene per sample was done with HT-seq v.0.6.1 (Anders et al. 2015) with “stranded” and “union” parameter options. Count files were filtered to remove genes with < 5 aligned reads to exclude non-expressed genes or alignment artifacts.

### Clustering analyses of gene expression

To evaluate whether samples clustered based on genotype, developmental stage, and/or library pool, we conducted a principal component analysis (PCA) with DESeq2 (Love et al. 2014) in R version 4.4.2. Before the implementation of normalization and PCA, we filtered out low abundance genes by removing any genes did not have more than 10 counts in more than 5 samples. We then normalized the remaining data using a variance stabilizing transformation (VST). The PCA was conducted with this filtered and normalized data (Fig. 1). We focused on the top two PCs, as they explained far more of the variation than any of the other PCs (Fig. S2). In addition to the PCA, we evaluated clustering of the same VST data with a heat map produced by the R pheatmap function.

**Figure 1:**
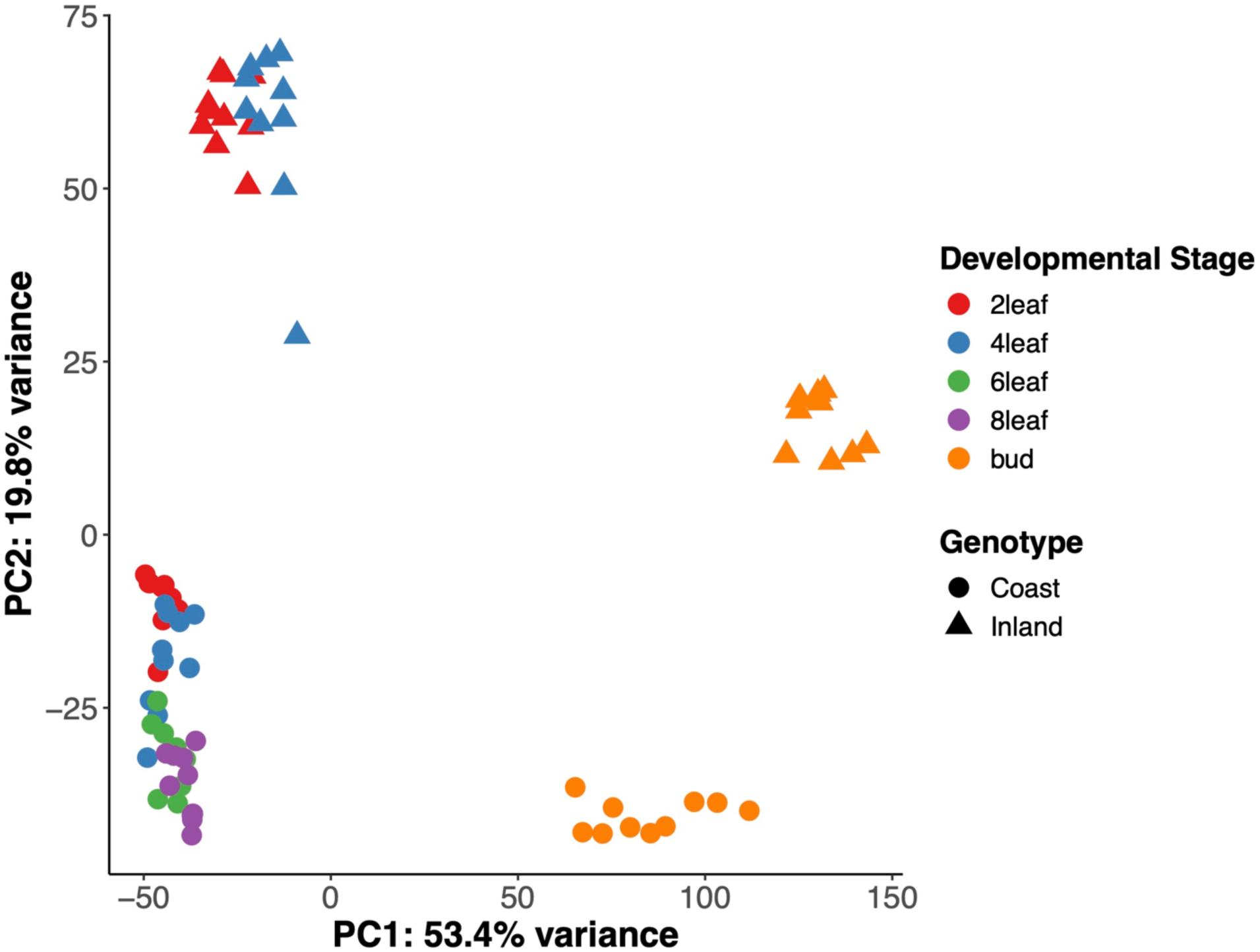
Principal component analysis of pre-filtered, variance-stabilizing transformed (VST) transcript count data from RNASeq. Different colors denote all combinations of developmental stages and ecotypes. The first principal axis (PC1) explains the variance in the dataset and primarily distinguishes differences between leaf (vegetative) and bud (reproductive) stages. The second principal axis explains the variance that primarily distinguishes differences between ecotypes.

### Differential gene expression analyses at the individual gene level

To evaluate differential expression analyses at the individual gene level, we used the edgeR-voom-limma framework (Law et al. 2014), implemented in R version 4.5.0. Voom-limma was selected for this analysis because it stabilizes the mean-variance relationships of count data for linear modeling and allows for complex experimental designs with multiple interactions. Our analysis of differential gene expression focused on the 2-Leaf, 4-Leaf, and bud developmental stages, as inland annual plants generally did not produce 6-Leaf and 8-Leaf tissue. Prior to model fitting, we filtered the raw count data using the filterByExpr function, which is part of the edgeR package (version 4.6.1; Robinson et al., 2010). This function retains only genes that have sufficient counts to achieve statistical significance through differential expression analysis, using the default parameters. We used version 3.64.0 of limma for this analysis. Normalization was then conducted for count data using the calcNormFactors function, which accounts for composition bias and library size differences. This normalized data was then converted to log2-counts per million (log2-CPM) using the voom transformation.

The following linear model was fit for each gene:

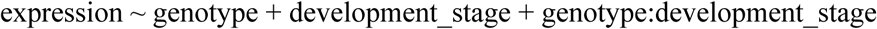

Empirical Bayes methods were used to shrink gene-wise variances, which improves the reliability of inference for genes with highly variable transcript abundance. The “Inland” genotype and “2leaf” developmental stage were set as the reference levels for statistical contrasts. We tested for differential expression of genes for five contrasts: 1) Main effect of genotype (Coast vs. Inland), 2) main effect of developmental stage for 4leaf vs. 2leaf), 3) main effect of developmental stage for bud vs. 2leaf, 4) interaction between genotype and developmental stage for 4leaf vs. 2leaf, and 5) interaction between genotype and developmental stage for bud vs. 2leaf. We controlled for multiple testing by calculating False Discovery Rate (FDR) adjusted *P*-values using the Benjamini-Hochberg method. To evaluate how closely overall gene expression analyzed with the voom-limma approach mirrored the analysis of all genes using DESeq2, we output a multidimensional scaling (MDS) plot for the 2leaf, 4leaf, and bud stages.

### Analysis of candidate pathways

To evaluate whether any hormone pathways were enriched for differentially expressed genes, we examined the expression of core genes involved in hormone pathways. These core pathway genes for each of the major hormone pathways (Table 1) were identified by custom Python scripts to parse the *Mimulus guttatus* v2.0 genome annotation file. Gene lists identified by these scripts were further refined through close inspection, followed by the removal of genes of lower confidence. We used the gene lists that we compiled for those pathways to extract the gene expression results for each gene from our individual gene analyses (above). Once we had identified the number of differentially expressed genes (FDR adjusted *P* < 0.05) and highly differenitally expressed genes (Log-Fold Change (LFC) > 2), we tested for enrichment with Fisher-Exact tests, with Benjamini-Hochberg FDR adjusted correction for multiple testing. In addition to hormone pathway genes, we also evaluated key genes in plant cellular homeostasis, including genes in the Salt Overly Sensitive (SOS) pathway, given prior evidence that these genes may be important for coastal adaptations in this system (Gould et al., 2017, 2018). As with the hormone pathway genes, we used custom python scripts to identify candidate genes from the v2.0 annotation file.

**Table 1:** Differential gene expression of hormone pathway genes between coastal and inland ecotypes and between the floral bud and second pair of true leaves (2-leaf) developmental stages.

| Hormone Pathway | Candidate Genes | Genes Evaluated | Ecotype DE | Ecotype LogFC > 2 | Bud vs 2-Leaf DE | Bud vs 2-Leaf LogFC > 2 |
| --- | --- | --- | --- | --- | --- | --- |
| Gibberellin | 64 | 19 | 12 | 4 | 14 | 6 |
| Jasmonic Acid | 82 | 49 | 40*** | 23*** | 35 | 15 |
| Absciscic Acid | 102 | 64 | 27 | 1 | 42 | 14 |
| Brassinosteroid | 49 | 29 | 16 | 1 | 20 | 5 |
| Auxin | 271 | 111 | 60 | 8 | 90 | 27 |
| Cytokinin | 99 | 38 | 23 | 2 | 23 | 4 |
| Ethylene | 61 | 28 | 14 | 1 | 23 | 5 |
| Salicylic Acid | 27 | 6 | 1 | 1 | 5 | 0 |
| Strigolactone | 13 | 7 | 2 | 0 | 3 | 1 |
| Genome-wide | 28140 | 13920 | 7238 | 926 | 10248 | 2347 |
| ***Significant enrichment at an FDR adjusted $P < 0.001$ | | | | | | |

**Table 2:** Hypothesized biological function and list of key hub genes for each of the nine modules identified by WGCNA analyses.

| Module | Biological function prediction based on GO enrichment and hub genes | Key hub genes |
| --- | --- | --- |
| Red | Vesicle trafficking and membrane dynamics, with a possible roles in stress responses. Coordinates phospholipid signaling, vesicle formation, membrane identity, and trafficking processes | Phospholipase D (PLD; Migut.B01671), an important signaling molecule that promotes membrane curvature and facilitates vesicle fusion and fission. Sec1/munc18-like (SM) protein (Migut.E01133), which is involved in vesicle trafficking. Nudix hydrolase 19 (Migut.J01169), which is a regulator of cellular redox homeostasis, crucial for plant stress responses. |
| Tan | Coordination of plant hormone responses with metabolism, lipid synthesis, posttranslational modifications, and cell wall remodeling. Likely important in growth and development. | Cellulase glycosyl hydrolase (Migut.A00708) and sucrose phosphate synthase (Migut.E01808) are involved in coordination between sugar metabolism and cell wall remodeling. Three of the hub genes were hormone genes involved in jasmonic acid biosynthesis (OPR1, Migut.B00746), gibberellin response (Migut.E01377), and auxin signaling (GH3, Migut.N02078). |
| Turquoise | Cell wall modifications, membrane dynamics, secretory pathway function, and cell surface signaling/environmental responses. | A pectin methylesterase inhibitor (Migut.C00970) that controls cell wall rigidity. A pectin lyase protein (Migut.M00012) that breaks down pectin in cell walls. EHD1 (Migut.H01794), a gene that regulates endocytic recycling. Two arabinogalactan proteins (AGP20 (Migut.F00245), AGP12 (Migut.K00009)), which are cell surface glycoproteins involved in signaling. |
| Green | Involved in balancing protein quality control and programmed cell death regulation. May be involved in acting as a molecular switch to determine whether stressed cells recover or undergo apoptosis. | Metacaspase 9 (Migut.K00195), a key protease that executes programmed cell death. A defender against death (DAD) protein (Migut.A01174). Galactinol synthase 2 (Migut.B00857), a gene involved in producing osmoprotectants to abiotic stress. A MADS-box transcription factor (Migut.K00994) that may play a key role in orchestrating stress responses and/or developmental impacts of the module. |
| Black | Ribosome biogenesis, RNA processing, and protein folding, likely serving as a hub for protein production and cellular growth. | Seven genes with ribosomal structure and function (Ribosomal L38e (Migut.F01852), S19 (Migut.G00718), S12/S23 (Migut.J00403), L35Ae (Migut.L01988), S8e (Migut.M01688), L14p/L23e (Migut.N02225), ans L1p/L10e (Migut.D02448). A peptidyl-prolyl isomerase (Migut.B00563) involved in protein folding. A cell cycle controlling cyclin (Migut.L00181). |
| Cyan | Enriched for microtubule processes, cell cycle regulation, chromosome dynamics, genome stability, and cytokinesis. Likely involved in plant growth and development through mitotic cell division and cell differentiation. | Microtubule and kinesin motor proteins (Migut.H01772, Migut.F00327). TPX2 (Migut.M00163), which is essential for spindle assembly. A cyclin-dependent kinase (Migut.C00348). Histone H2A6 (Migut.D02117) and a DNA methyltransferase (CMT3, Migut.D02536). |
| Lightcyan | Likely plays a central role in rapid responses to biotic stresses through pathogen/herbivore resistance and oxidative stress management. | Two MLP-like defense proteins (MLP423, Migut.N00310, Migut.N00312) involved in pathogen recognition/response. A serine protease inhibitor (Migut.A00978) involved in protection against herbivore and pathogen proteases. A peroxidase superfamily protein (Migut.J00066). An auxin responsive protein (Migut.H01293). |
| Blue | Chloroplast function and photosynthesis, including with both photosystems (I and II), light harvesting complexes, and thylakoid membrane components. The module also is involved in protein folding and redox homeostasis. | A PsbP-like protein 1 (PPL1, Migut.B00505), which is a component of photosystem II. A chloroplast stem-loop binding protein (CSP41A, Migut.D00296). DnaJ/Hsp40 molecular chaperone (Migut.K00479). Ascorbate peroxidase 4 (APX4; Migut.C00083), an antioxidant enzyme. Three chloroplast ribosomal genes (RPL9, Migut.D02295.1; RPL4, Migut.N00890.1; S5, Migut.H01481). |
| Greenyellow | Auxin response and metabolism, sulfur transport, redox homeostasis, protein structure/stability, and defense compound synthesis. Likely involved in the coordination of growth and stress responses. | Amidase 1 (AMI1; Migut.F01717), which converts indole-3-acetamide to active auxin. Heat stable protein 1 (HS1; Migut.N00022), which protects proteins during heat stress. FAD/NAD(P)-binding oxidoreductase (CTF2A; Migut.A01046). MLO (Migut.G00795), a transmembrane protein involved in defense responses. |

### Gene co-expression module analyses

To identify gene co-expression modules across genotypes and developmental stages, we conducted a Weighted Gene Co-expression Network Analysis (WGCNA; Langfelder and Horvath 2008). As with the analysis of candidate pathways, we focus this analysis on the 2leaf, 4leaf, and bud developmental stages. Before network construction, we re-conducted a VST normalization with DESeq2 in R version 4.5.0. To improve the signal in the data, we filtered out low-expressed genes with fewer than 10 counts in fewer than 5 samples. Following WGCNA recommended best practices, we used a scale-free topology model fit of *R²* > 0.80. To minimize the influence of outliers, biweight midcorrelations were used instead of Pearson correlations. We used dynamic tree cutting with a minimum module size of 30 genes and deepSplit parameter of 2 to identify modules. We then merged similar modules that had correlations among module eigengenes, representing the first principal component of each module, of greater than 0.75. Module membership of genes was established through correlations between gene expression and the eigengene of each module. We defined candidate hub genes for each module as the top ten genes with the highest module membership.

To interrogate the functional significance of each identified module, we conducted a GO enrichment analysis of genes within each module using an annotation file from the *Mimulus guttatus* v2.0 genome. Here, enrichments analyses of GO terms were performed using a hypergeometric test of a comparison of lists of genes in each module to all annotated genes using a Benjamini-Hochberg FDR for multiple testing. We also closely inspected the annotations of hub genes to further understand the functional significance of each module.

## RESULTS

### Sequencing and alignment

Overall, sequencing resulted in a total of 343,561,167 raw reads. Across the 80 samples, there was a minimum of 4,271,651 reads and a maximum of 10,858,583 reads (median of 7,403,646 reads per sample). We identified 28,140 unique gene transcripts.

### Clustering analyses of gene expression

For the PCA, we retained 18,054 genes after filtering for low expression. The PCA (Fig. 1) of the prefiltered, VST-normalized expression values for these transcripts revealed a distinct grouping of samples along two major axes. Comparing gene expression through development between the coastal perennial and inland annual ecotypes of *M. guttatus* is challenging, as these ecotypes often develop at different rates and put on different numbers of leaves before flowering. Both *M. gutatus* ecotypes go through two-leaf and four-leaf vegetative stages. However, the inland ecotype often flowers at the next node after the 4-leaf stage, while the coastal ecotype transitions through 6-leaf and 8-leaf stages before forming floral buds. Despite this concern, there was a high level of clustering across all leaf stages for both the coastal perennial and the inland annual plants in the PC analysis (Fig. 1). PC1 corresponded most closely to differences between leaf and bud tissue and accounted for 53.4% of the expression variance. PC2 corresponded to the divergence between the coastal and inland ecotype samples and accounted for 19.8% of the variance. The heat map visualization results were consistent with our PCA results, and also revealed that there was no clustering of samples based on the library pool (Fig. S3).

### Differential gene expression analyses at the individual gene level

Analysis of individual gene expression using the EdgeR-voom-limma approach revealed numerous significantly differentially expressed genes for the five major contrasts derived from linear modeling (Figs. S3-S7). After filtering by the filterByExpr function, 13,920 genes were retained for statistical analyses. Of these, the following numbers of genes were significantly differentially expressed with a Benjamini-Hochberg adjusted *P*-value of <0.05: 7238 genes were differentially expressed for the main effect of genotype (Coast vs. Inland), 2398 for the main effect of developmental stage (4leaf vs. 2leaf), 10,248 for the main effect of developmental stage (Bud vs. 2leaf), 886 for the interaction between genotype and developmental stage for 4leaf vs. 2leaf, and 5337 for the interaction between genotype and developmental stage for bud vs. 2leaf. For the genotype main effect, 3417 genes had significantly higher expression in inland annual plants, while 3821 genes had significantly higher expression in coastal perennial plants. Complete gene lists with statistical results can be found in the supplementary materials. MDS visualization of the 2-leaf, 4-leaf, and bud stages for 13,920 genes retained after filtering (Fig. S9) revealed a strikingly similar ordination of samples to the PCA for all samples (Fig. 1).

### Analysis of candidate pathways

Across the nine hormone pathways, we identified 768 candidate genes using the v2.0 annotation (Table 1). Of those, 351 genes were expressed at a high enough level for individual gene analyses (above), with 195 differentially expressed between the ecotypes (genotype main effect).

There were 41 highly differentially expressed (LFC>2) hormone genes, of which 35 were in the auxin, gibberellin, or jasmonic acid hormone pathways. The jasmonic acid pathway genes in particular were overrepresented for differential expression (FDR adjusted *P* = 0.00027) and highly differential expression (FDR adjusted *P* = 2.6x10^-13^) between the coastal and inland ecotypes. While the gibberellin pathway was overrepresented for highly differentially expressed genes, the pattern was not significant after FDR adjustment.

Because of our a priori hypothesis about the role of the gibberellin and jasmonic acid pathways in divergence of the coastal and inland ecotypes, we evaluated differentially expressed genes in the context of their position in the biosynthetic and signaling pathways for these hormones (Fig. 2). Striking patterns of gene expression were observed across the core GA and JA pathway genes (Fig. 2). In the jasmonic acid pathway, all seven of the differentially expressed (genotype main effect) *JAZ* genes in the signaling pathway and all ten of the differentially expressed JA catabolism/degradation genes had higher expression in inland plants than coastal plants. In contrast, the two orthologs of the *COI1* (coronatine-insensitive protein 1) were downregulated.

**Figure 2:**
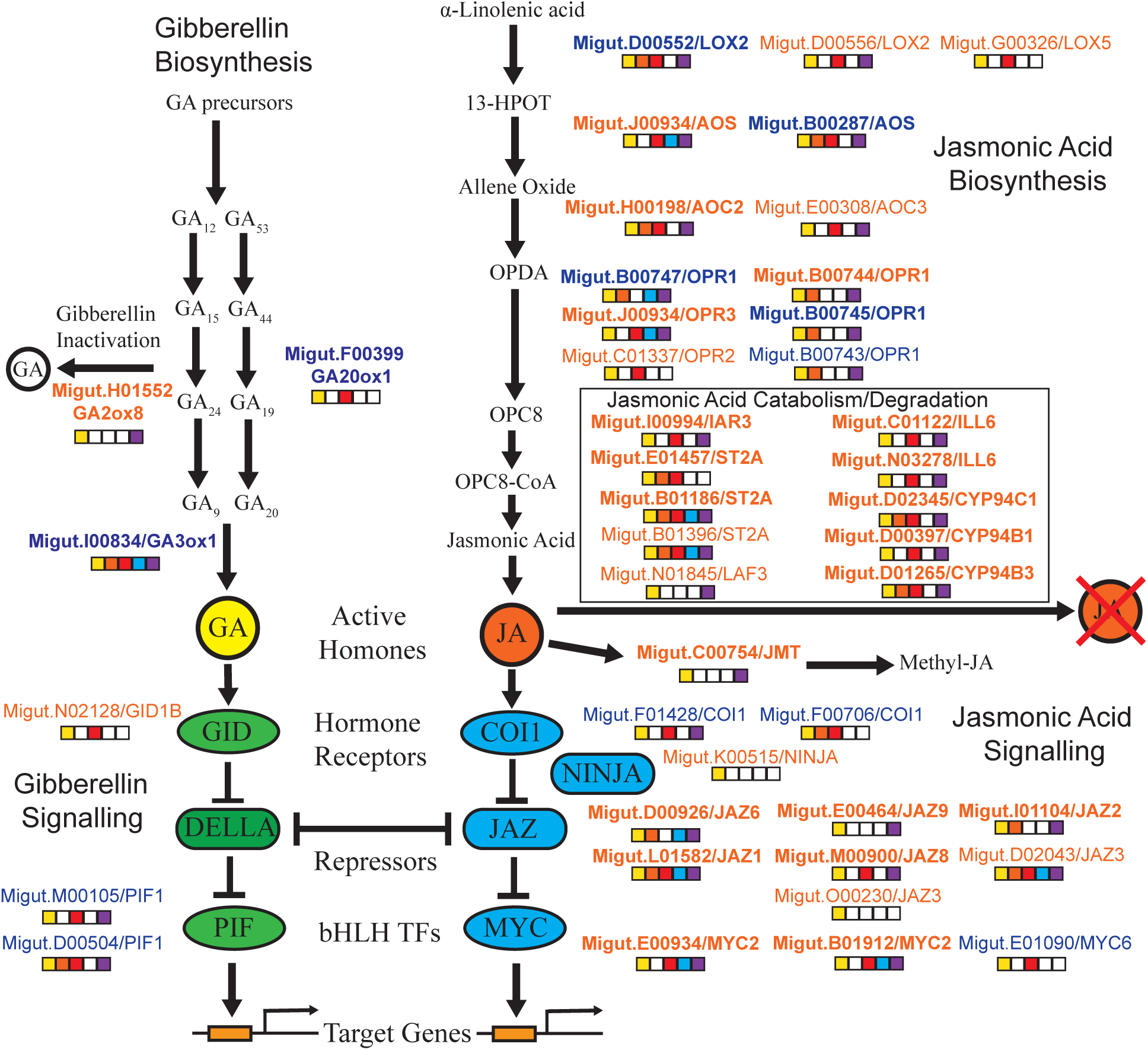
Gene expression patterns for the candidate gibberellin (GA) and jasmonic acid (JA) biosynthetic and signaling pathways. Genes that were significantly differentially expressed between coastal and inland plants (genotype main effect, yellow filled square) are shown. The names of genes with higher expression in inland plants are indicated with orange letters, while genes with higher expression in coastal plants are indicated by blue letters. Other significant contrasts are also indicated by filled squares: 4leaf vs. 2leaf (orange), bud vs. 2leaf (red), genotype interaction with 4leaf vs. 2leaf (blue), and genotype interaction with bud vs. 2leaf (purple). Bolded gene names indicate LFC > 2 for the genotype main effect.

*COI* and *JAZ* genes are co-receptors of bioactive jasmonic acid-isoleucine (JA-Ile; Monte et al. 2022). In the gibberellin pathway, genes promoting GA biosynthesis (*GA20ox1* and *GA3ox1*) were upregulated in coastal plants, while the only differentially expressed gene involved in GA inactivation (*GA2ox8*) was upregulated in inland plants. In the GA signaling pathway, an ortholog of the gibberellin receptor *GID1B* was upregulated in inland plants, while two phytochrome-interacting factor (*PIF1*) orthologs were upregulated in coastal plants. The only *DELLA* gene in *M. guttatus* (Migut.H02266) was not significantly differentially expressed between the genotypes. However, this *DELLA* gene was differentially expressed between the bud and 2-leaf developmental stages (LFC = -3.17; adjusted *P* = 8.49 x 10^-9^), with higher expression in leaves than in buds.

For each of the other hormone pathways, we report which genes were highly differentially expressed (LFC >2) here: Eight auxin genes were highly differentially expressed, including two orthologs of *ILL6* (Migut.C01122, Migut.N03278), an ortholog of *ILL3* (Migut.F00137), an ortholog of *SHY2* (Migut.N02182), and four auxin-responsive genes (Migut.F01336, Migut.F01339, Migut.F01341, and Migut.N01563). Only one brassinosteroid candidate gene was highly differentially expressed, a *BRI1* kinase inhibitor 1 (*BKI1*; Migut.N02706), which had greater expression in inland plants. The two highly differentially expressed cytokinin genes were orthologs of *ARR12* (Migut.K00347) and *APRR7* (Migut.L01650). The one highly differentiated abscisic acid gene was an *ABA4* ortholog, which had greater expression in coastal plants. The only differentially expressed salicylic acid gene, a *Mov34/MPN/PAD-1* family protein (Migut.O00742), was also highly differentially expressed. There was only one highly differentially expressed ethylene gene, an ortholog of an ethylene responsive element binding factor 1 (*ERF1*; Migut.O00367).

For genes involved in sodium homeostasis, we identified 15 high-confidence candidate genes. Of these, only five were expressed at a high enough level for individual gene statistical analyses (Fig. 3). Three of those genes were significantly expressed between the coastal and inland plants (genotype main effects): Orthologs of *SOS1* (Migut.E00570), *HKT1* (Migut.L01905), and *CIPK8* (Migut.N02833). Only *SOS1* had an LFC greater than 2 (Fig. 3, 4).

**Figure 3:**
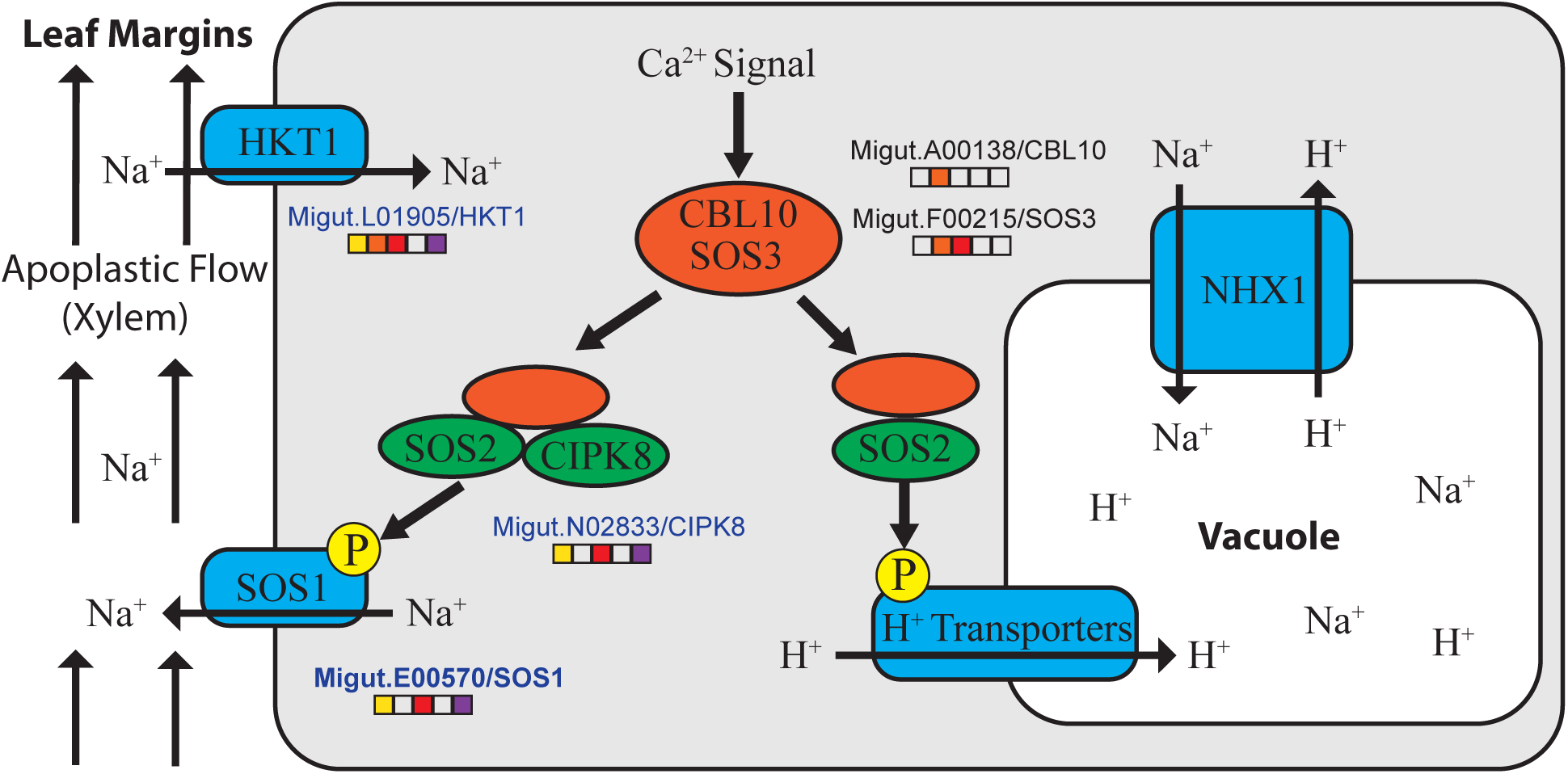
Expression of genes involved in sodium homeostasis. Model of cellular sodium homeostasis genes. *CBL10* and *SOS3* play similar roles in propagating the calcium signal resulting from salt stress, but their roles differ across cell types. Genes significantly differentially expressed between coastal and inland plants (genotype main effect) are indicated by yellow-filled squares. Names of genes with higher expression in coastal plants are indicated by blue letters. Other significant contrasts are also indicated by filled squares: 4leaf vs. 2leaf (orange), bud vs. 2leaf (red), genotype interaction with 4leaf vs. 2leaf (blue), and genotype interaction with bud vs. 2leaf (purple). Bolded gene names indicate LFC > 2 for the genotype main effect.

### Gene co-expression module analyses

Analyses with WGCNA revealed nine major gene co-expression modules after the merging of correlated modules. The following number of genes were assigned to each module: red = 2315, tan = 1874, turquoise = 4937, green =1686, black = 2633, cyan =184, lightcyan = 62, blue =3131, greenyellow = 1232. The eigengenes of each of these modules were differentially correlated with genotype and developmental stage (Figure 5; Fig. S10-S27).

**Figure 4:**
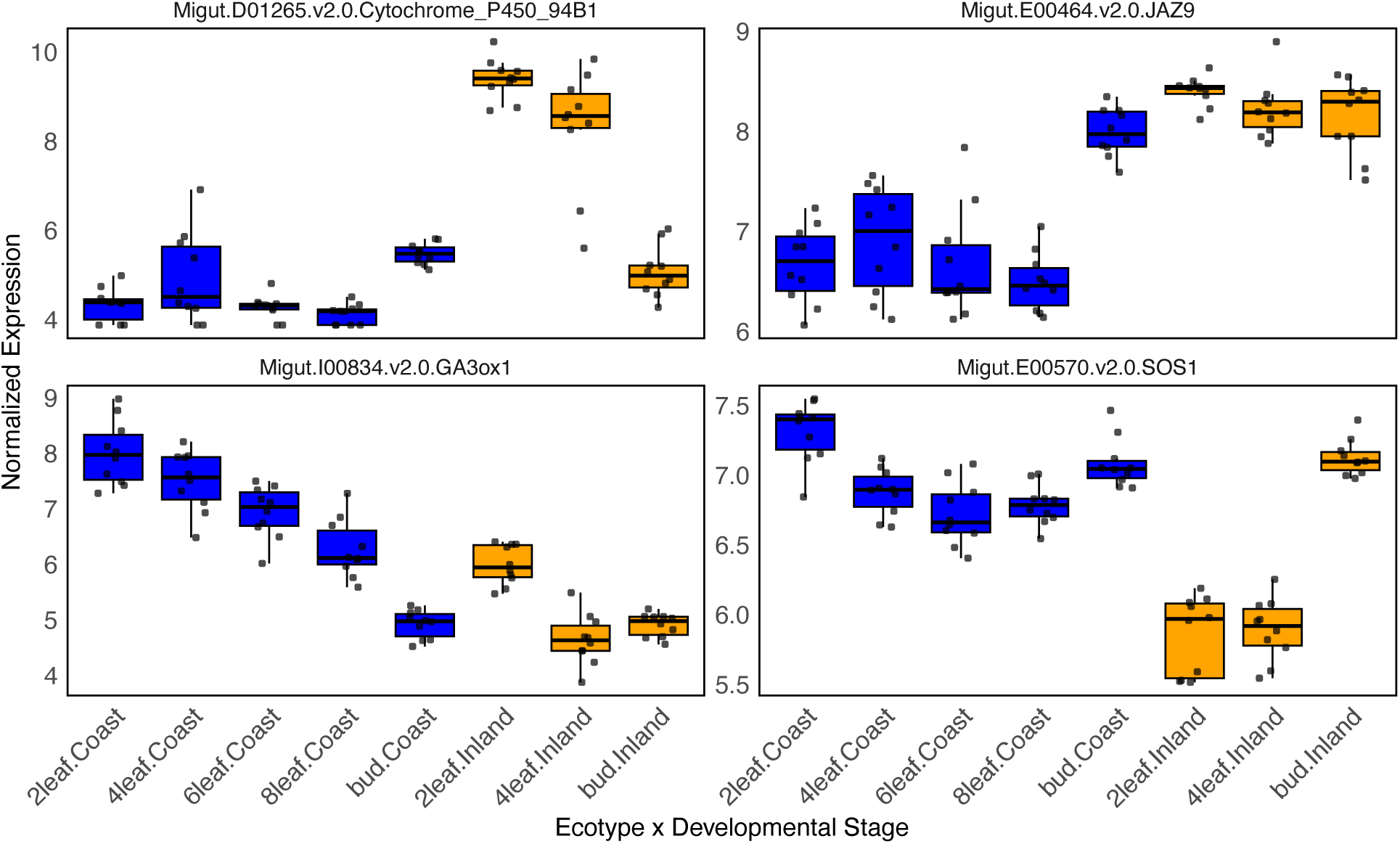
Ecotype x developmental stage interactions in gene expression. Patterns of expression for four candidate genes through developmental stages that had LFC > 2 for the contrast between the genotypes. Cytochrome P450 94B1 is the first gene in the jasmonic acid degradation pathway. JAZ9 is one of the key regulatory genes in the jasmonic acid signaling pathway. GA3ox1 is an important gibberellin biosynthesis gene. SOS1 is the key gene that pumps sodium from the cytoplasm of plant cells into the apoplast. Gene expression in the plot was VST normalized.

**Figure 5:**
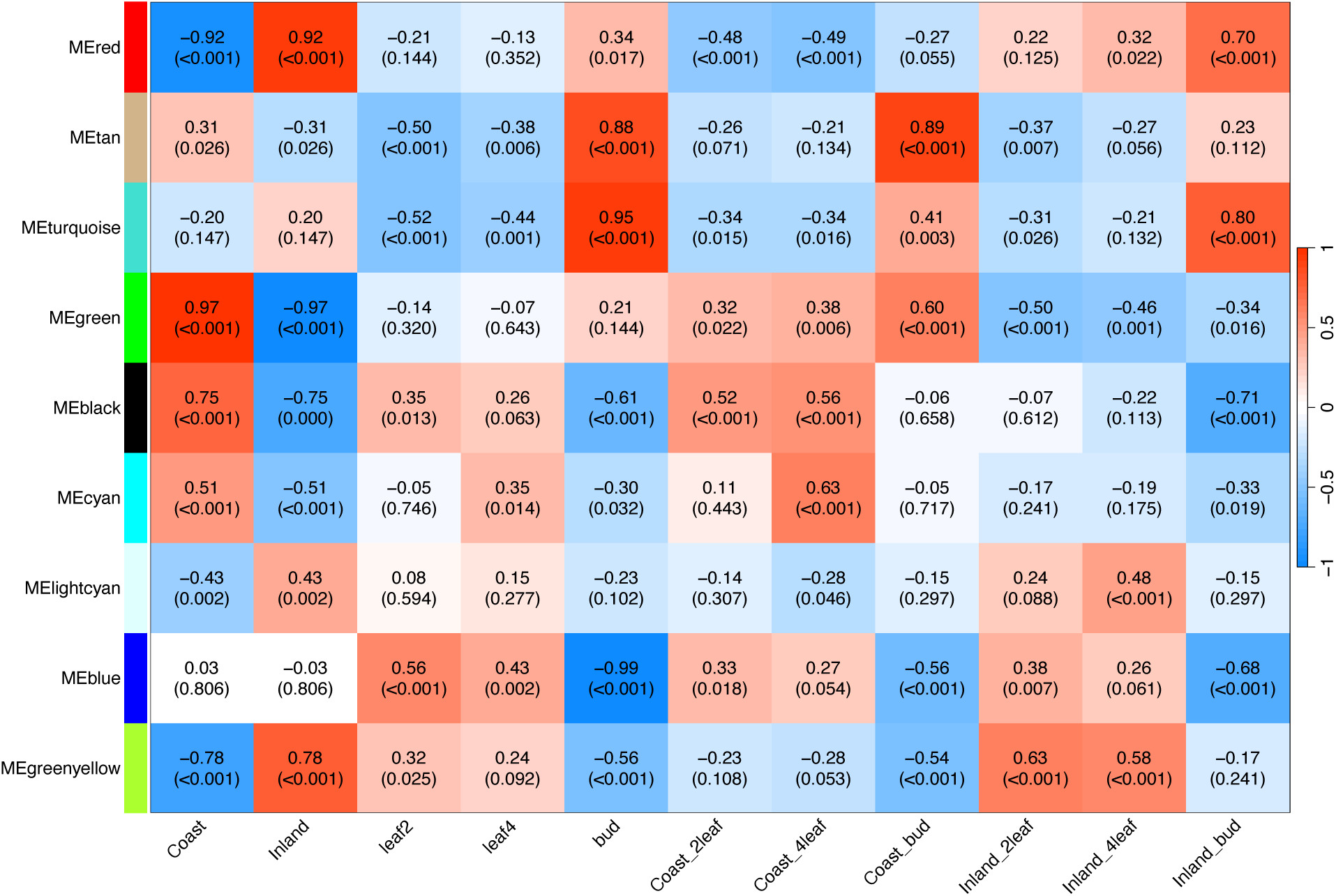
Gene co-expression module correlations with ecotype and developmental stages. The correlation between the eigengene of each module with ecotype, stage, and ecotype x stage combinations are reported above FDR adjusted *P*-values.

Gene Ontology (GO) enrichment analysis, combined with an examination of the functions of hub genes, revealed distinct potential functions for each of the nine modules. The modules that most differentiated the coastal and inland genotypes were the green, red, greenyellow, and black modules. The green module appears to differentiate coastal and inland plants in stress responses and cell death decisions. The red module appears to be primarily associated with vesicle trafficking and membrane dynamics.

The greenyellow module is likely involved in the coordination of growth and stress responses, with auxin biosynthesis and signaling potentially Genotype−Stage Combination playing a key role. One of the hub genes in the greenyellow module is a key auxin gene, *Amidase 1* (Migut.F01717).

The cyan and lightcyan modules were differentiated between the ecotypes at the leaf stages, but less differentiated at the bud stage (Fig. 5, S15, S16). Based on the gene composition of the lightcyan module, it appears to be involved in early stress responses, including plant biotic stress defense and oxidative stress management. The cyan module is likely involved in cell division and differentiation, and thus may play a key role in regulating plant growth.

Modules most associated with the developmental transition between leaf and floral bud included the blue, tan, and turquoise modules. The blue module had much greater expression in leaves than buds (Fig. 5, 6, S17) and was enriched for genes involved in photosynthesis. The tan and turquoise modules both are likely involved in the developmental transitions through changes in the expression of genes involved with the construction of cell wall and other processes.

**Figure 6:**
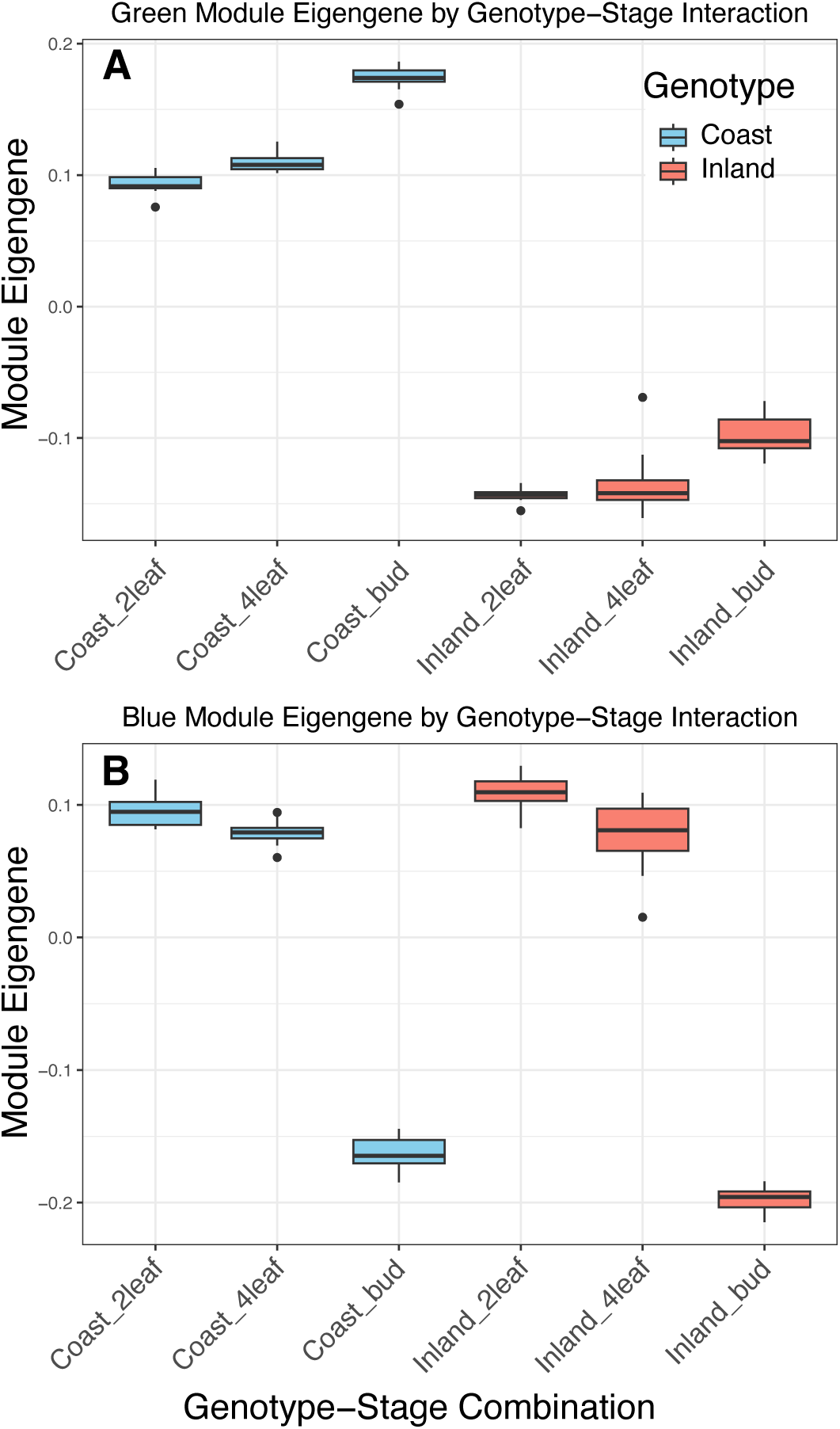
Ecotype x developmental stage of co-expression modules. (A) The green module contained genes that principally differed between the ecotypes. (B) The blue module genes were primarily differentially expressed between leaves and floral buds.

## DISCUSSION

In this study, we evaluated gene expression divergence between a coastal perennial and an inland annual accession of *M. guttatus* during development. Overall, there were clear patterns of differentiation between ecotypes and across developmental stages, with large shifts between leaf and floral bud stages. Our analysis of gene expression in candidate pathways revealed a striking elevation in differential expression between the ecotypes for genes in the jasmonic acid hormone pathway, with expression patterns in that pathway consistent with the known divergence in growth, reproduction, and defense between the coastal perennial and inland annual ecotypes.

While no other candidate pathways had significantly elevated expression differentiation, the patterns of expression for some of the key genes in those pathways were informative. In contrast to our candidate pathway analyses, the weighted gene network analyses revealed only limited evidence for a role of plant hormone pathways in the divergence of the ecotypes. Instead, these network analyses primarily revealed significant differences in cellular division, differentiation, function, and growth between the ecotypes and across various developmental stages. These differences at the microscopic level may ultimately help to give rise to organismal-level divergence of the ecotypes, but have yet to be evaluated in detail in this system. By further elucidating the mechanisms of ecotype formation, our work brings us closer to understanding the complex changes involved in the evolution of ecogeographic and phenological reproductive isolation.

### Gene regulatory divergence and the evolution of ecotypes

The finding of elevated gene expression divergence for genes in both the jasmonic acid biosynthesis and signalling pathways is a key result of our study. The higher gene expression of the full complement of genes involved in JA catabolism/degradation in inland annual plants suggests that jasmonic acid is not degraded across all stages at a high rate in the coastal perennial ecotype, while JA may be quickly degraded and recycled in the leaves of the inland ecotype. Key candidate genes, such as *CYP94B1*, exemplify this overall pattern. CYP94B1 is responsible for the first step of degradation of bioactive jasmonic acid (Koo et al. 2014) and was the single most differentially expressed gene (out of 13,920) between the ecotypes genome-wide (LFC = - 9.09; Fig. 3; Supplementary Information). The coastal ecotype had a low expression level that was consistent across all developmental stages. In contrast, the inland annual plants had a 40X higher expression at the two-leaf stage but had a similar level of expression to the coastal ecotype in the floral bud stage (Fig. 4). This pattern of gene expression suggests that jasmonic acid is not degraded at a high rate across all stages in the coastal perennial ecotype while JA may be quickly degraded and recycled in the leaves of the inland ecotype. Similarly, most of the genes in the jasomic acid signalling pathway were expressed in a pattern consistent with previous studies that found a higher production of defensive compounds in the leaves (Fig. 4) of perennial than annual *M. guttatus* populations (Holeski et al. 2013; Lowry et al. 2019; Blanchard et al. 2024).

Trade-offs between rapid growth and herbivore resistance have long been recognized by plant biologists. However, it is only recently that the underlying molecular mechanisms of such trade-offs have begun to be elucidated. The gibberellin pathway is often seen as being at odds with the jasmonic acid pathway in terms of regulating the trade-off between growth and herbivore defense (Fujita et al. 2006; Yang et al. 2012; Züst & Agrawal 2017). An important recent discovery in this area is that the gibberellin and jasmonic acid hormone pathways directly interact in an antagonistic manner though the DELLA-JAZ signaling node (Kazan & Manners 2012; Havko et al. 2016; Wasternack 2017). If this antagonistic interaction were driving trade-offs between growth and defense in *M. guttatus*, we would expect a higher level of gibberellin activity in the leaves of inland plants. This does not appear to be the case, as the pattern of gene expression in the gibberellin pathway suggests more gibberellin biosynthesis and activity of the pathway in coastal plants than inland plants. While most of the gibberellin candidate genes were not expressed at a high enough level for individual gene analyses, the candidate genes in the biosynthesis pathway all suggested a higher level of gibberellin biosynthesis, especially in leaf tissues (Fig. 2, 4)

This finding of both higher gibberellin and jasmonic acid pathway activity in coastal leaves may at first seem contradictory, except that it fits with our general understanding of how trade-offs occur in this system. Coastal plants allocate more resources to both defense and vegetative growth; thus, higher levels of expression of both the gibberellin and jasmonic acid pathways in coastal plant leaves are consistent with this model. The major trade-off in this system is with reproduction, for which the inland annual *M. guttatus* plants typically allocate more resources to reproduction over vegetative growth and resistance to insect herbivores (Hall & Willis 2006; Lowry et al. 2008; Hall et al. 2010; Baker & Diggle 2011; Baker et al. 2012; Holeski et al. 2013; Lowry et al. 2019). By allocating resources to reproduction, annual plants can successfully escape the terminal end-of-season drought that characterizes summers in western North America. In contrast, perennial *M. guttatus* plants preferentially allocate resources to vegetative growth and resistance to insect herbivores (Holeski et al. 2013; Lowry et al. 2019; Blanchard et al. 2024). The consistently lower temperatures and summer fog along the Pacific coast of North America (Gilliam 2002; Garcia-Reyes & Largier 2012) provide conditions that facilitate a perennial life history and, thus, this pattern of resource allocation.

While gibberellin biosynthesis and activity may be lower in inland plants than coastal plants at the level of leaves, this pattern is likely reversed for the region of the shoot apex, where major developmental patterns are laid down. Recent studies in this system have found that growth architecture of coastal plants undergoes greater changes after the application of bioactive gibberellin (GA3; Lowry et al. 2019). Furthermore, multiple studies suggest that the GA biosynthesis gene *GA20ox2* plays a critical role in the divergence of the coastal perennial and inland annual ecotypes (Gould et al. 2017; Kollar et al. 2025). *GA20ox2* is an allele frequency outlier between coastal and inland populations (Gould et al. 2017) and is located within the locally adaptive chromosome inversion that drives much of the divergence between the coastal and inland ecotypes (Lowry & Willis 2010; Friedman 2014; Blanchard et al. 2024). *GA20ox2* is expressed at a higher level in the shoot apices of inland plants than coastal plants (Kollar et al. 2025). We are currently conducting functional tests with this gene to determine whether it contributes to the developmental signal that drives morphological divergence between the ecotypes (Stanley & Lowry, 2024).

Beyond the jasmonic acid and gibberellin pathways, our study suggests the possible involvement of the auxin hormone pathway in the divergence of the ecotypes. While not a significant enrichment, eight of the auxin genes that we *a priori* identified were highly differentially expressed between the ecotypes. Further, auxin was implicated in three of the modules in the network analyses (tan, cyan, and greenyellow). In contrast, jasmonic acid and gibberellin were only clearly implicated in the tan module, which appears to primarily capture gene expression differences between leaf and bud stages, rather than divergence between the ecotypes (Fig. S11). With both the cyan and greenyellow modules highly differentiated between the ecotypes, the core role for auxin should be considered in future studies. Of particular interest is the ortholog of *Amidase 1* (Migut.F01717), which mediates growth and stress responses by converting indole-3-acetamide to the bioactive auxin (Pérez-Alonso et al., 2021). As a member of the greenyellow module, this *Amidase 1* ortholog may play a key role in the differences in stress responses in sulfur metabolism for coastal and inland ecotypes, as well as differences key physiological traits. Sulfur metabolism can play a critical role in plant resilience (Takahashi et al., 2023) and has previously been shown to differ in concentration between leaves of coastal and inland *M. guttatus* plants (Lowry et al., 2012).

While the evolution of prostrate coastal ecotypes is a common pattern across plant species, whether shifts in hormone pathways play an integral role in the evolution of ecotypes is still unresolved. The only other system we know where appreciable progress has been made in understanding this type of ecotype divergence is between headland and dune ecotypes of *Senecio lautus*, where ethylene, gibberelins, and especially auxin have been implicated (James et al. 2021a,b; Wilkinson et al. 2021; Broad et al. 2024). No gene in either *S. lautus* or *M. guttatus* has yet been confirmed to be involved in coastal ecotype divergence through functional genetic studies.

Our global gene network expression analysis using WGCNA yielded additional insights into the pathways that differ between ecotypes and across developmental stages. Interestingly, this network analysis alone would have resulted in a different interpretation than our candidate pathway analysis. While the gene network analyses captured new pathways to consider for divergence between the ecotypes, as well as through development, it did not capture the same insights about hormone pathway divergence. The main new insight into coastal and inland ecotype divergence from our study was the recognition that gene regulation may greatly differ in genes that control microscopic phenotypes, including vesicle trafficking and cell death regulation. These microscopic processes likely serve as a critical intermediate between the ultimately causative genetic variation at the DNA sequence level and phenotypic variation at the organismal level. Another major insight from the network analysis was that there were major differences in the regulation of photosynthetic processes between leaf and bud tissues, but not between the ecotypes, as represented by the blue module (Fig. 6b). This finding suggests that the natural variation in components of the photosynthetic machinery is likely not important for the evolution of these coastal and inland ecotypes. Natural variation in photosynthetic efficiency has been observed in other systems (Sharwood et al. 2022; Theeuwen et al. 2022), but it remains unclear whether this is due to functional allelic variation within the photosynthetic machinery itself or if it is caused by variation in genes peripheral to the core machinery.

### Salt spray adaptations of the coastal perennial ecotype

Oceanic salt spray plays a key role in the local adaptation of coastal plant populations, leading to the evolution of morphological, phenological, and physiological traits (Boyce 1954; Ahmad & Wainwright 1976; Popovic & Lowry 2020; Itoh et al. 2021, 2024). While salt spray is important for coastal population adaptations, the mechanisms underlying adaptations to this stress are poorly understood (Du & Hesp 2020). In *M. guttatus*, we have previously demonstrated that coastal perennials have higher tolerance to salt spray than inland annuals (Lowry et al. 2008, 2009; Popovic & Lowry 2020; Toll et al. 2024).

Salt spray adaptations likely share some mechanisms with soil salinity adaptations, especially in terms of leaf tissue tolerance (Munn & Tester 2008). Leaf tissue tolerance to high sodium accumulation is generally thought to involve the transport of sodium ions across cellular and vacuolar membranes to maintain cellular homeostais, which includes genes in the SOS pathway (Fig. 3; Ji et al. 2013; Yang & Guo 2018; Yin et al. 2020). The upregulation of *CIPK8* and *SOS1* in coastal plants (Fig. 3, 4) is consistent with sodium being pumped out of the cytoplasm cells into the apoplastic flow. The only other major cellular sodium homeostasis gene that was significantly differentially expressed between the ecotypes was *HKT1*, which was also upregulated in coastal plants and serves the function of importing sodium into the cytoplasm from the apoplast. These seemingly contradictory patterns of gene expression suggest that coastal plant cells are both exporting and importing sodium at higher rates than inland plants. However, this result may be explained by our RNA-seq data being quantified across entire leaves. The regulation and functions of *HKT* genes across different tissues are known to be complex (Munns & Tester 2008; Zamani Babgohari et al. 2014; Gholizadeh et al. 2024). Thus, it is possible that some coastal plant cells export sodium, while others take it up in a manner that maintains overall leaf homeostasis. With single-cell sequencing, it may be possible in subsequent studies to test the hypothesis that sodium is being actively partitioned between cells to maintain overall leaf homeostasis (Li et al. 2024).

In addition to the core cellular sodium homeostasis genes, shifts in hormone levels can confer adaptations to salt. Recent studies have found that both the jasmonic acid and gibberellin pathways can alter salt tolerance in plants, although in inconsistent ways. Context matters, as some studies have found that jasmonic acid can improve salt tolerance (reviewed in Khan et al. 2025), while others have found that it can make plants more susceptible to salt stress (Song et al. 2021). Context also appears to matter in the case of gibberellin, where the addition of this hormone generally confers higher tolerance to soil salinity (Iqbal & Ashraf 2013; Farooq et al. 2022; Liu et al. 2023), but, in contrast, makes *M. guttatus* plants more susceptible to oceanic salt spray in the field (Toll et al. 2024).

### Towards a general understanding of plant ecotype formation

Ecotypes are thought to represent an important early stage in the process of speciation (reviewed in Lowry 2012; Stankowski & Ravinet 2021; Butlin & Faria 2024). While most ecotypes will likely not go on to become new species, it is important to understand their formation as part of the speciation process, as some of them will complete the speciation process. The divergence of inland and coastal populations have served as an excellent model for understanding the speciation process, all the way back to when studies on them were used to coin the term “ecotype” in the first place (Turesson 1922a). Our study provides further evidence that the evolutionary divergence of hormone networks is a key component in the formation of locally adapted ecotypes. We hypothesize that such shifts in the complex hormone networks drive the evolution of shifts from perennality to annuality (Friedman 2020; Lundgren & Des Marais 2020) and the accumulation of both extrinsic and intrinsic reproductive isolation. Indeed, a recent study of headland and dune ecotypes of *S. latifolius* found that the evolutionary shifts in the auxin network associated with different levels of gravitropism of the ecotypes may also cause hybrid sterility (Wilkinson et al. 2021). Ultimately, functional studies of key candidate genes, currently underway, will be necessary to test the hypothesis that changes in the regulation of hormone pathways directly lead to the formation of ecotypes.

## Acknowledgments

This work was supported by grants (IOS-1855927, IOS-2153100) from the National Science Foundation and start-up funds from Michigan State University. We thank Madison Plunkert, Lauren Stanley, Joanna Feehan, Colette Berg, Daniel Ortiz-Barrientos, and two anonymous reviewers for comments that greatly helped improve this document.

## Data Accessibility and Benefit-Sharing

Raw sequence reads and processed count data are publicly available and accessible for reuse through the NCBI Gene Expression Omnibus (GEO) under accession numbers GSE280929 and GSM8607613-GSM8607692. Custom scripts for analysis can be found on GitHub at https://github.com/lowrylab/Mimulus_expression_across_development

## Author contributions

DBL and YC designed the experiment. YC conducted the experiment, performed RNA extractions, and prepared the genomic libraries. JDO, BAG, and DBL conducted the data analyses. DBL and JDO wrote the manuscript, with input from YC and BAG.

## Supporting/Supplementary information

Additional supporting information can be found online in the Supporting Information section and includes the following:

## Supplementary Methods

**Supplementary Figures S1-S27**

**Supplementary Data Files**

