## Supplementary Data Files for "Gene expression divergence between locally adapted inland annual and coastal perennial ecotypes of *Mimulus guttatus* across developmental stages": Read_Me.txt

This directory contains an annotation file for the Mimulus guttatus v2.0 genome.
The file was downloaded from Phytozome from the following webpage:
https://phytozome-next.jgi.doe.gov/info/Mguttatus_v2_0
