## Supplementary Data Files for "Gene expression divergence between locally adapted inland annual and coastal perennial ecotypes of *Mimulus guttatus* across developmental stages": Jasmonate Catabolism Genes.txt

Classification of Jasmonate Catabolism Genes in Mimulus guttatus
1. JA-Ile Hydroxylation
These genes encode cytochrome P450 enzymes that hydroxylate bioactive JA-Ile, which is the first step in its inactivation.
CYP94B1/B3 (JA-Ile 12-hydroxylases)
Convert JA-Ile to 12-hydroxy-JA-Ile

Migut.D00397.1 (CYP94B1)
Migut.D01843.1 (CYP94B1)
Migut.D01265.1 (CYP94B3)
Migut.F00069.1 (CYP94B3)

CYP94C1 (JA-Ile oxidases)
Further oxidize 12-hydroxy-JA-Ile to 12-carboxy-JA-Ile

Migut.D02345.1 (CYP94C1)
Migut.J00965.1 (CYP94C1)

Other CYP94 Family Members
Potential JA metabolizing enzymes with less characterized roles

Migut.J01622.1 (CYP94D2)

2. JA-Ile Cleavage
These genes encode amidohydrolases that cleave the JA-Ile conjugate back to JA and isoleucine.
IAR3/ILL6 (JA-Ile amidohydrolases)
Directly cleave JA-Ile conjugates

Migut.C01122.1 (ILL6)
Migut.I00994.1 (IAR3/JR3)
Migut.K00570.1 (IAR3/JR3)
Migut.N03278.1/2 (ILL6)

Other Amidohydrolases
Related enzymes that may have roles in JA-amino acid conjugate cleavage

Migut.B01364.1 (N-terminal asparagine amidohydrolase)
Migut.M02005.1 (Allantoate amidohydrolase)
Migut.N00242.1 (Ureidoglycolate amidohydrolase)
Migut.N01845.1 (Amidohydrolase family)

3. JA Methylation
These genes encode enzymes that convert JA to methyl-JA, a volatile form involved in plant-plant communication.
JMT (JA carboxyl methyltransferases)
Convert JA to methyl-JA

Migut.C00261.1 (JMT)
Migut.C00754.1 (JMT)
Migut.D01546.1 (JMT)
Migut.H02147.1 (JMT)

4. JA Conjugation with Sulfate
These genes encode sulfotransferases that may add sulfate groups to hydroxylated forms of JA.
ST2A/ST2B (Sulfotransferases)
May catalyze sulfation of hydroxylated JA derivatives

Migut.B01186.1 (ST2A)
Migut.B01350.1 (ST2A)
Migut.B01351.1 (ST2A)
Migut.B01393.1 (ST2B)
Migut.B01396.1 (ST2A)
Migut.D00840.1 (ST2A)
Migut.E01052.1 (ST2A)
Migut.E01455.1 (ST2A)
Migut.E01456.1 (ST2A)
Migut.E01457.1 (ST2A)
Migut.E01676.1 (ST2A)
Migut.G00741.1 (ST2A)
Migut.G01121.1 (ST2A)
Migut.N00726.1 (ST2A)
Migut.N00727.1 (ST2A)
Migut.N00892.1 (ST2A)

Tyrosylprotein Sulfotransferases
Different class of sulfotransferases, likely not directly involved in JA metabolism

Migut.D02233.1/2/3/4 (TPST)

Summary of Jasmonate Catabolism Genes
Jasmonate inactivation in plants occurs through multiple pathways that work together to regulate bioactive JA-Ile levels. The major routes include:

ω-Hydroxylation pathway: CYP94B1/B3 enzymes hydroxylate JA-Ile at the ω position (carbon-12), and CYP94C1 further oxidizes it to the carboxy form, both reducing hormone activity.
Cleavage pathway: IAR3/ILL6 amidohydrolases cleave JA-Ile back to free JA and isoleucine.
Methylation pathway: JMT converts JA to volatile methyl-JA, which serves in plant-plant communication.
Sulfation pathway: Sulfotransferases like ST2A may conjugate hydroxylated JA forms with sulfate groups to further metabolize them.

The presence of multiple genes for each enzyme type suggests functional redundancy and/or tissue-specific roles in JA metabolism in Mimulus guttatus.
