## Supplementary Data Files for "Gene expression divergence between locally adapted inland annual and coastal perennial ecotypes of *Mimulus guttatus* across developmental stages": Jasmonate Pathway Genes.txt

Classification of Jasmonate Pathway Genes in Mimulus guttatus
1. Biosynthesis Genes
These genes are involved in the synthesis of jasmonic acid from linolenic acid.
Lipoxygenases (LOX)
Convert linolenic acid to 13-hydroperoxy linolenic acid

Migut.D00552.1 (ATLOX2, LOX2)
Migut.D00553.1 (ATLOX2, LOX2)
Migut.D00554.1 (ATLOX2, LOX2)
Migut.D00555.1 (ATLOX2, LOX2)
Migut.D00556.1/2 (ATLOX2, LOX2)
Migut.D00557.1 (ATLOX2, LOX2)
Migut.G00326.1/2 (LOX5)
Migut.N02059.1 (ATLOX1, LOX1)
Migut.N02072.1 (ATLOX1, LOX1)

Allene Oxide Synthase (AOS)
Converts 13-hydroperoxy linolenic acid to allene oxide

Migut.B00287.1 (AOS, CYP74A, DDE2)
Migut.C00664.1 (AOS, CYP74A, DDE2)
Migut.N01221.1 (AOS, CYP74A, DDE2)
Migut.N02681.1 (AOS, CYP74A, DDE2)
Migut.O00603.1 (AOS, CYP74A, DDE2)

Allene Oxide Cyclase (AOC)
Converts allene oxide to 12-oxo-phytodienoic acid (OPDA)

Migut.E00308.1 (AOC3)
Migut.E00309.1 (AOC3)
Migut.H00198.1 (AOC2)

12-Oxophytodienoate Reductase (OPR)
Reduces OPDA to 3-oxo-2-(2'-pentenyl)-cyclopentane-1-octanoic acid

Migut.B00740.1 (ATOPR1, OPR1)
Migut.B00741.1 (ATOPR1, OPR1)
Migut.B00742.1 (ATOPR1, OPR1)
Migut.B00743.1 (ATOPR1, OPR1)
Migut.B00744.1 (ATOPR1, OPR1)
Migut.B00745.1/2 (ATOPR1, OPR1)
Migut.B00746.1 (ATOPR1, OPR1)
Migut.B00747.1 (ATOPR1, OPR1)
Migut.B00748.1 (ATOPR1, OPR1)
Migut.C01337.1/2 (ATOPR2, OPR2)
Migut.J00934.1 (DDE1, OPR3)

2. JA-Modifying Enzymes
These genes are involved in modifying JA to create active forms or derivatives.
Jasmonic Acid Carboxyl Methyltransferase (JMT)
Converts JA to methyl-JA

Migut.C00261.1 (JMT)
Migut.C00754.1 (JMT)
Migut.D01546.1 (JMT)
Migut.H02147.1 (JMT)

3. Signaling Pathway Genes
These genes are involved in jasmonate perception and signal transduction.
Jasmonate Receptor - COI1
F-box protein that forms the JA-Ile co-receptor with JAZ proteins

Migut.F00706.1 (COI1)
Migut.F01428.1 (COI1)

JAZ Repressors
Repressors of JA-responsive genes that are degraded upon JA perception

Migut.D00926.1/2/3 (JAZ6/JAZ1)
Migut.D02043.1 (JAZ3)
Migut.H01355.1/2 (JAZ8)
Migut.I01104.1/2 (JAZ2)
Migut.K01038.1/2/3 (JAZ10)
Migut.L01582.1/2 (JAZ1)
Migut.M00900.1/2 (JAZ8)
Migut.O00230.1/2 (JAZ3)
Migut.E00464.1 (JAZ9)

JA Signaling Regulators
Proteins that regulate JA signaling pathway

Migut.K00515.1 (NINJA) - Negative regulator of JA responses

Transcription Factors
Transcription factors that regulate JA-responsive gene expression

Migut.B01912.1 (MYC2) - Master regulator of JA responses
Migut.E00934.1 (MYC2) - Master regulator of JA responses
Migut.A00733.1 (GL3/MYC6.2) - bHLH transcription factor
Migut.E01090.1/2/3 (GL3/MYC6.2) - bHLH transcription factor
Migut.H00411.1 (DEAR1) - ERF transcription factor responsive to ethylene and jasmonate
Migut.L00172.1 (DEAR1) - ERF transcription factor responsive to ethylene and jasmonate

Summary Counts

Biosynthesis genes: 34 (9 LOX, 5 AOS, 3 AOC, 17 OPR)
JA-modifying enzymes: 4 (all JMT)
Signaling pathway genes: 25 (2 COI1 receptors, 15 JAZ proteins, 1 NINJA regulator, 7 transcription factors)
Total genes: 63
