## Supplementary Data Files for "Gene expression divergence between locally adapted inland annual and coastal perennial ecotypes of *Mimulus guttatus* across developmental stages": Read_Me.txt

This directory contains sub-directories for each of the candidate pathway analyses. 

In each subdirectory, there is a list of candidate genes, an annotation file for those
genes, and the results from the voom-limma contrasts for each of the genes expressed
at a high enough level for individual gene analyses. 
