## Supplementary Data Files for "Gene expression divergence between locally adapted inland annual and coastal perennial ecotypes of *Mimulus guttatus* across developmental stages": Read_Me.txt

This directory includes the two files from which all analyses of gene expression were 
conducted for this study. 

All_Counts_R_Input.txt contains the raw count data for all genes for the 80 samples.

All_Counts_Sample_Metadata.csv contains the meta-data for the 80 samples.

The two files are connected by the sample numbers 1-80.
