## Supplementary Data Files for "Gene expression divergence between locally adapted inland annual and coastal perennial ecotypes of *Mimulus guttatus* across developmental stages": Read_Me.txt

This directory contains the results of the individual gene analysis using the voom-limma 
approach. 

Overall_Voom_limma_results.txt contains the overall results for each major contrast of the 
model.

voom_all_genes_genotype_effect.csv contains the results for each gene for the genotype main 
effect (coast vs inland).

voom_all_genes_4leaf_vs_2leaf_effect.csv contains the results for each gene for the contrast 
in gene expression between the fourth (4leaf) and second set of true leaves (2leaf).

voom_all_genes_bud_vs_2leaf_effect.csv contains the results for each gene for the contrast 
in gene expression between the floral bud and second set of true leaves (2leaf).
 
voom_all_genes_interaction_4leaf_effect.csv contains the results for each gene for the 
genotype interaction with the contrast in gene expression between the fourth (4leaf) and 
second set of true leaves (2leaf).

voom_all_genes_interaction_bud_effect.csv contains the results for each gene for the 
genotype interaction with the contrast in gene expression between the floral bud and 
second set of true leaves (2leaf).
