## Supplementary Data Files for "Gene expression divergence between locally adapted inland annual and coastal perennial ecotypes of *Mimulus guttatus* across developmental stages": Read_Metxt.txt

This directory contains the results of the WGCNA analyses and subsequent GO enrichment 
analyses of each discovered module of gene co-expression.

eigengenes_by_sample_and_module.csv contains the eigengene value for each sample (1-20,41-80)
in each of the nine modules. The sample number corresponds to the sample number in the 
raw count input files found in the Raw_Count_Input_Files directory.

The subdirectories contain the lists of genes in each module and the enrichment analyses. 
