## Supplementary Methods for "Gene expression divergence between locally adapted inland annual and coastal perennial ecotypes of *Mimulus guttatus* across developmental stages"

### RNA-Seq library sample preparation for sequencing on the Illumina HiSeq – v6

Following the protocol developed by Meyer & Matz RNA-Seq sample preparation for sequencing on the SOLiD System (Meyer, Aglyamova et al. 2011).

**Important points before starting**

This protocol describes in detail the procedures used to prepare cDNA fragment libraries for quantitative analysis of gene expression (RNA-Seq) by deep sequencing on the Illumina HiSeq System.

About 1 µg DNAse-treated total RNA is required per sample, and this starting material should be carefully quantified and analyzed by gel electrophoresis prior to beginning these procedures to verify that the RNA is intact, and free of genomic DNA contamination. RNA concentration is measured by Nanodrop, DNA concentration is measured by Qubit(RTSF at the basement of PLB).

The procedure can be reasonably completed within three days.

Day 1: RNA is fragmented and used for cDNA synthesis (steps 1-2).

Day 2: cDNA is amplified, sample-specific barcodes are incorporated, and size-selection is accomplished by means of gel extraction (steps 3-4).

Day 3: The quality of preparations is evaluated by PCR and DNA quantified by qPCR (the qPCR step is performed by staffs at the RTSF).

**Procedure**

1. RNA fragmentation
   a. Aliquot 1 µg of total 100 ng/ul RNA in 10 µl of super-clean water.
   b. Incubate RNA in PCR tubes at 95°C (using thermocycler) for **12** minutes to fragment to the desired size range (350 – 600 bp) (See Figure 1).
   c. Analyze 100 ng from fragmented RNA, alongside the intact RNA from the same sample, on a standard 2% agarose gel to evaluate the extent of RNA fragmentation. The smear must extend all the way up into the region where ribosomal RNA bands were, while the bands themselves should be mostly gone. No need to run every sample on the gel, can just run a few to check for proper degradation.

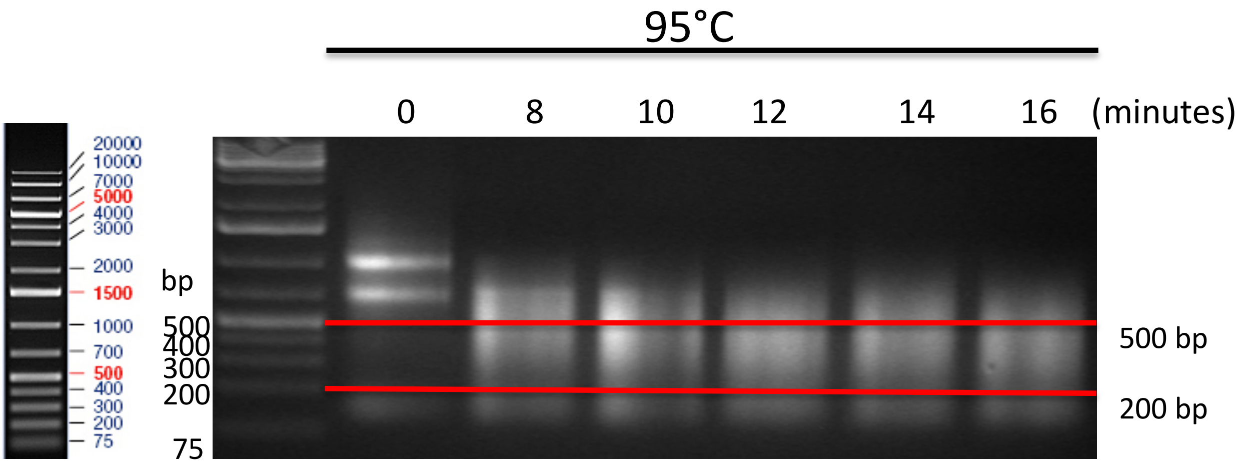


Figure 1 RNA fragmentation

2. <u>First-strand cDNA synthesis</u>

*NOTE: if RNA quantity is not limiting, first-strand cDNA should be synthesized using 1 µg of fragmented RNA to ensure adequate representation of all transcripts. The reaction shown below is intended for ~ 1 µg; this can be doubled or halved as needed if the amount of RNA is very different from 1 µg.*

a. Measure the volume remaining after fragmentation using a pipette. The following recipe assumes a starting volume of 9 µl (10 µl minus evaporation or loss of 1 ul that was run on the gel in step1), so if the volume is lower than this add additional water to achieve 9 µl.

b. Add 1 µl of the 10 µM oligonucleotide 3ILL-30TV to each well. Incubate at 65°C for 3 minutes in a thermocycler, then transfer immediately onto ice.

c. Prepare a cDNA synthesis master mix. The following volumes are intended for a single reaction, so multiply these values by the number of reactions plus a small amount (~10%) to account for pipetting error.

(Volumes given in µl)

| | |
|---|---|
| $H_2O$ | 1 |
| dNTP (10 mM each) | 1 |
| DTT (0.1 M) | 2 |
| 5X first-strand buffer | 4 |
| 10 µM S-ILL-swMW (RNA oligonucleotide; stored at -80°C) | 1 |
| SuperScript II Reverse Transcriptase (Invitrogen #18064-022) | 1 |
| Total volume | 10 |

d. Add 10 µl of this master mix to the RNA from (2b), mix thoroughly, and incubate for one hour at 42°C. Next incubate at 65°C for 15 minutes to inactivate the RT. Accomplish it in a thermocycler. **<u>Do NOT dilute</u>**! Store on ice or at -20°C until ready to proceed to the next step, where you will use 12 ul from this reaction.

3. <u>cDNA amplification</u>

.

(Volumes given in µl)

| Reagent | A | B |
|---|---|---|
| $H_2O$ | 13.2 | 12.8 |
| dNTP (2.5 mM ea) | 2 | 2 |
| 10X PCR buffer | 2 | 2 |
| 10 µM 5ILL oligo | 0 | 0.4 |
| 10 µM 3ILL-30TV oligo | 0.4 | 0.4 |
| Titanium Taq polymerase (Clontech # 639208) | 0.4 | 0.4 |
| FS-cDNA template from step 2e | 2 | 2 |
| Total volume | 20 | 20 |

a. Amplify in a thermocycler using the following profile:

95°C 5 min, (95°C 40 sec, 63°C 1 min, 72°C 1 min) X **<u>18</u>** cycles

b. Check 5-10 µl of the PCR products for all reactions on a standard 2 % agarose gel. A "smear" of cDNA (~200-500 bp) should be faintly visible in reaction B (Figure 2), and nothing or reduced smear intensity/profile should be detected in the reactions A.

c. prepare a single large-scale reaction for each cDNA sample as follows. This recipe assumes 12 µl of <u>undiluted</u> template, so if you use more template adjust the water accordingly.

| | (All volumes given in µl) |
|---|---|
| $H_2O$ | 25 |
| dNTP (2.5 mM ea) | 5 |
| 10X PCR buffer | 5 |
| 10 µM 5ILL oligo | 1 |
| 10 µM 3ILL-30TV oligo | 1 |
| Titanium Taq polymerase | 1 |
| FS-cDNA template from step 2e | 12 |
| Total volume | 50 |

d. Amplify in a thermocycler using the following profile:

95°C 5 min, (95°C 40 sec, 63°C 1 min, 72°C 1 min) X **18** cycles (for *Mimulus*)

e. After PCR, check 5 µl of the product on a 2 % agarose gel to verify that the reaction worked as expected <u>before</u> freezing or purifying the product.

f. Purify PCR products using Macherey-Nagel NucleoFast PCR Clean-up protocol (Cat. No. 743500.4) according to the manufacturer's instructions. Exact time of each step varies, but can be easily monitored for column drying:

  a. Transfer PCR samples (~45 ul each) to NucleoFast 96 PCR plate.
  b. Filter contaminants to waste under vacuum for 12 min.
  c. Wash membrane with 100 ul of RNase-free water under vacuum for 12 min.
  d. Recover purified PCR samples add 30 ul of RB, incubate for **30** minutes at room temperature after the addition.
  e. Quantify the purified products by Qubit HS. Ideally, your concentration will be 5 ng/ul. But very often it is less, so you will adjust your volumes accordingly in the next step. If concentration is below 1.5 ng/ul, you do not have enough DNA. Repeat PCR with more template or more cycles.

g. Prepare 30 µl of the purified PCR products diluted to 5 ng $\mu l^{-1}$ (in the RB elution buffer from the PCR-cleanup kit).

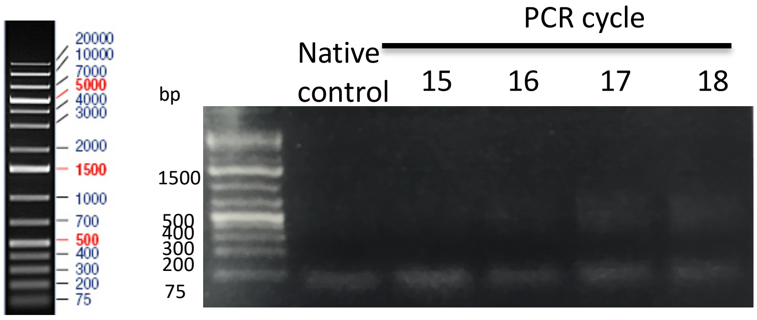


Figure 2. cDNA PCR test

4. <u>Adaptor extension and size selection</u>

*NOTE: Because the size distribution of templates is a critical factor for successful emulsion PCR, any templates intended for sequencing on the Illumina System should be carefully size-selected prior to emPCR. The directions below outline a simple procedure for selecting fragments ranging from 350-500 bp in size that does not require any special equipment. Other methods of size selection could be substituted provided they achieve this same size range.*

a. First, two test-scale PCRs are prepared for a representative subset (6-8) of samples to verify yield and specificity of the reaction, each using 10 ng (2 µl) PCR product (step 3h above) as template.

b. Prepare two separate master mixes for small-scale test PCR. The following volumes are for a single reaction, so multiply these values by the total number of samples plus a small additional amount to account for pipetting error. The values shown here assume the use 10 ng (2 µl from step 3h above) clean PCR product as template, so if you change this be sure to change the volume of water accordingly. Be sure to write down which barcode and multiplex is assigned to each sample at this stage, since this cannot be easily determined later in the process.

| | A | B |
|---|---|---|
| H2O | 5.8 | 5.4 |
| dNTP (2.5 mM ea) | 1 | 1 |
| 10X PCR buffer | 1 | 1 |
| Multiplex oligo (10 µM) | 0 | 0.2 |
| Barcode oligo (10 µM) | 0 | 0.2 |
| Titanium Taq polymerase | 0.2 | 0.2 |
| cDNA template from step 3h | 2 | 2 |
| Total volume | 10 | 10 |

c. Amplify in a thermocycler using the following profile:

95°C 5 min, (95°C 40 sec, 63°C 1 min, 72°C 1 min) X **7** cycles

d. After 4 cycles check 5 µl of the PCR products for all reactions on a gel (For rice 6 cycles are OK). The ideal result is a faint smear in reaction B, with no visible product in reaction A (Figure 3). However, reaction A may has some smear, but should be considerably less, sometimes with changed profile. If nothing is detected in reaction B, add 1-2 more cycles and check the results on a gel. If no product is visible before 6 cycles, repeat the reaction with a larger volume of template (in our experience this has never been required). A small amount of product in the controls can be tolerated, but if reaction A are comparable in intensity to reaction b something is wrong.

e. When the optimum number of cycles and volume of template have been determined, prepare a large-scale reaction based on those parameters with 50 ng (10 µl of the diluted purified cDNA, step 3h) template in 50 µl total volume. The following master mix assumes the use of 10 µl of template per 50 µl reaction; if you adjust this template volume be sure to adjust the volume of water accordingly. This recipe is for a single reaction, so multiple these values by the number of samples to be prepared plus a small additional amount for pipetting error.

(Volumes given in µl)

| | |
|---|---|
| $H_2O$ | 27 |
| dNTP (2.5 mM ea) | 5 |
| 10X PCR buffer | 5 |
| Multiplex oligo (10 µM) | 1 |
| Titanium Taq polymerase | 1 |
| PCR template from step 3h | 10 |
| Barcode oligo (10 µM) | 1 |
| Total volume | 50 |

f. Amplify these reactions using the same profile and cycle number as determined above.

95°C 5 min, (95°C 40 sec, 63°C 1 min, 72°C 1 min) X **7** cycles

g. Run a small amount (5 ul) on 2% gel to make sure you have amplified your DNA well. This step can be omitted by seasoned users. Or only a select number of samples can be checked.

h. Combine all 32 reactions in a single tube. We estimate that 32 different barcoded reactions can be mixed together to still yield 5 million reads per sample. Thus, all 32 reactions can be combined at this step, and this combined DNA purified through NucleoFast 96 PCR plate. 50x32=1,600 ul, so it will take several to many hours for the whole sample to go through the purification step. You can split that sample over 2-4 wells in the plate if time is an issue, or keep it in a single well. Upon elution from the plate combine your fractions together again to obtain one pool of 32 samples.

i. Prepare an agarose gel for size selection. This preparative gel should consist of 1.5% agarose. A low-molecular weight ladder is required for accurate selection of the appropriate sizes; we recommend 100 bp ladder (Invitrogen, #15628-019), or DNA MW marker 100 bp (amresco, #n550-300ul) or Low MW Ladder (NEB, #N3233S) or something similar. Be sure to use large volume combs (tape them if necessary) to allow loading of the entire 35-100 µl reaction (depending on the number of wells used in the previous step) and loading dye into a single well.

j. Important. Run the gel at low voltage (not more than 80 V). Also important is 1.5%, not 2% agarose.

k. Load samples and run the gel until marker bands in the 100-500 bp size ranges are well separated. Illuminate the gel very briefly (< 30 seconds total exposure time) on a UV-transilluminator set at low intensity, for just long enough to mark the appropriate region (300-500 bp) with a clean razor blade. Cut only the middle of the lane; leave the edges (see picture). Turn off the UV light and carefully cut out the marked region, transferring it into a microcentrifuge tube (1.7-2 ml tubes).

l. Extract the cDNA from this gel slice using the Invitrogen's PureLink Quick Gel extraction and PCR purification combo kit (#K220001), following manufacturer's suggestion. No further purification procedures are necessary.

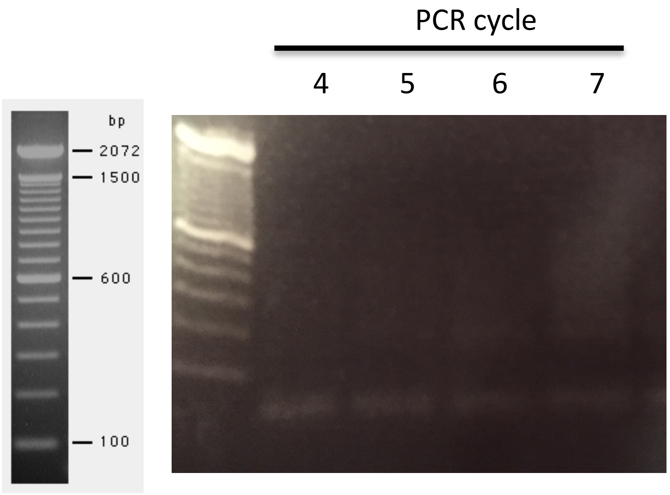


Invitrogen ladder (15628-019)

Figure 3. Adaptors extension

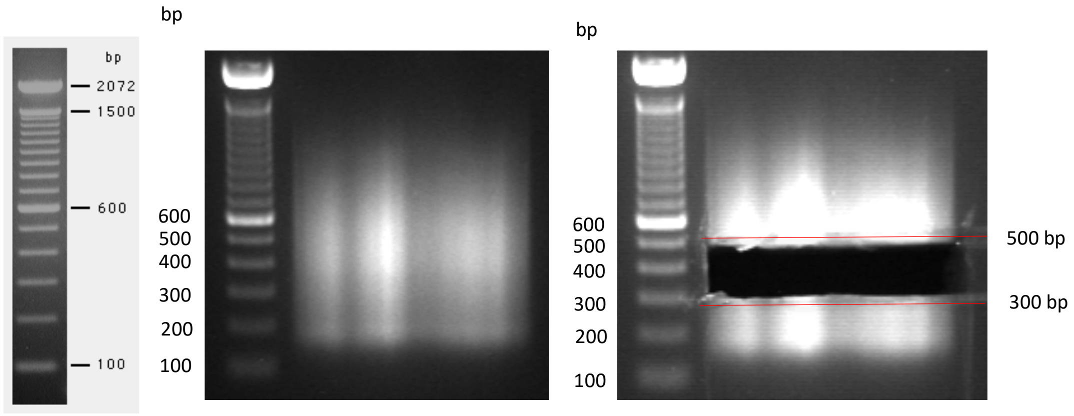


Invitrogen ladder (15628-019)

Figure 4. Size selection

5. Quality check

*NOTE: Illumina preps other than RNAseq can be similarly processed using the same primers and scripts; quantify all the preps that are to be mixed on the same lane simultaneously. For checking quality and quantity of gel eluted DNA we do two PCR – one –to check the product size on gel – it should be the same as the band we cut out – no additional products. For mixing samples together in equal proportions we perform qPCR with P5 and P7 primers and mix samples according to Ct analysis.*

a. For quality check prepare a PCR master mix. The following volumes are for a single reaction, so multiply these values by the total number of reactions plus a small additional amount to account for pipetting error.

(Volumes given in µl)

| | |
|---|---|
| $H_2O$ | 6.4 |

| | |
|---|---|
| dNTP (2.5 mM ea) | 1 |
| 10X PCR buffer | 1 |
| IC-P7 primer (10 µM) | 0.2 |
| IC-P5 primer (10 µM) | 0.2 |
| Titanium Taq polymerase | 0.2 |
| DNA template from step 4l | 1 |
| Total volume | 10 |

b. Amplify in a PCR-thermocycler using the following profile:

95°C 5 min, (95°C 40 sec, 63°C 1 min, 72°C 1 min) X 12 cycles

Run 5 (or less) µl on gel. The size of the product should match the size you aiming when cut a band for gel-extraction.

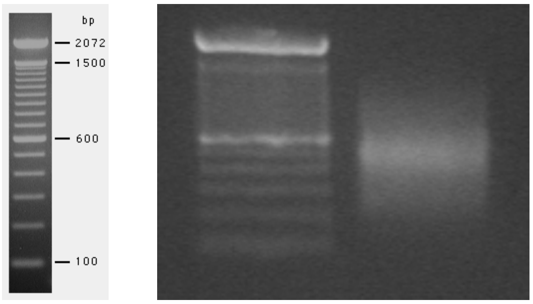


Invitrogen ladder (15628-019)

Figure 5: Quality test

**For routine things you can stop here…**

6. qPCR quantification for mixing on the same HiSeq lane

c. This step is only necessary when you are just optimizing your protocol. For qPCR quantification, for mixing samples in equal proportions into the library prepare two dilutions of each sample (clear elute from step 4l), 1/10 and 1/50 in 10 mM Tris-HCl. Arrange the dilutions in a 96-well plate for easy pipetting. For each sample, make two identical dilution series – these will be your technical replicates. Do not simply split the same dilution series into two – we aim to replicate the process of making those dilutions.

d. Mix SYBR-based qPCR Master mix appropriate for your qPCR instrument with water and two primers, according to the recipe below, and aliquot 14 µl per reaction. Plan for two no-template-control (NTC) reactions.

(Volumes given in µl)

| | |
|---|---|
| $H_2O$ | 6.1 |
| SYBR Green mix | 7.5 |
| IC-P7 primer (10 µM) | 0.2 |
| antiT30 primer (10 µM) | 0.2 |
| Dilution template from step 5c | 1 |
| Total volume | 15 |

m. Amplify in a qPCR machine with some NTC (no-template control) using the following profile:

95°C 5 min, (95°C 40 sec, 63°C 1 min, 72°C 1 min) X 20 cycles

(The product should amplify around 8-12 cycles; the NTC controls should be empty).

n. Arrange the data in Excel in the form of a table with four columns: sam (sample name), lane (intended HiSeq lane), conc (DNA dilution; use 0.1 for 1/10, 0.02 for 1/50), and ct (qPCR result for this sam-conc combination). There must be at least two technical replicates for each combination of sam-conc (i.e. two rows with the same sam and conc and different ct values). If all samples are to be mixed on the same lane, enter '1' throughout the lane column. The order of columns and rows does not matter, but the names of the columns do matter (they are case sensitive).

o. Export the data from Excel as comma-separated values (.csv). Open script mix_illumina_qpcr.R in R, follow the instructions given in the comments within the script.

p. Mix samples (elutes from the gel slices, step 3m) according to final mixing table produced by the script. This material is in principle ready for Illumina sequencing, except you might need to concentrate the resulting sample 2-3 fold to meet the sequencing facility requirements. In that case, we recommend mixing a larger volume of the all-barcodes mixture and concentrating it using SpeedVac.

**Sequences 5' - 3'of oligonucleotide primers used in this protocol**

3ILL-30TV ACGTGTGCTCTTCCGATCTAATTTTTTTTTTTTTTTTTTTTTTTTTTTTTTV

5ILL CTA CAC GAC GCT CTT CCG ATC T

S-Ill-swMW ACCCCATGGGGCTACACGACGCTCTTCCGATCTNNMWGGG

(*Note: This custom RNA oligo should be stored in multiple aliquots at -80°C to prevent degradation of this labile and expensive reagent*).

TruSeq-Mpx-2n (similar to the Illumina_Universal)(56 pb length)
AATGATACGGCGACCACCGAAAAATACACTCTTTCCCTACACGACGCTCTTCCGAT

ILL -BC (BC=*NNNNNN*)(62 pb length)
CAAGCAGAAGACGGCATACGAGAT*NNNNNN*GTGACTGGAGTTCAGACGTGTGCTCTTCCGAT

**BC(*NNNNNN*)**

| | | | |
|---|---|---|---|
| AATGCT | TCTATA | ACTTGA | TGAAGG |
| GACACA | TGCAAA | CCGTCC | GGTGTG |
| GAGTGG | TGGCAC | TAATCG | AGCGAG |
| CACCGG | TGTTAG | TATAAT | AGCTTT |
| GAAGTT | AAGGGA | TCATTC | TGGTCT |
| GCAGGA | GCACCC | TCCCGA | ACCGGC |
| GTATTA | TACGTG | TCGAAG | AAAAGT |
| TCACAT | TTAGGC | AAACAC | ACATCT |

Total adaptors length 118 pb

The barcoding oligos given in the table are a selection of standard Illumina TrueSeq barcodes that have The best "Hamming distance" barcode to complete a line (32 samples*5 million of reads each = 160 millions of reads); more barcode sequences can be found elsewhere.

If ordering from IDT, order them as "ultrameres" with no purification; this seems to be the best quality-cost balance. Remember that the barcode will be read in a reverse-complement orientation compared to the sequence in the table.

IC-P5 AATGATACGGCGACCACCGA

IC-P7 CAAGCAGAAGACGGCATACGA

antiT30 AAATTAGATCGGAAGAGCACAC

#### Figures

Figure 1. Degradation time (Step 1b). 0 in wells 1 and 2 mean samples without degradation. 8 mean 8 minutes of degradation, and so far.

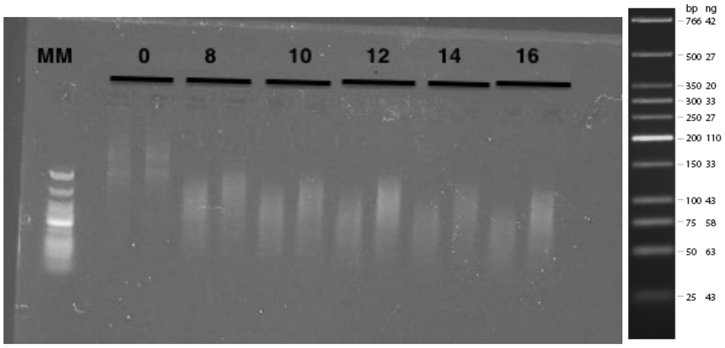


MM: Low molecular weight DNA ladder, New England BioLabs, Catalogue number N3233S. 2 % agarose gel, 100 ng RNA, 20 min 115 V.

Figure 2. cDNA PCR test

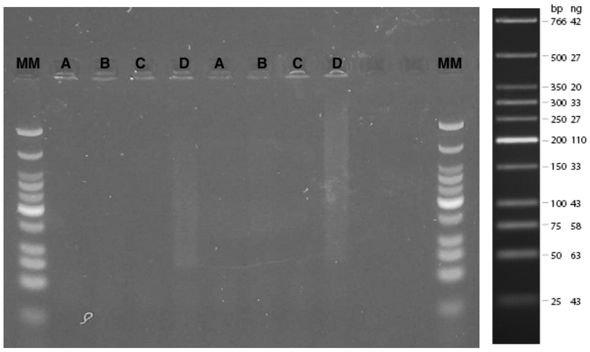


2% agarose, 10 ul PCR product loading, 35 minutes, 100 V. MM: Low molecular weight DNA ladder, New England BioLabs, Catalogue number N3233S. 2 % agarose gel, 100 ng RNA, 20 min 115 V.

Figure 3. Adaptors extension

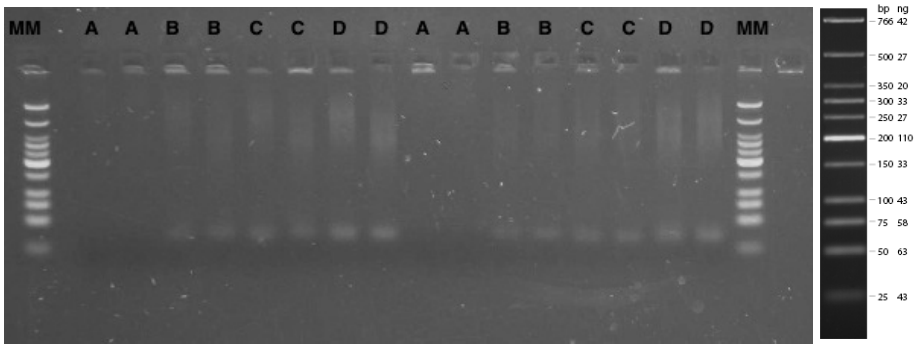


2 % agarose gel, 10 ul product, 30 minutes at 100 V.

Figure 4. Size selection

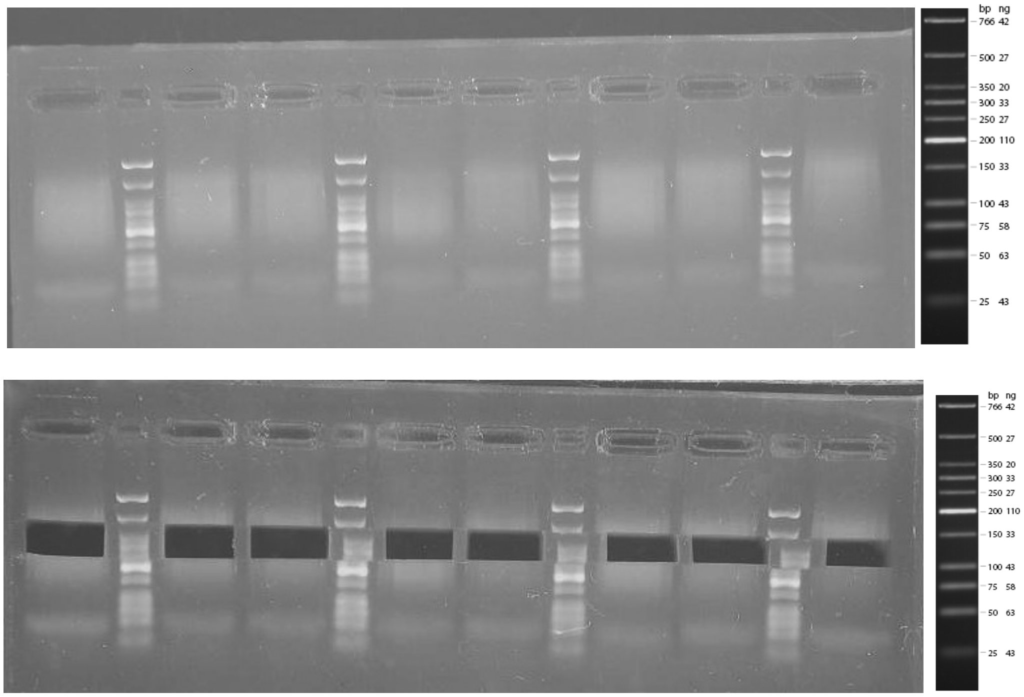


2 % agarose gel, 40 ul product, 45 minutes at 130 V.

Figure 5: Quality test

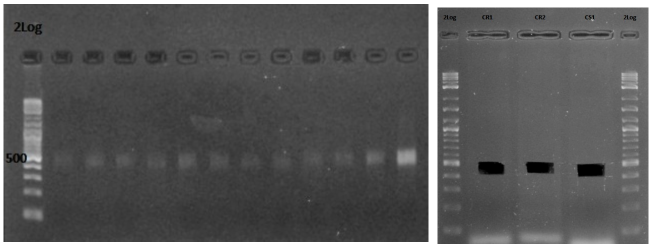

**Submission process to Genomic Sequencing and Analysis Facility (GSAF):**

1. Go to the GSAF homepage

   https://wikis.utexas.edu/display/GSAF/Illumina+-+all+flavors

   Go to “how to submit samples to GSAF” and download the Excel spreadsheet “Sample_Submission_Template.xls”

2. Fill the Excel spreadsheet file. Take care to:
   1. Use just two decimal numbers in the concentration column
   2. Save the file with .xls extension (Excel 97-2004 wordbook).

3. In the GSAF homepage go to “submit Samples to the GSAF” and them open “online project submission form”
   - In the field “brief job description” put:
     “3’ tag RNA-seq libraries and please retain CIF images files from seq run” and submit the job, take note to the Job Request number and wait for the JA number (It can take one or two days).
   - In requested platform: HiSeq 2500 run
   - In requested RunType: SR 100

4. Meanwhile do the mix with the data from qPCR (Step 5e).

5. Put the JA number int the tube (cap and body) and bring it to the GSAF facility room MBB 4.102. And wait the long queue.... for the data.

### Plant RNA Extractions

1. Grind approx 0.1 g tissue by GenoGrinder to a good powder.

2. Add 1 mL of Trizol reagent to the ground powder in the tube and mix immediately.

*Note: Do not let ground plant tissue thaw without being in the presence of Trizol. The sample in Trizol can be stored at -80 ℃ for long term.*

3. Add 200 µL of chloroform and shake vigorously by hand for approx 50 sec.

4. Let stand at room temperature for 10 min.

5. Centrifuge at 12,000g for 15 min at room temperature.

6. Carefully transfer 600 ul upper aqueous phase to a new tube

*Note: ensure no interface debris is transferred.*

7. Add 400 ul of isopropyl alcohol and invert to mix well.

8. Centrifuge at 12,000g for 15 min at room temperature.

9. Carefully discard supernatant.

*Note: You can see the pellet at this step.*

10. Add 500 ul cold 75% ethanol and invert the tube to float the pellet.

*Note: The pellet of RNA can be stored in 75% ethanol at -80 ℃ for long term.*

11. Centrifuge at 12,000g for 5 min at at room temperature.

12. Discard the supernatant. Centrifuge again for a short time and use the pipette to remove the residue.

13. Put the tube on the laminar flow bench and allow the pellet to air-dry for 10 min.

*Note: Usually the pellet will turn clear after 10-20 min.*

14. Dissolve the pellet in 30 μL of RNase-free water by very gently sucking the liquid up and down with a pipet.

15. Quantify the RNA, check the purity and degradation, and either store at –20 or –80°C until used.
