## Supplementary Figures for "Gene expression divergence between locally adapted inland annual and coastal perennial ecotypes of *Mimulus guttatus* across developmental stages"

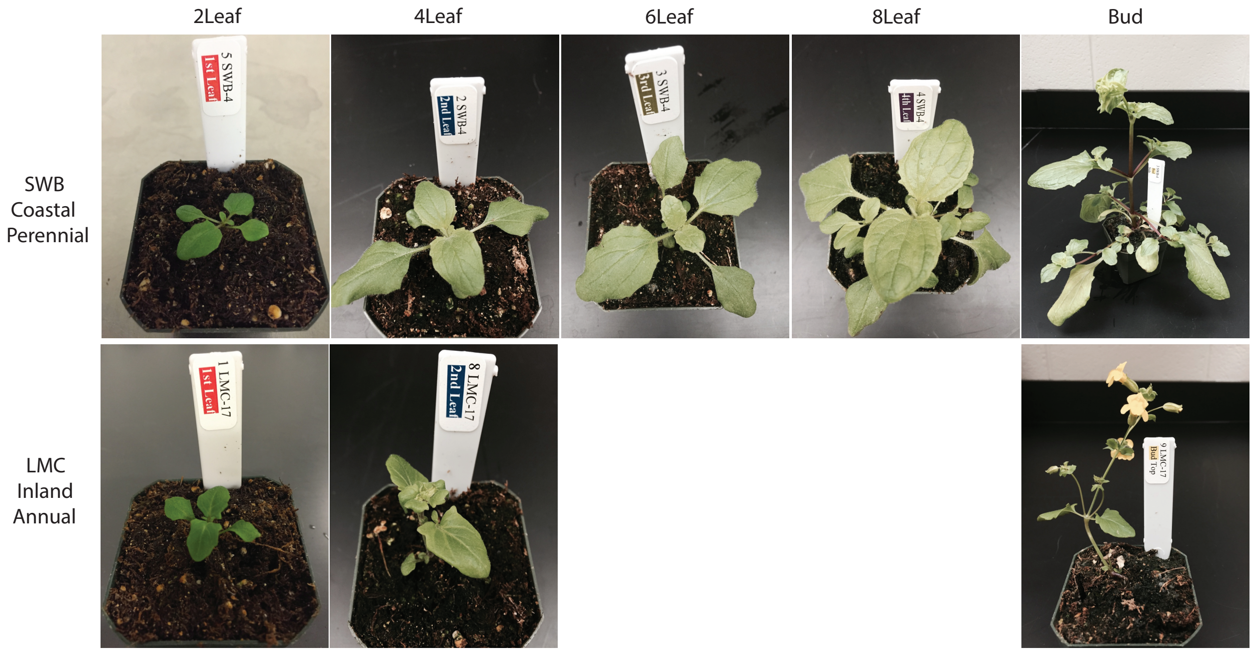


**Figure S1.** Photos of experimental plants on the day of tissue collection for RNA-seq through developmental stages for the two genotypes.

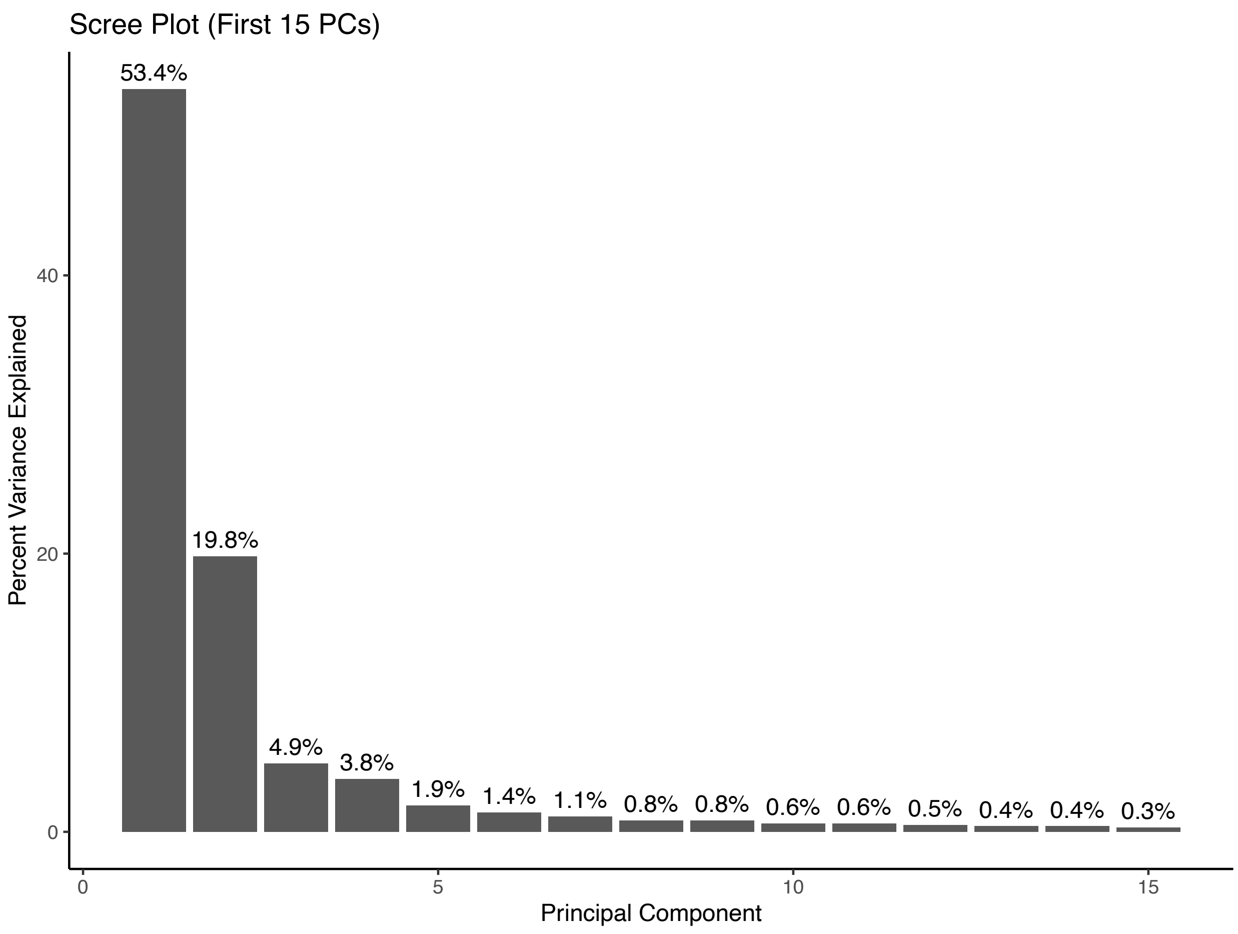


**Figure S2.** Scree plot of variance explained by the first 15 PCs for the principal component analysis (PCA) performed on variance stabilizing transformed (VST), pre-filtered read count data.

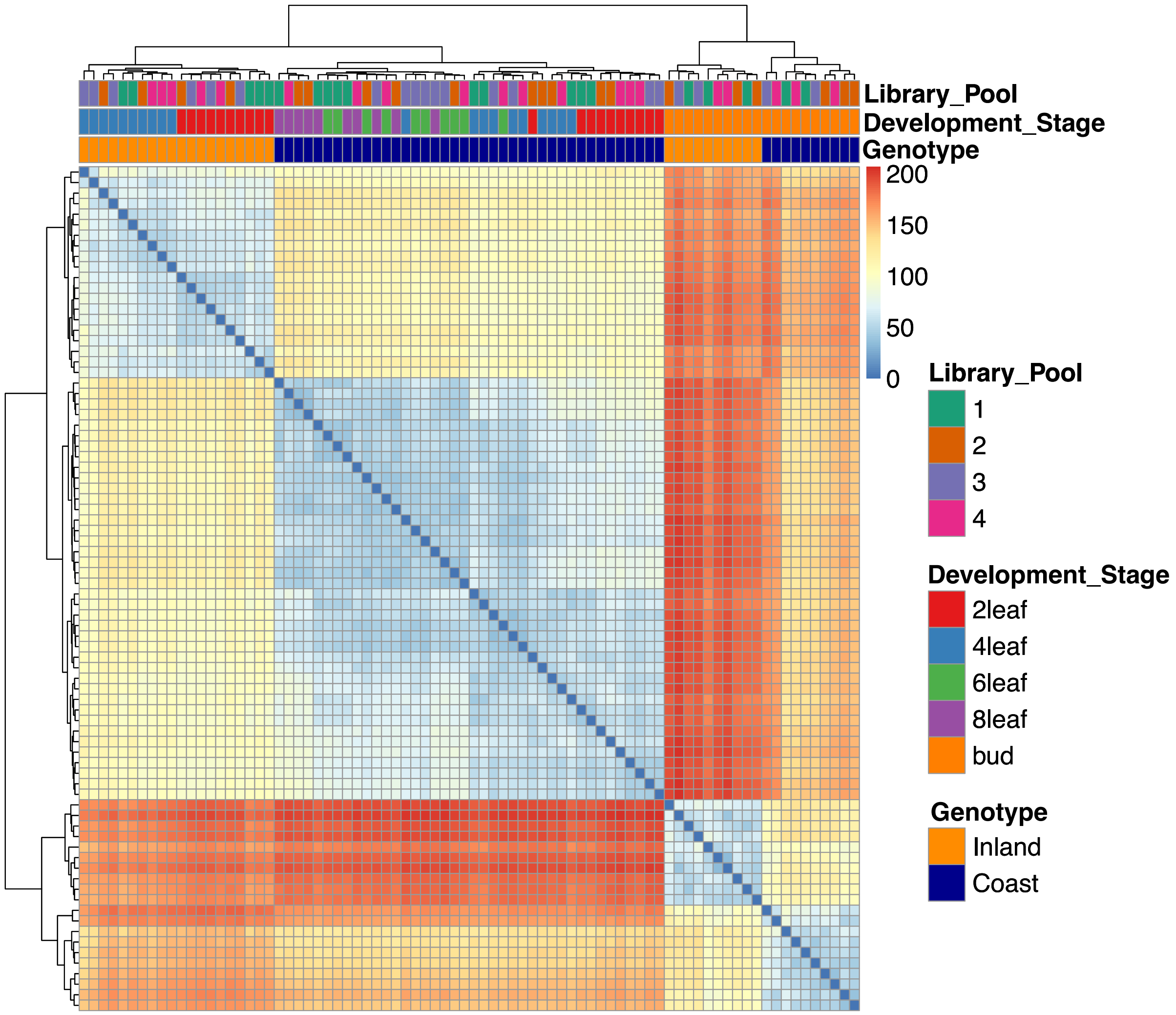


**Figure S3.** Heat map clustering of the 80 RNA-seq samples by genotype and developmental stage. There was no clustering by the four library pools, across which the samples were randomized.

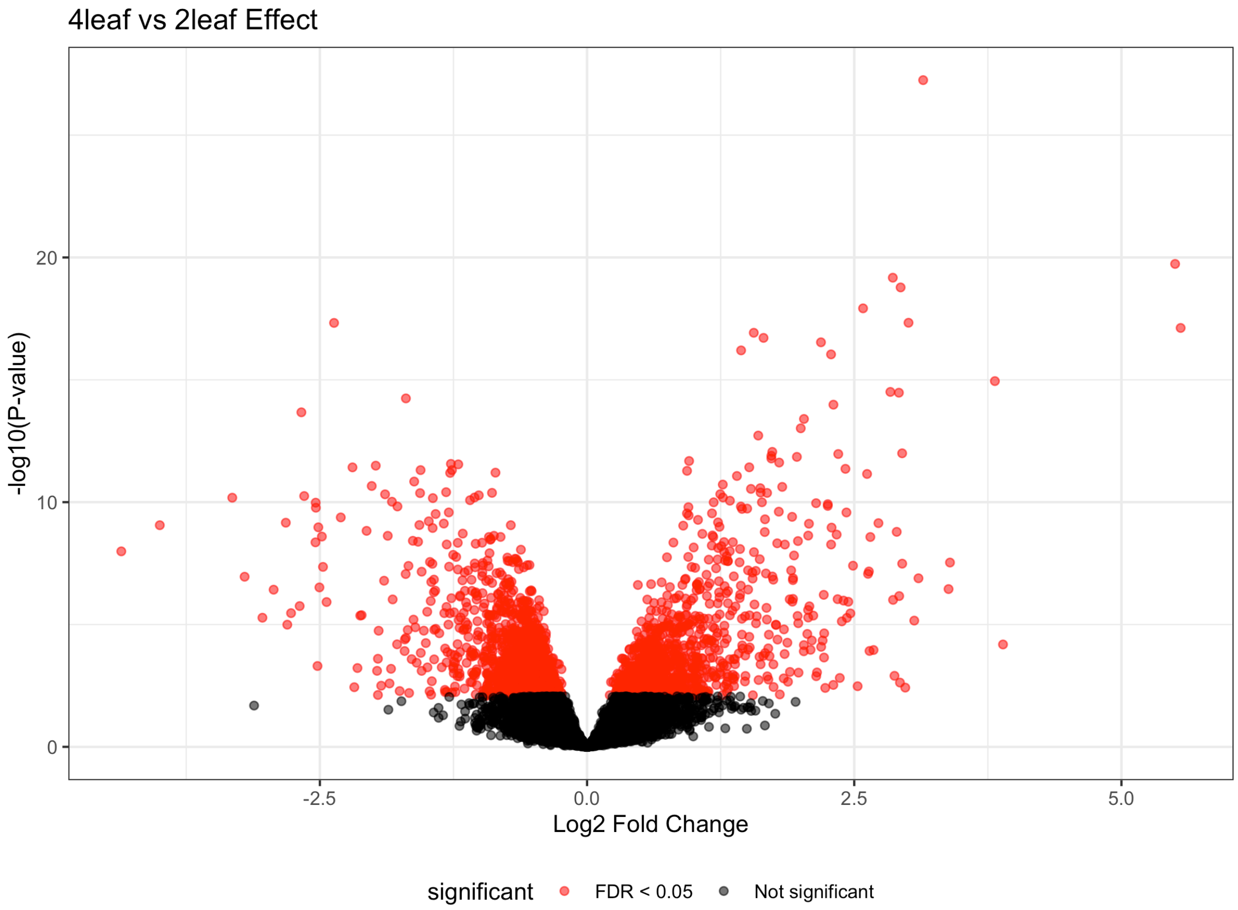


**Figure S4.** Volcano plot depicting the distribution of significantly differentially expressed genes (red) for the contrast between the second and fourth set of true leaves.

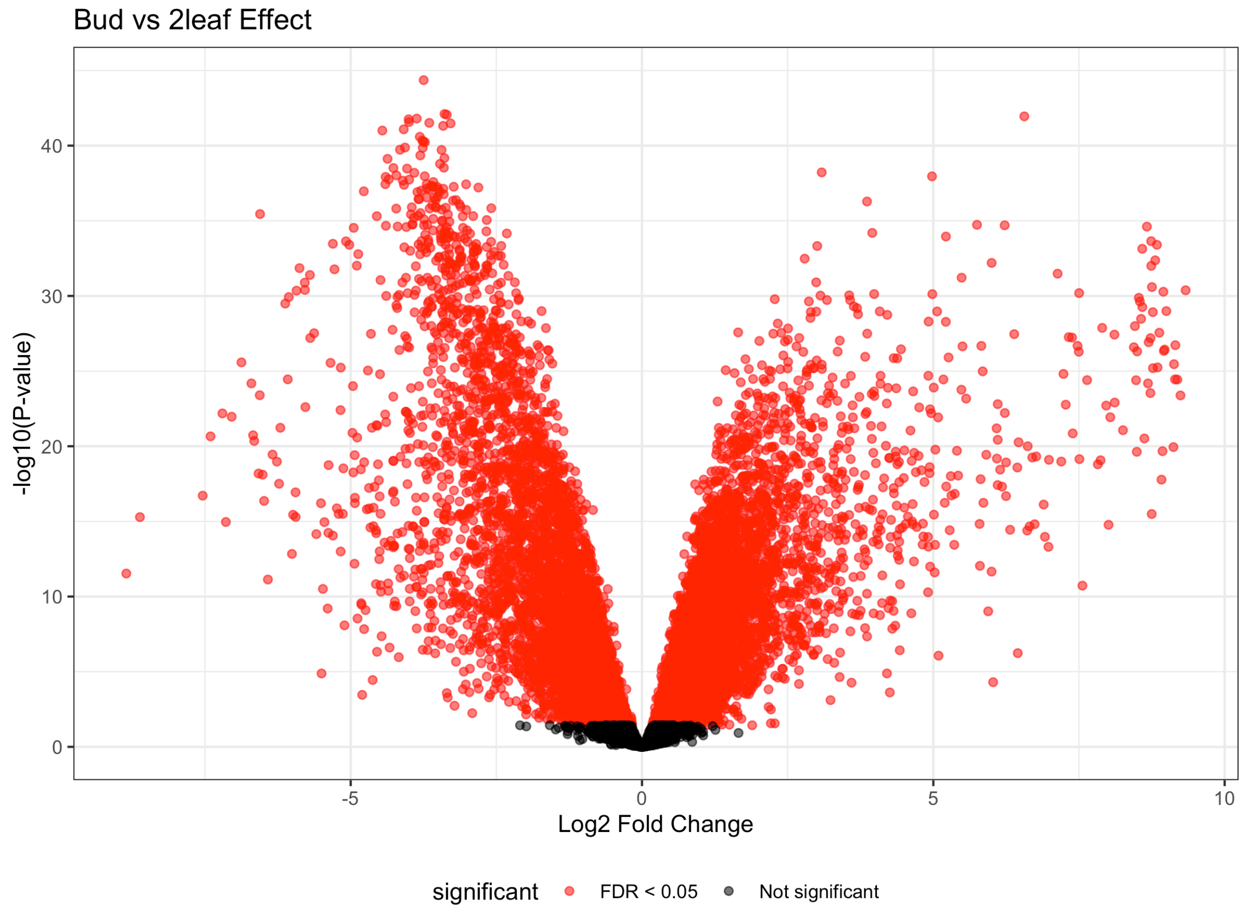


**Figure S5.** Volcano plot depicting the distribution of significantly differentially expressed genes (red) for the contrast between floral buds and the second set of true leaves.

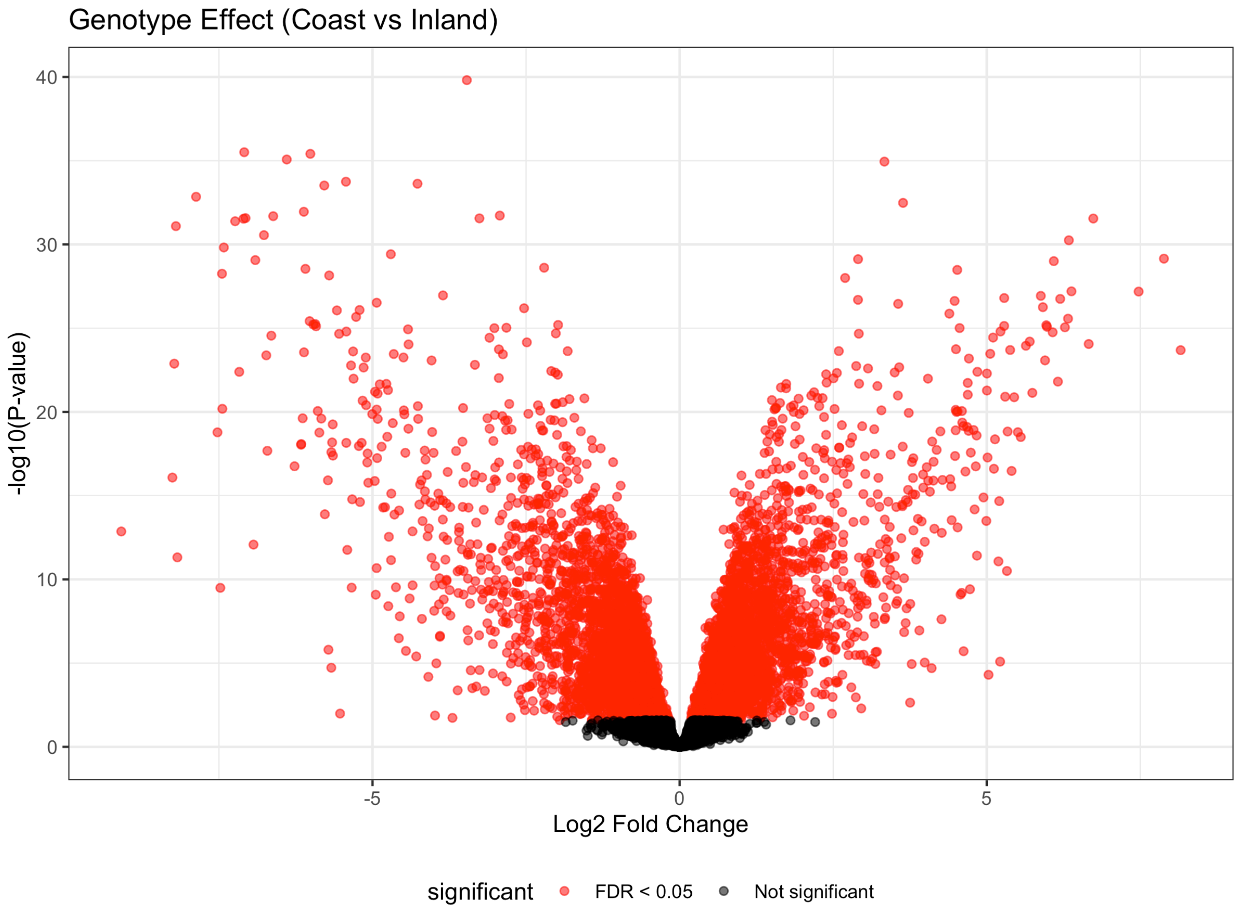


**Figure S6.** Volcano plot depicting the distribution of significantly differentially expressed genes (red) for the contrast between coastal perennial and inland annuall samples.

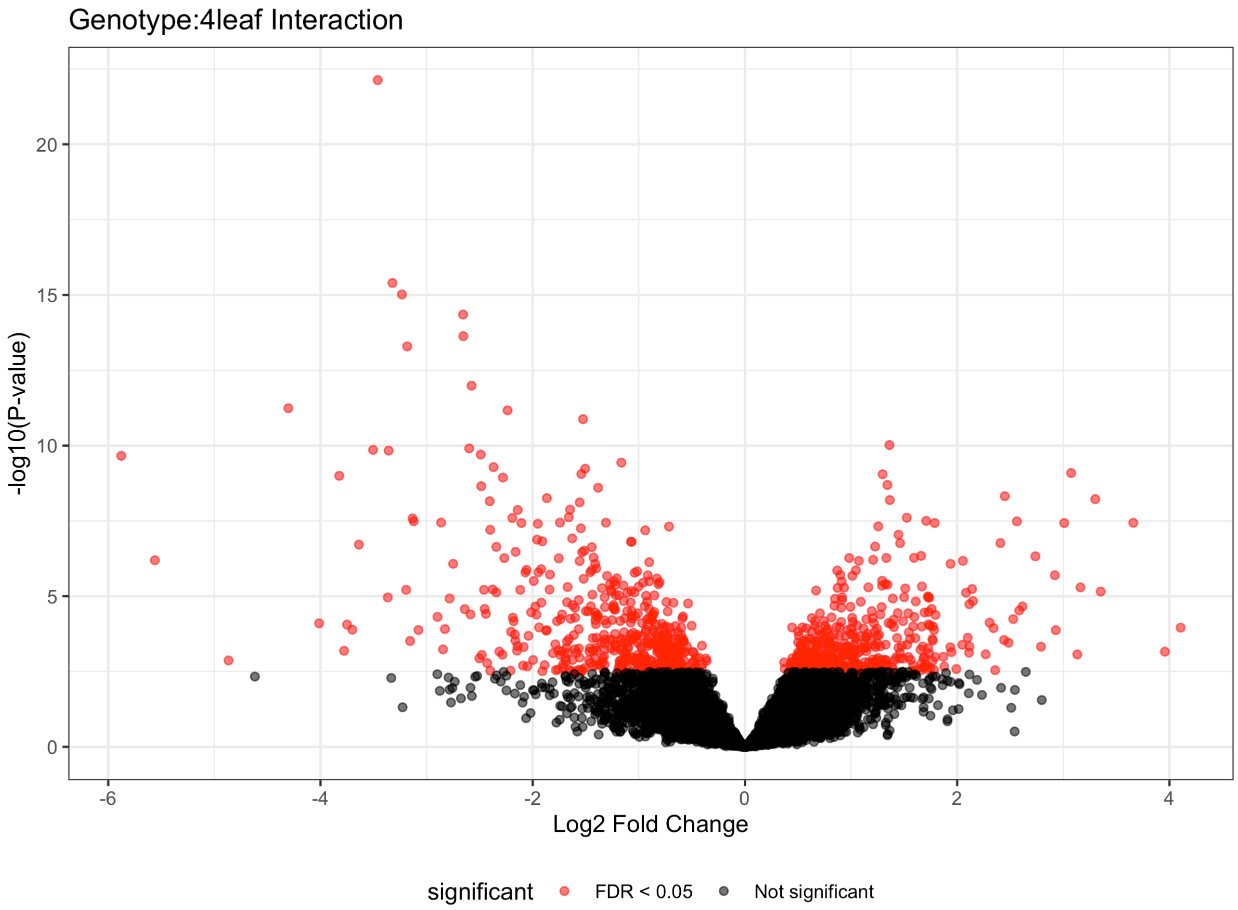


**Figure S7.** Volcano plot depicting the distribution of significantly differentially expressed genes (red) for the interaction of genotype with the contrast of the second and fourth set of true leaves.

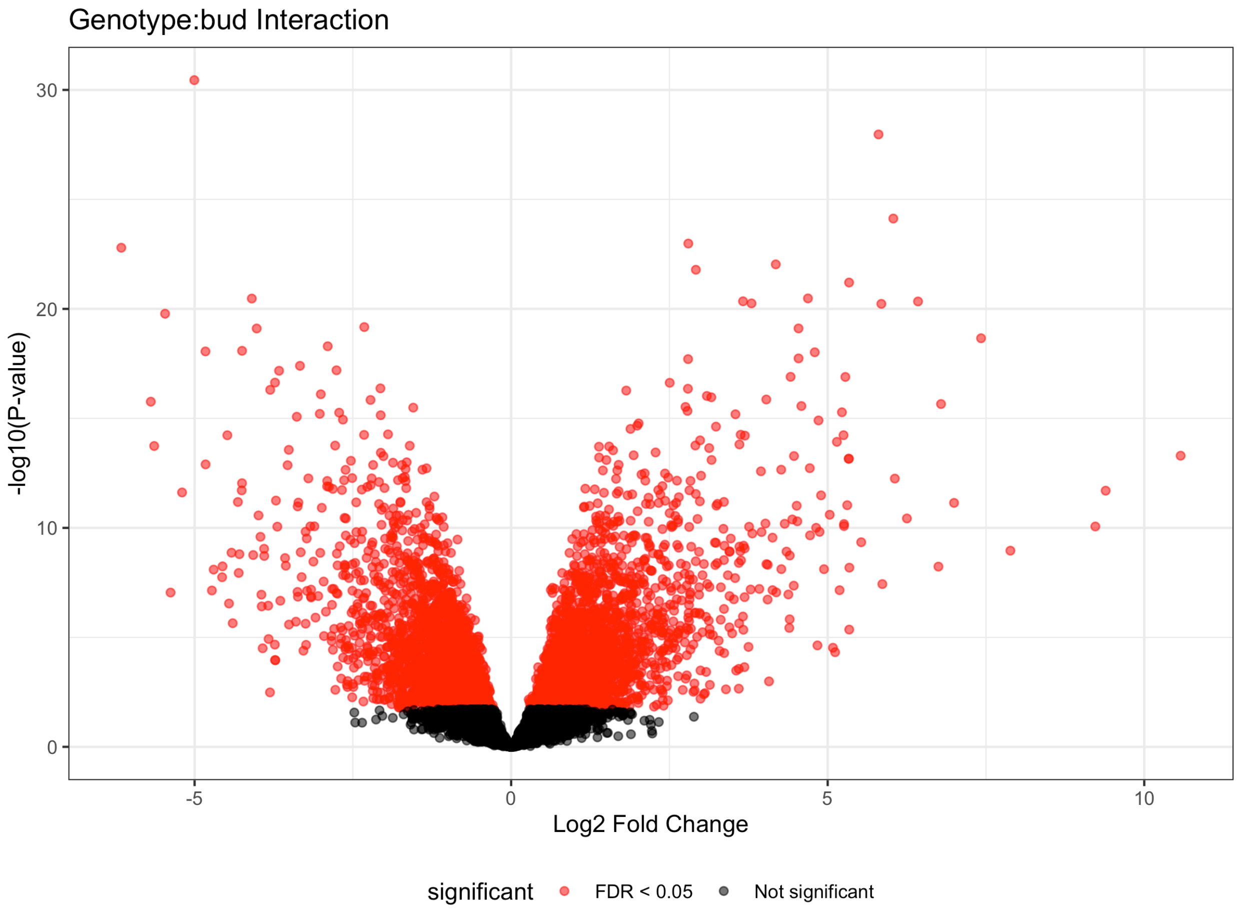


**Figure S8.** Volcano plot depicting the distribution of significantly differentially expressed genes (red) for the interaction of genotype with the contrast of floral bud and the second set of true leaves.

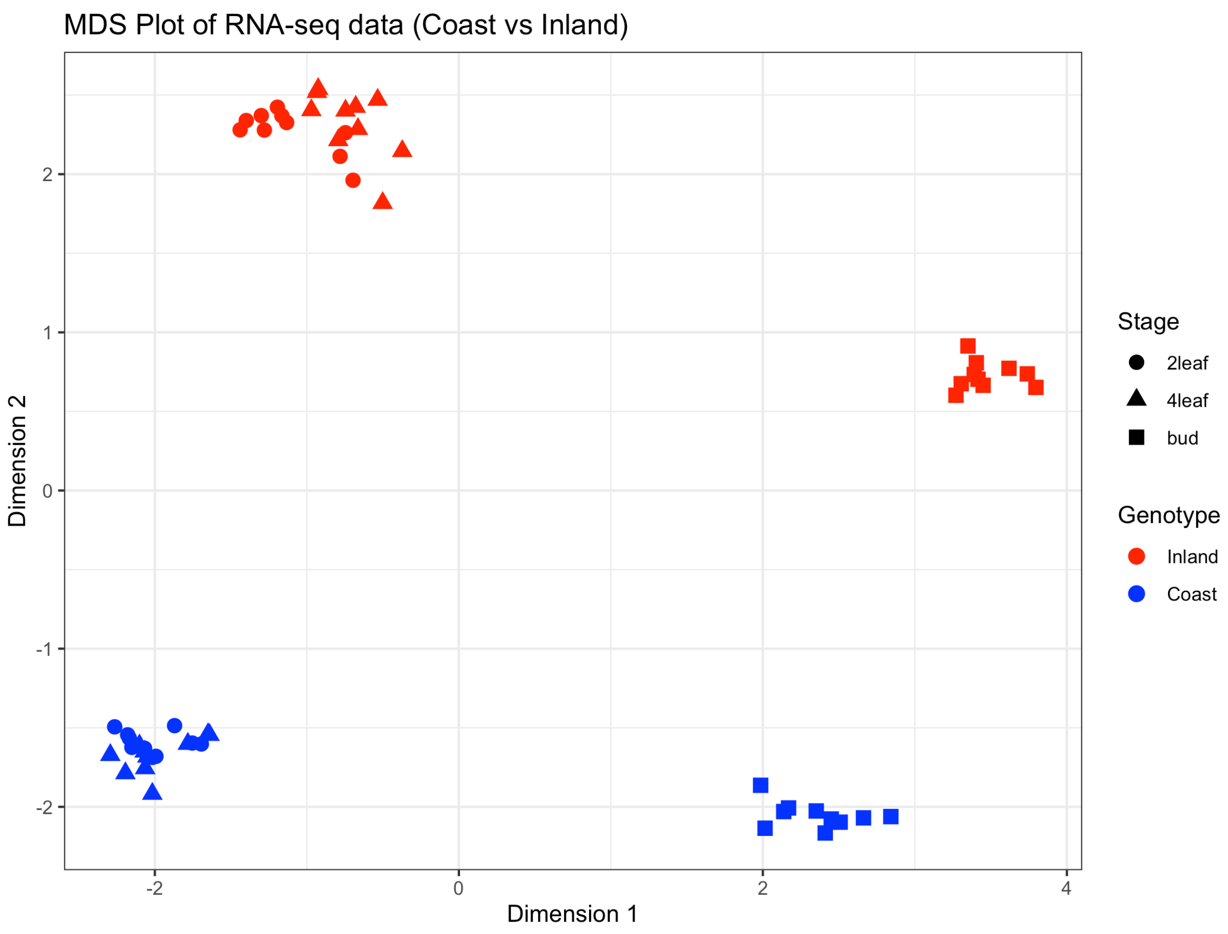


**Figure S9.** Multidimensional scaling (MDS) plot of the relationship of samples based on the 13,920 genes individually evaluated for differential expression using the voom-limma approach.

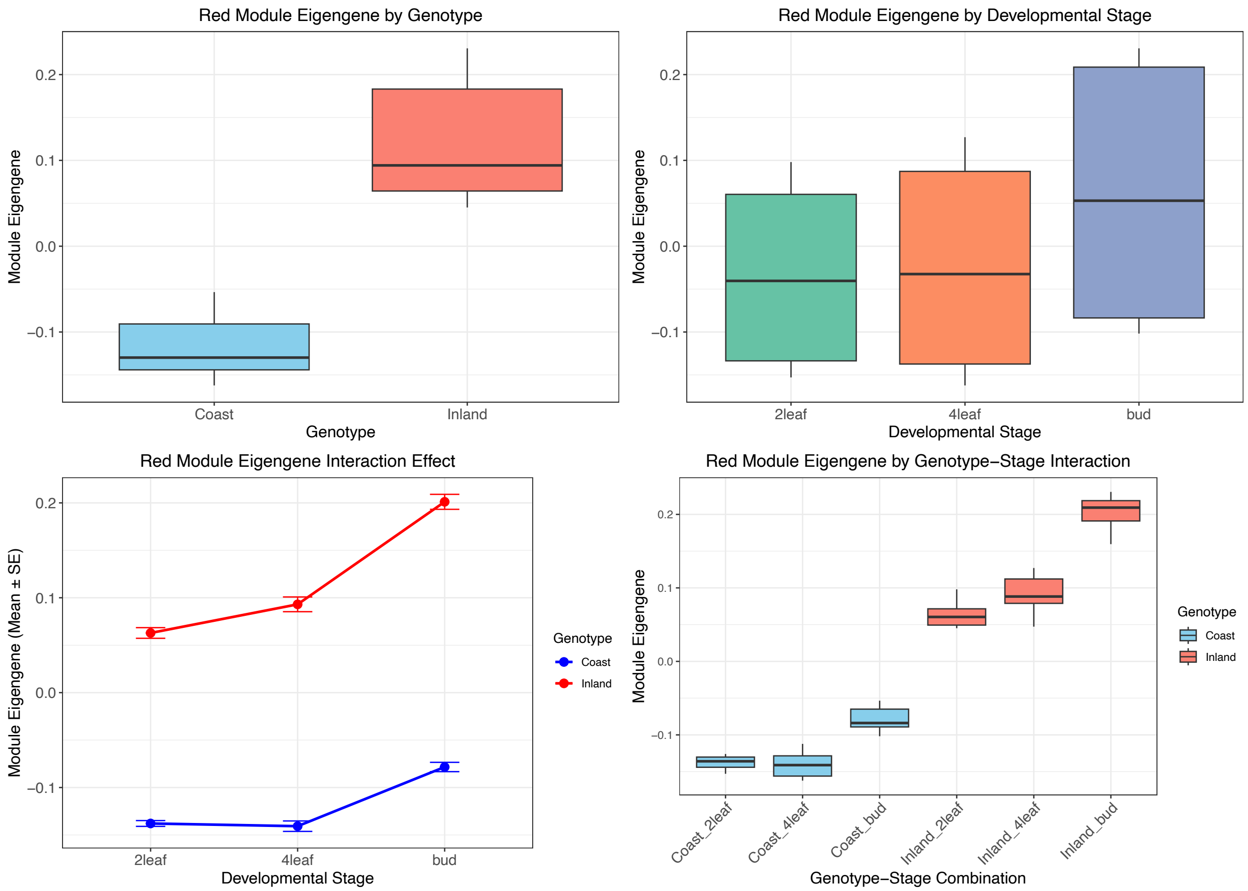


**Figure S10.** The main effects (genotype and developmental stage) and interactions for the eigengene of the red module from the WGCNA analysis.

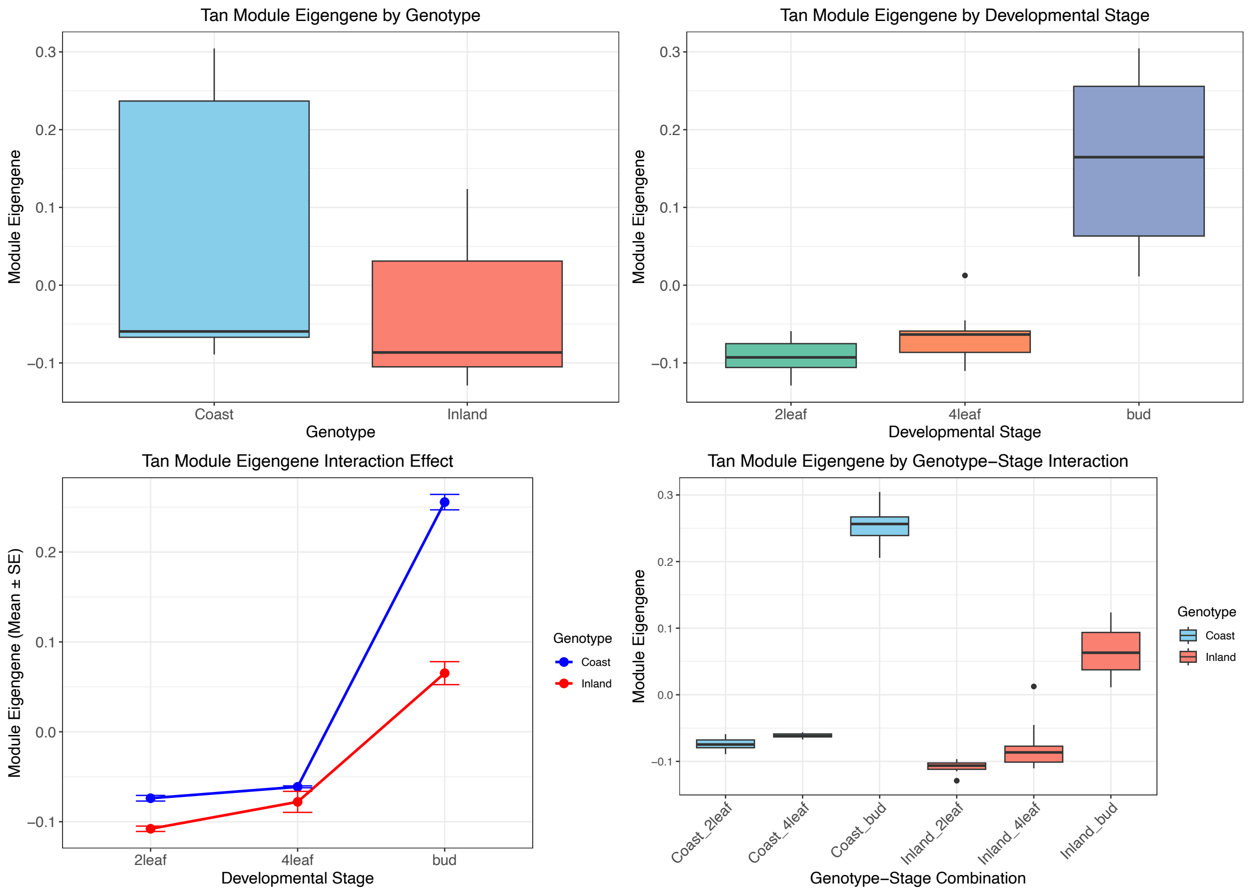


**Figure S11.** The main effects (genotype and developmental stage) and interactions for the eigengene of the tan module from the WGCNA analysis.

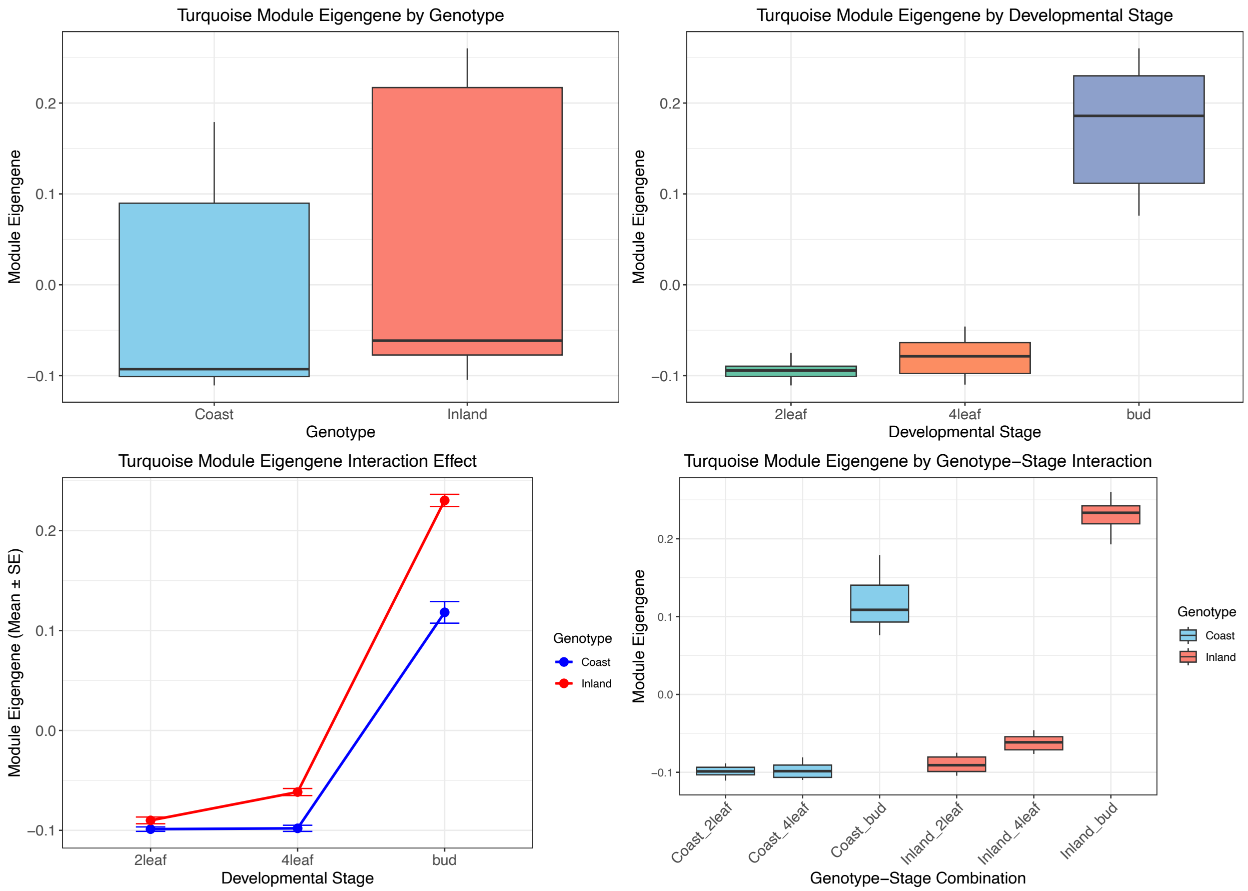


**Figure S12.** The main effects (genotype and developmental stage) and interactions for the eigengene of the turquoise module from the WGCNA analysis.

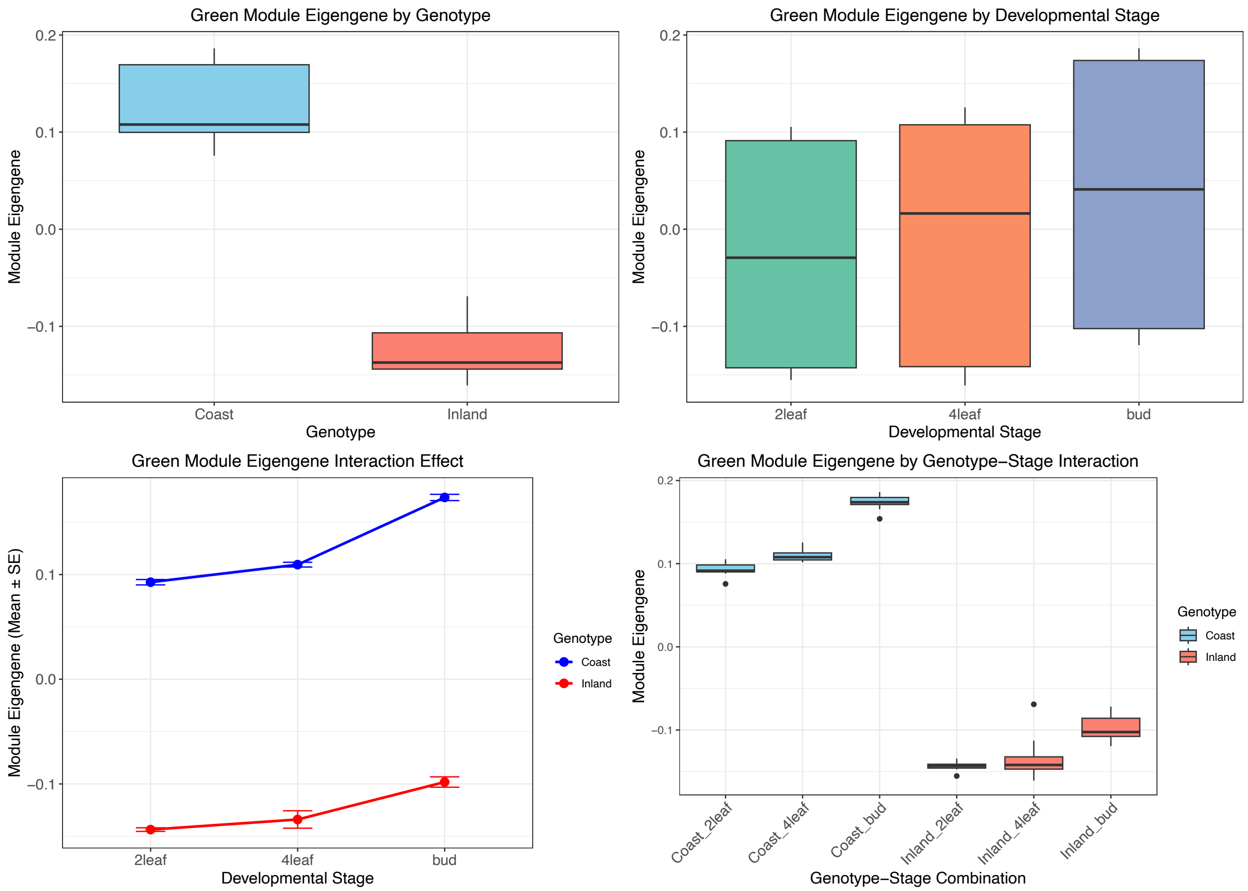


**Figure S13.** The main effects (genotype and developmental stage) and interactions for the eigengene of the green module from the WGCNA analysis.

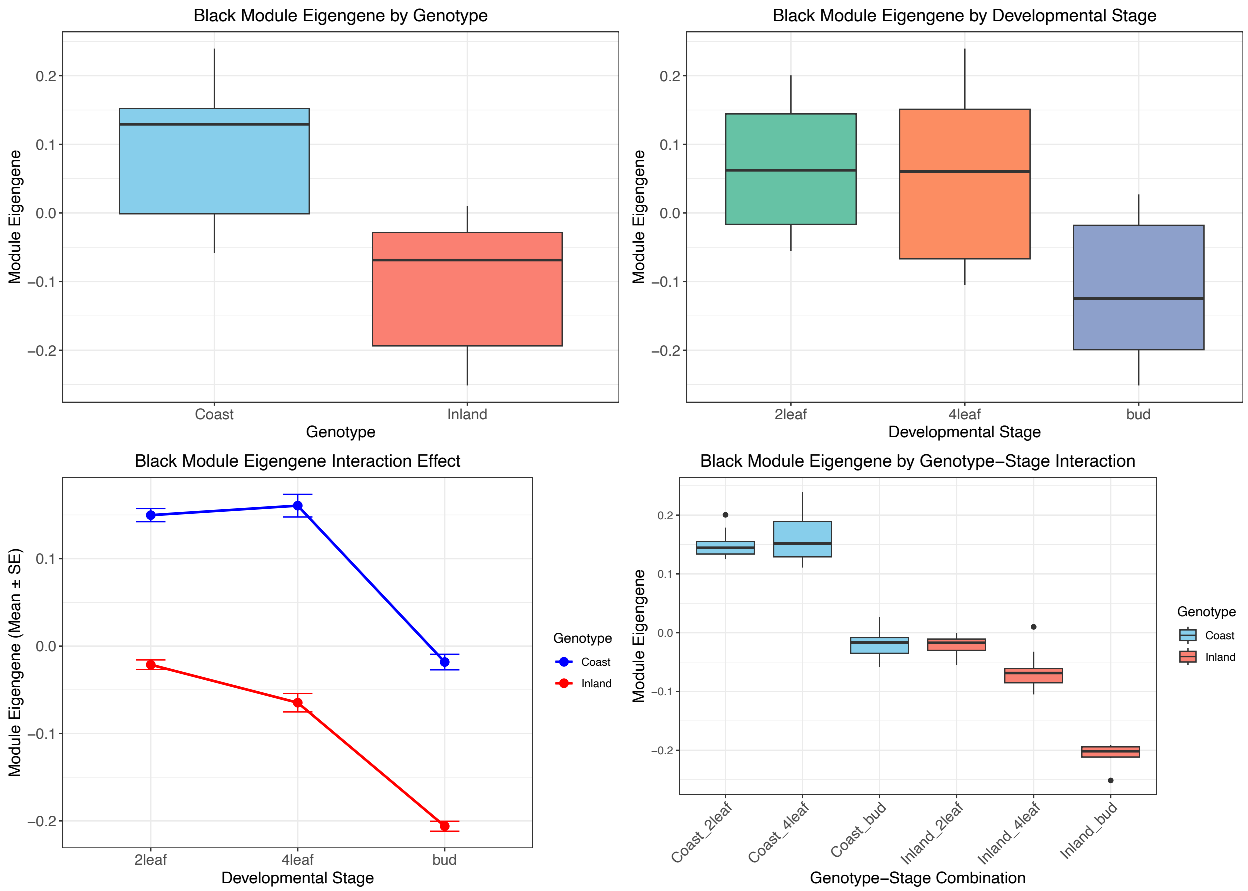


**Figure S14.** The main effects (genotype and developmental stage) and interactions for the eigengene of the black module from the WGCNA analysis.

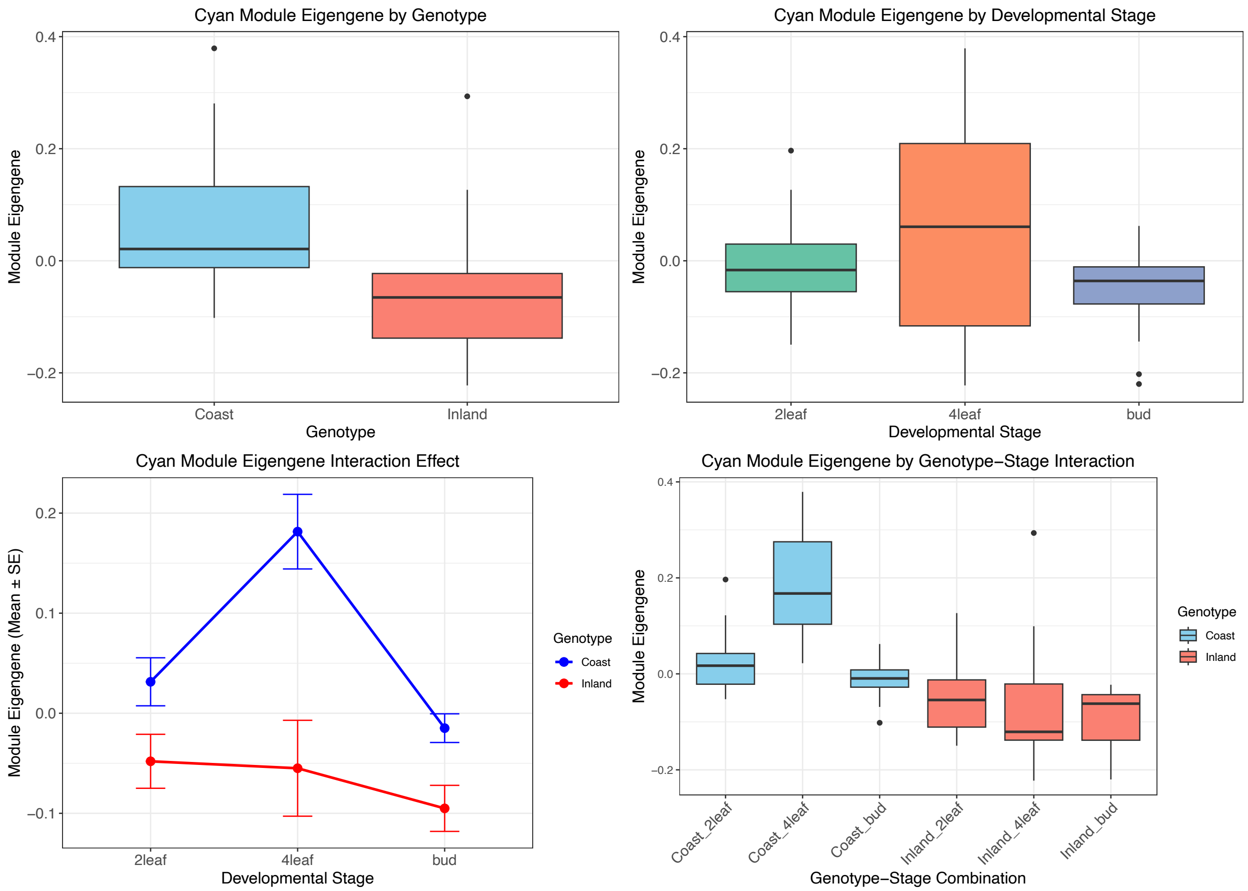


**Figure S15.** The main effects (genotype and developmental stage) and interactions for the eigengene of the cyan module from the WGCNA analysis.

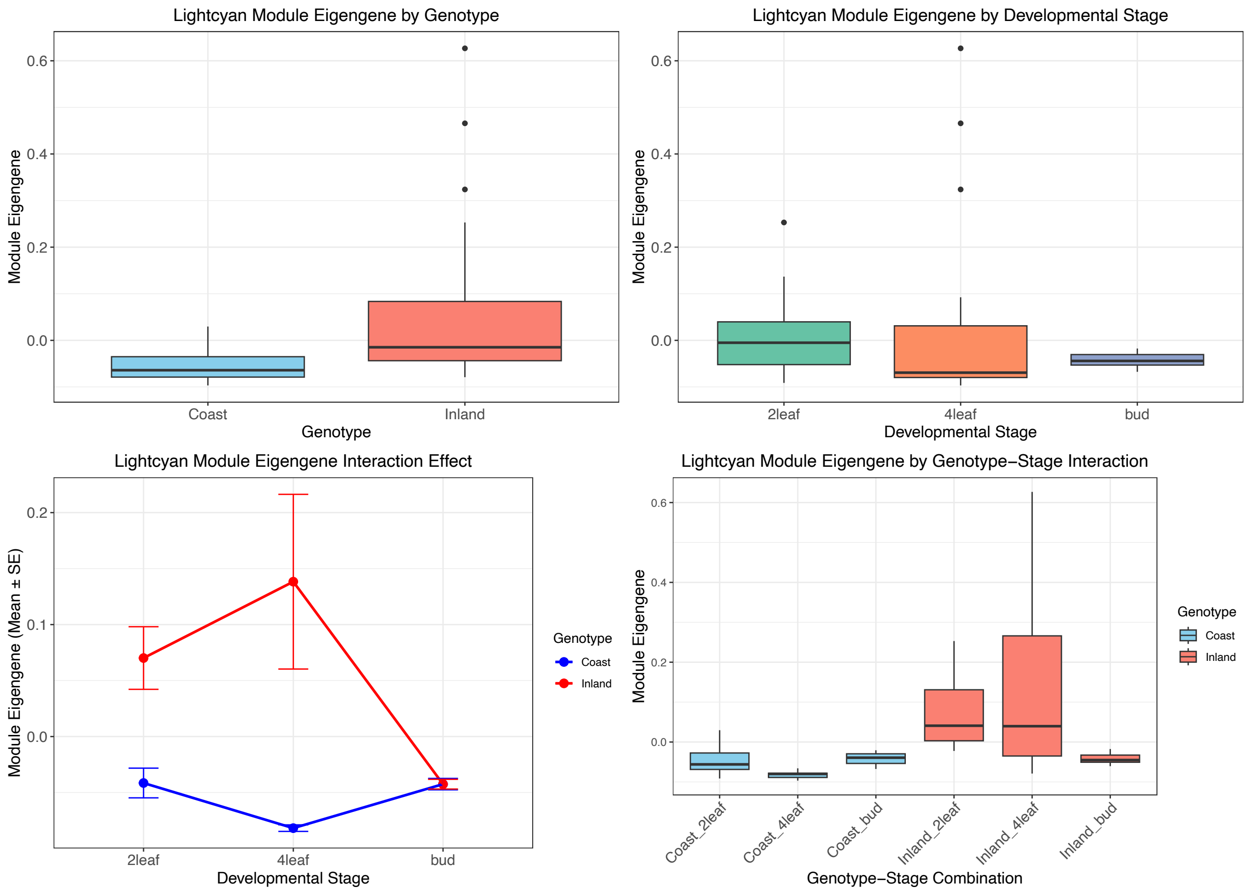


**Figure S16.** The main effects (genotype and developmental stage) and interactions for the eigengene of the lightcyan module from the WGCNA analysis.

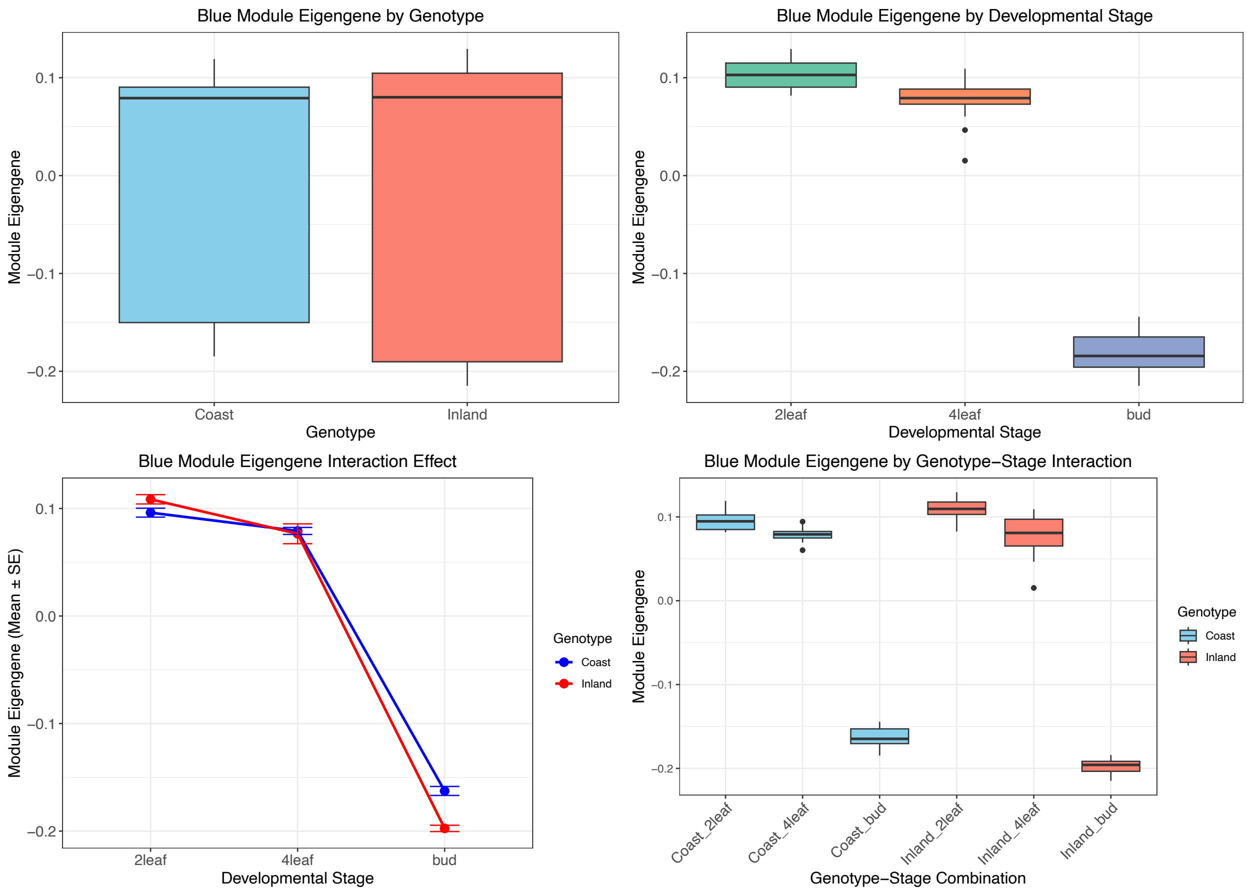


**Figure S17.** The main effects (genotype and developmental stage) and interactions for the eigengene of the blue module from the WGCNA analysis.

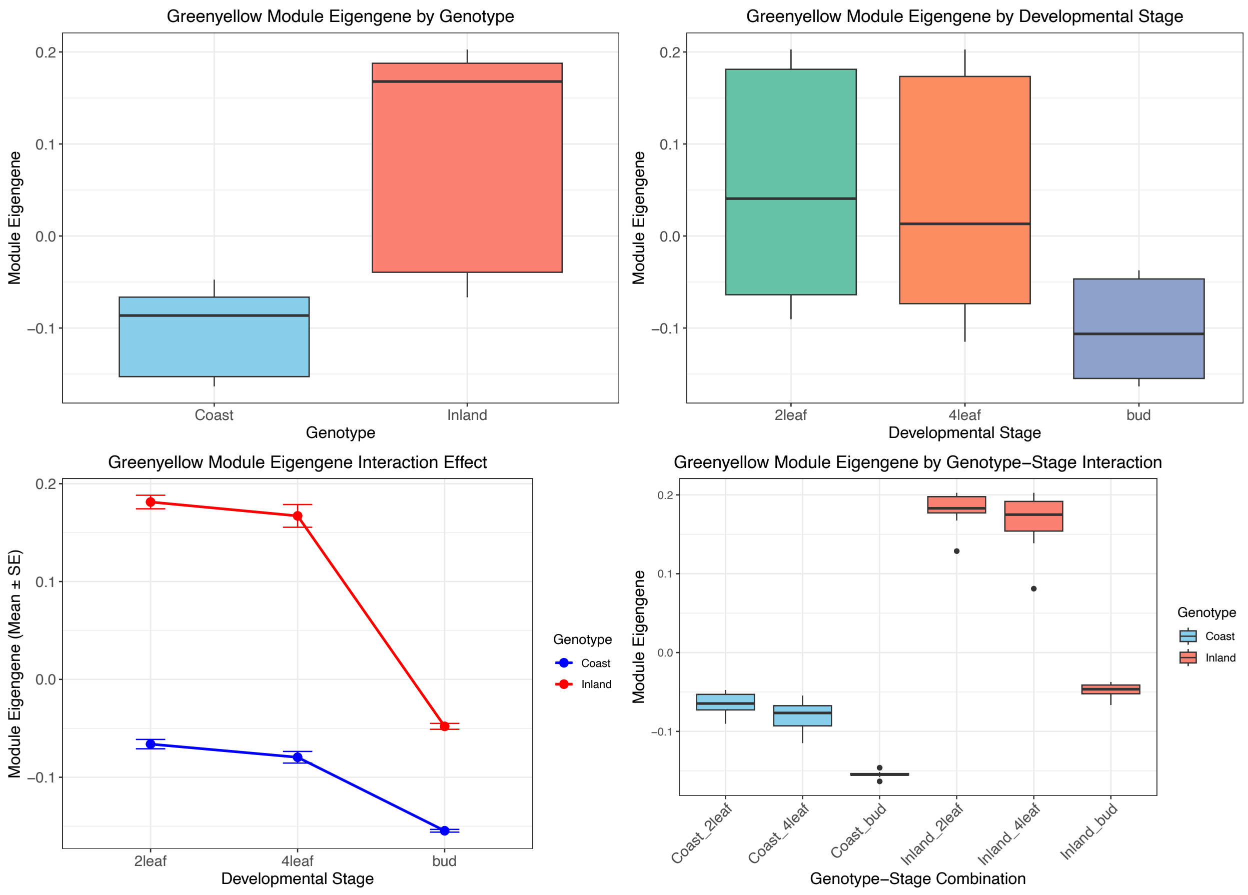


**Figure S18.** The main effects (genotype and developmental stage) and interactions for the eigengene of the greenyellow module from the WGCNA analysis.

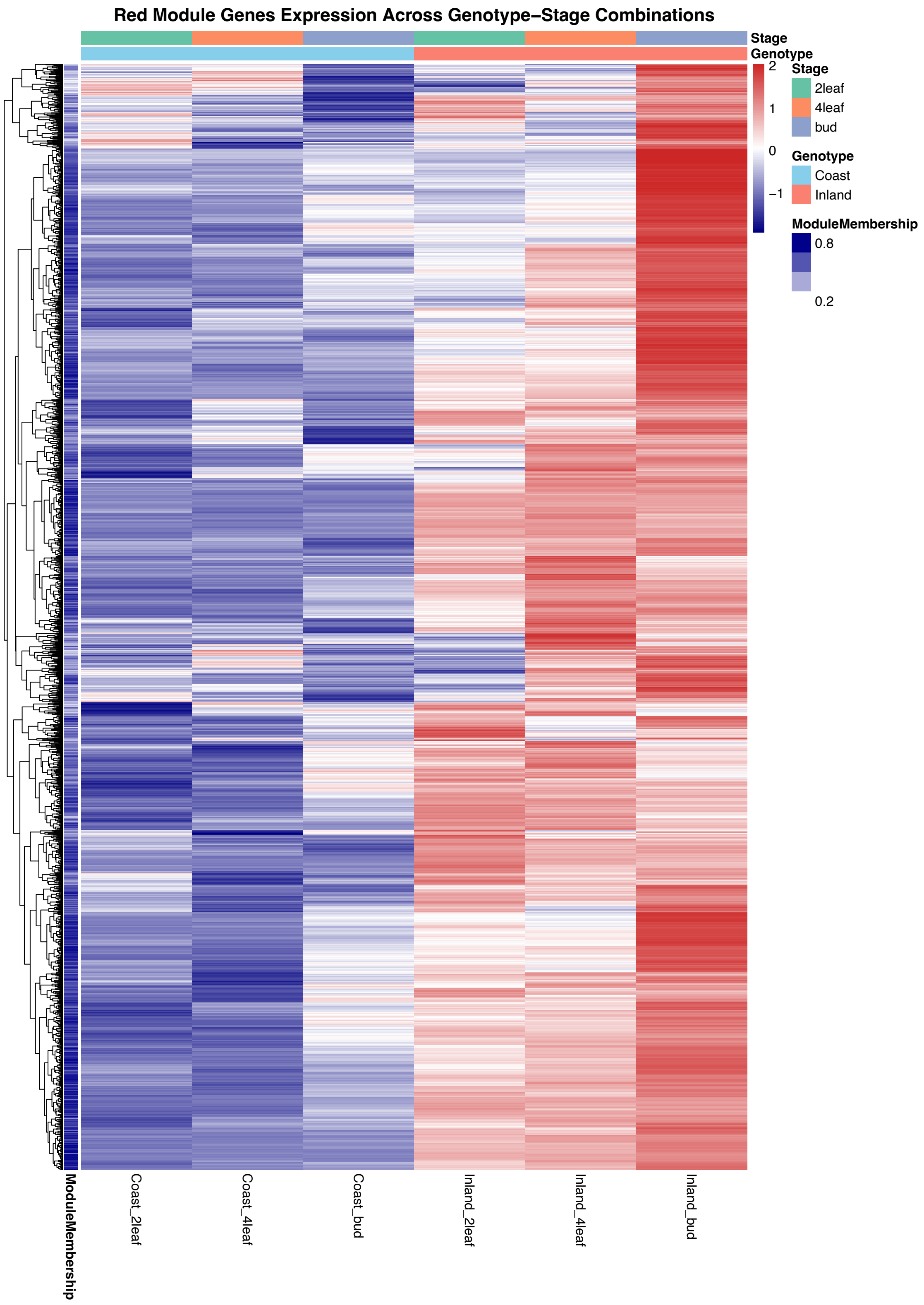


**Figure S19.** Heatmap of the expression of genes within the red module across genotypes and developmental stages.

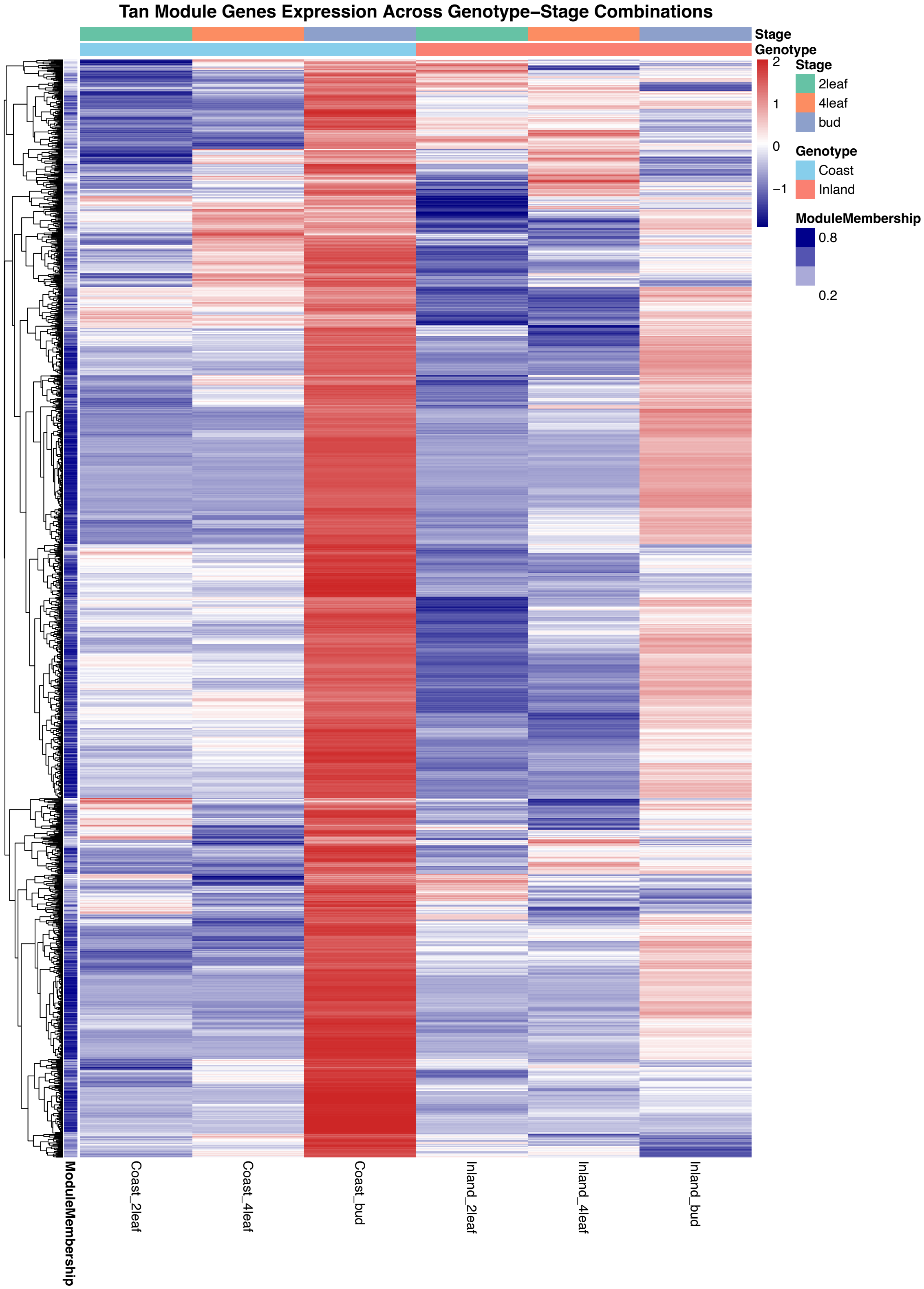


**Figure S20.** Heatmap of the expression of genes within the tan module across genotypes and developmental stages.

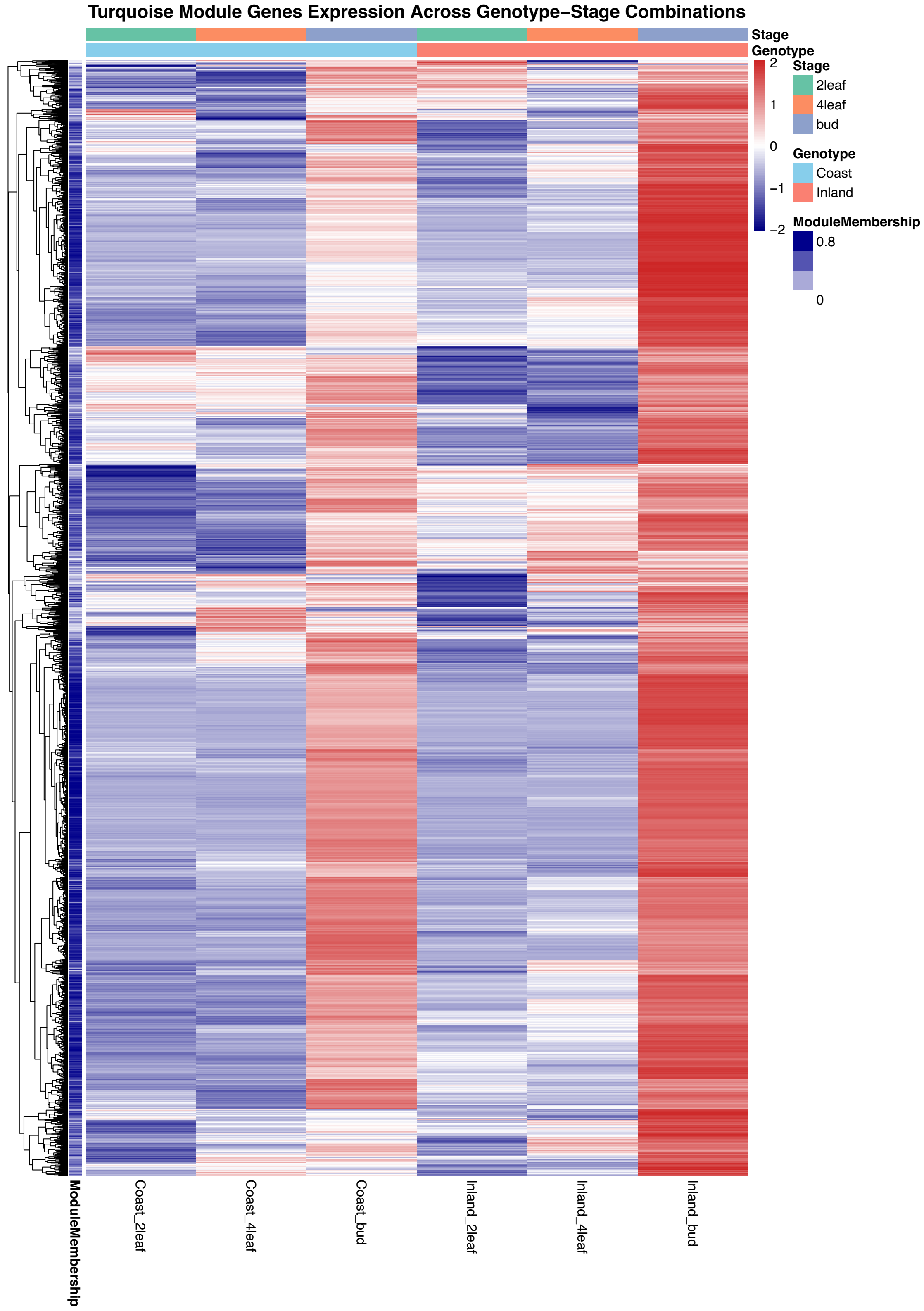


**Figure S21.** Heatmap of the expression of genes within the turquoise module across genotypes and developmental stages.

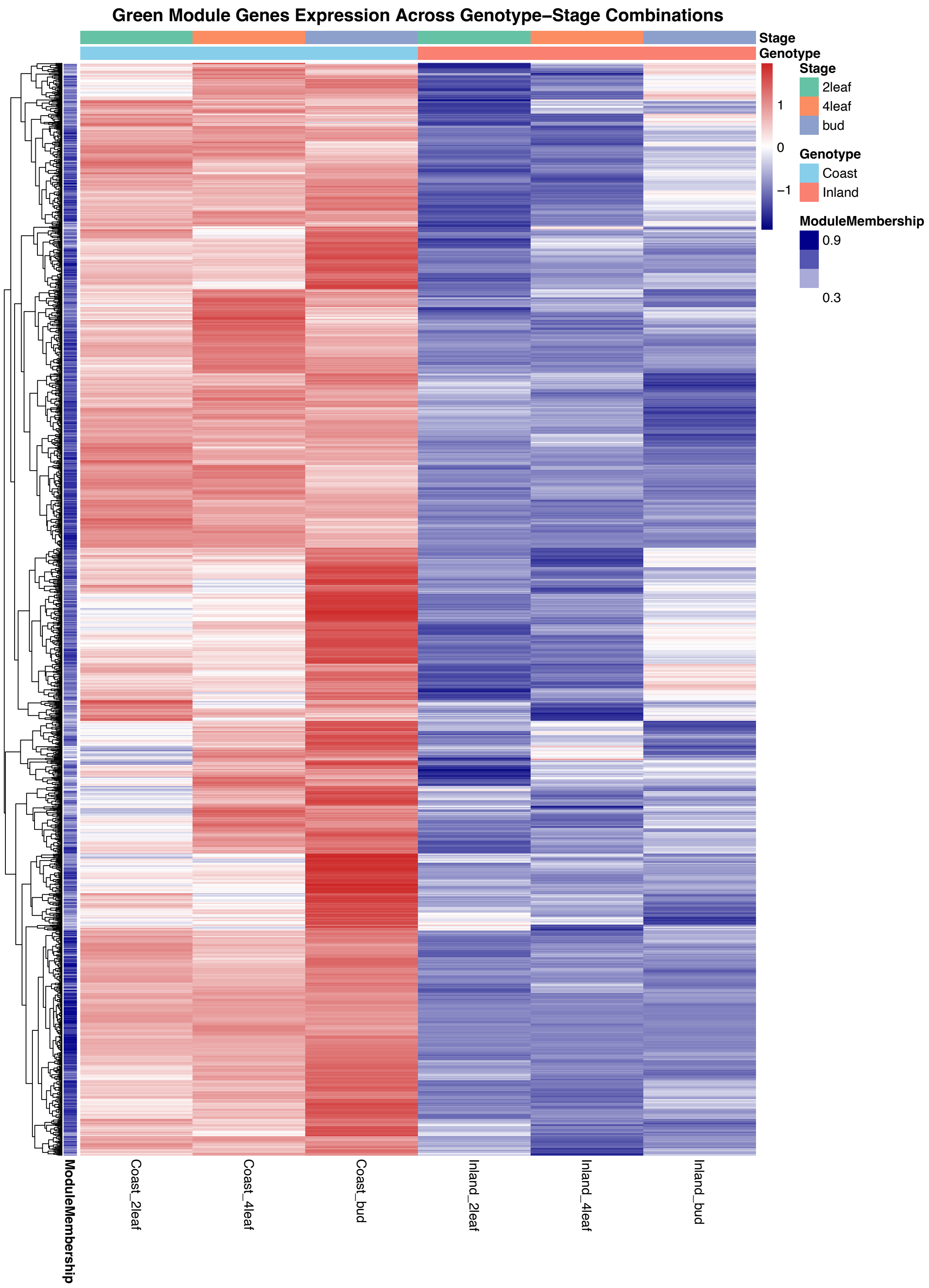


**Figure S22.** Heatmap of the expression of genes within the green module across genotypes and developmental stages.

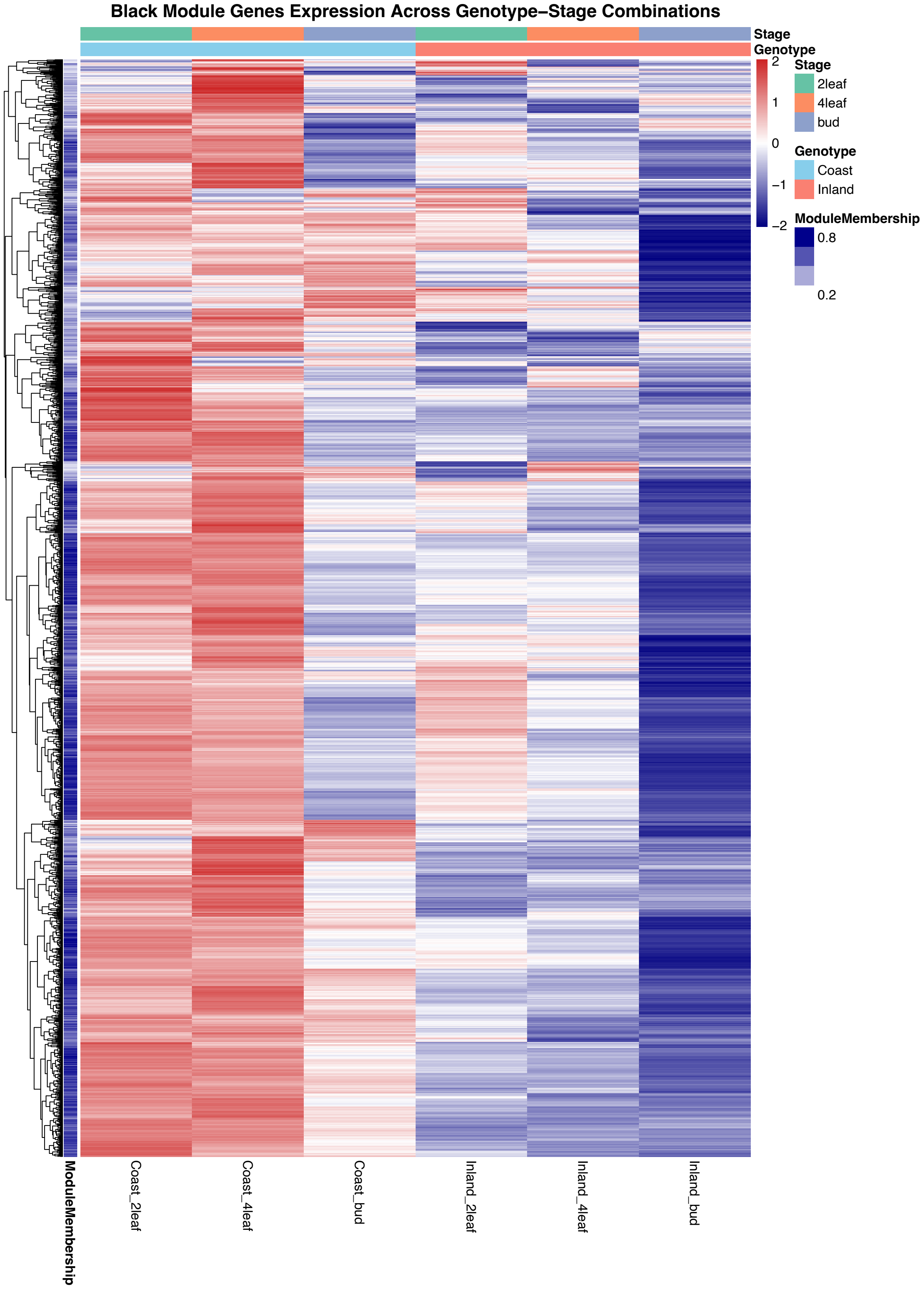


**Figure S23.** Heatmap of the expression of genes within the black module across genotypes and developmental stages.

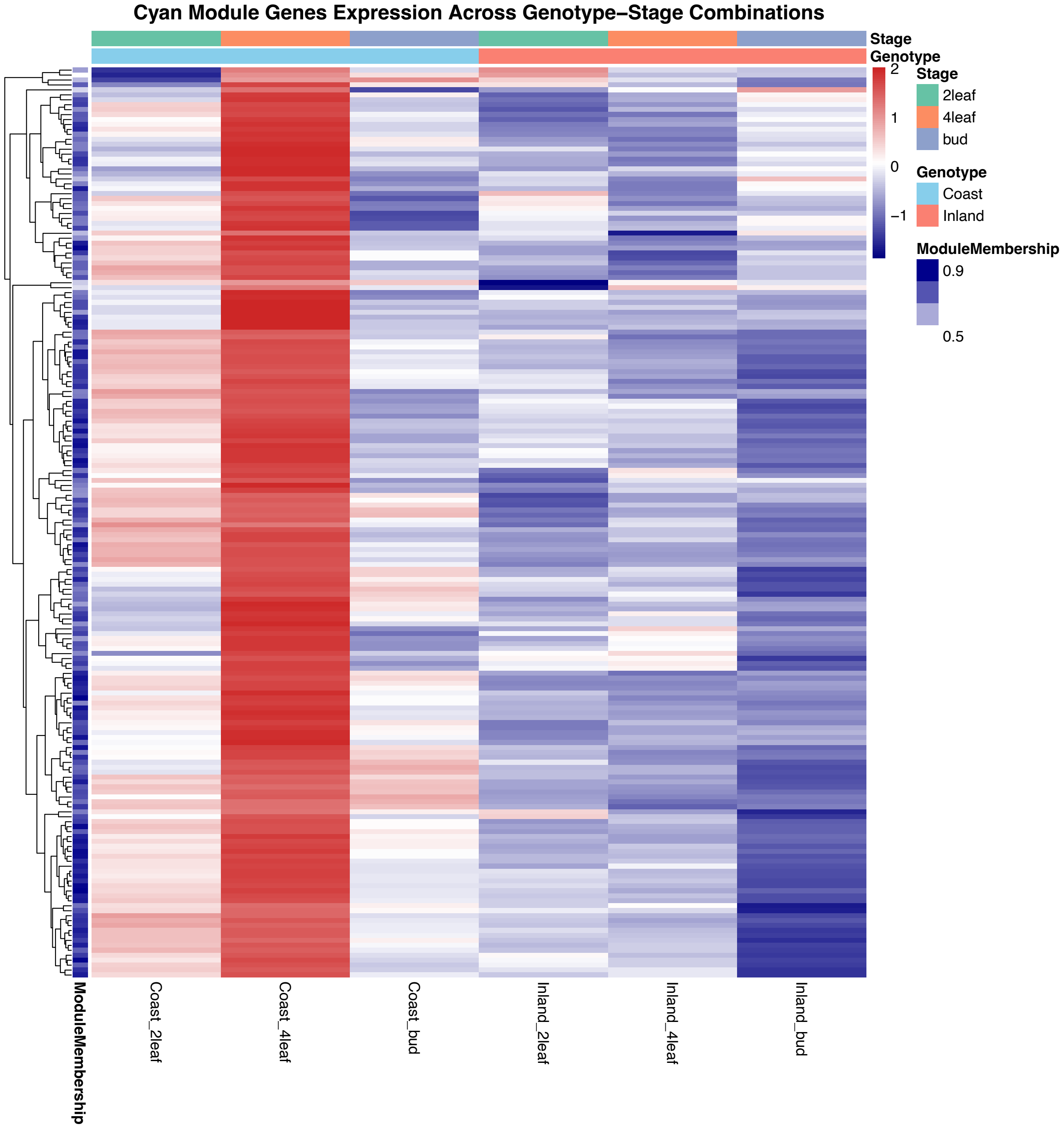


**Figure S24.** Heatmap of the expression of genes within the cyan module across genotypes and developmental stages.

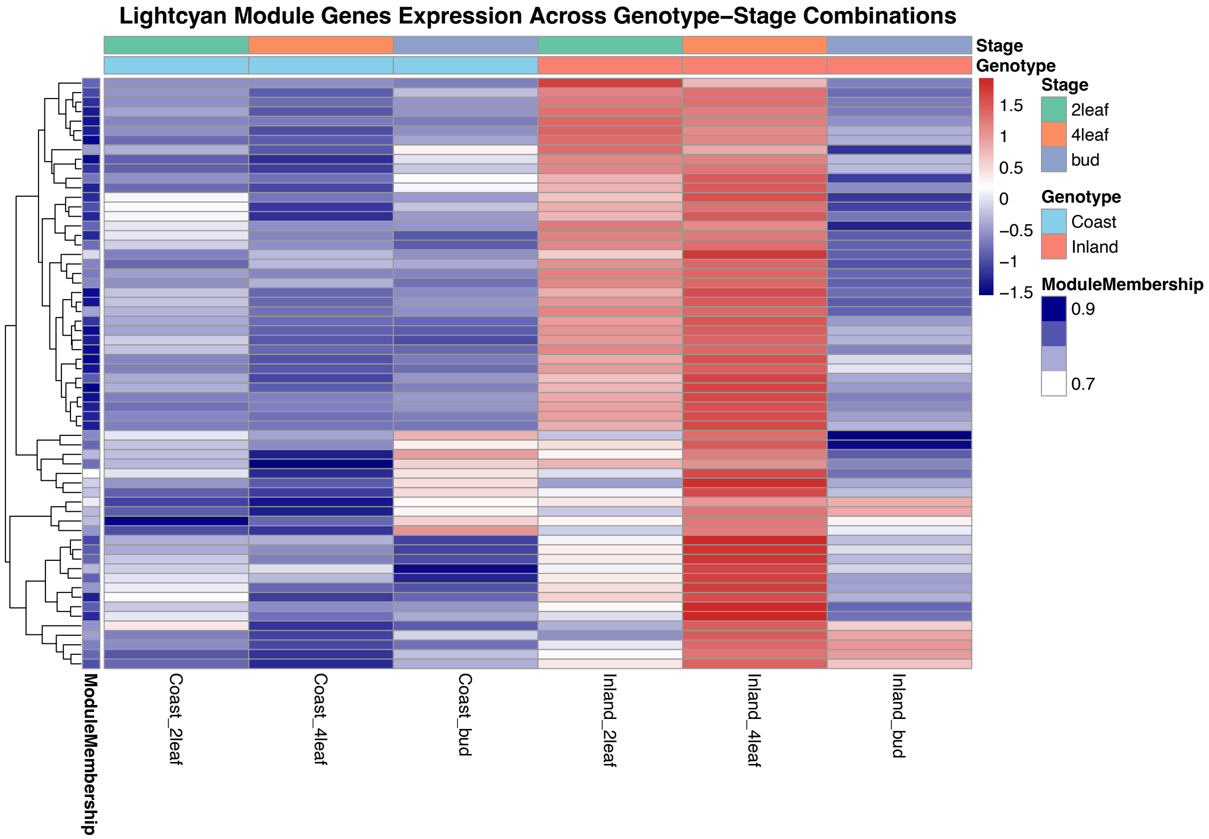


**Figure S25.** Heatmap of the expression of genes within the lightcyan module across genotypes and developmental stages.

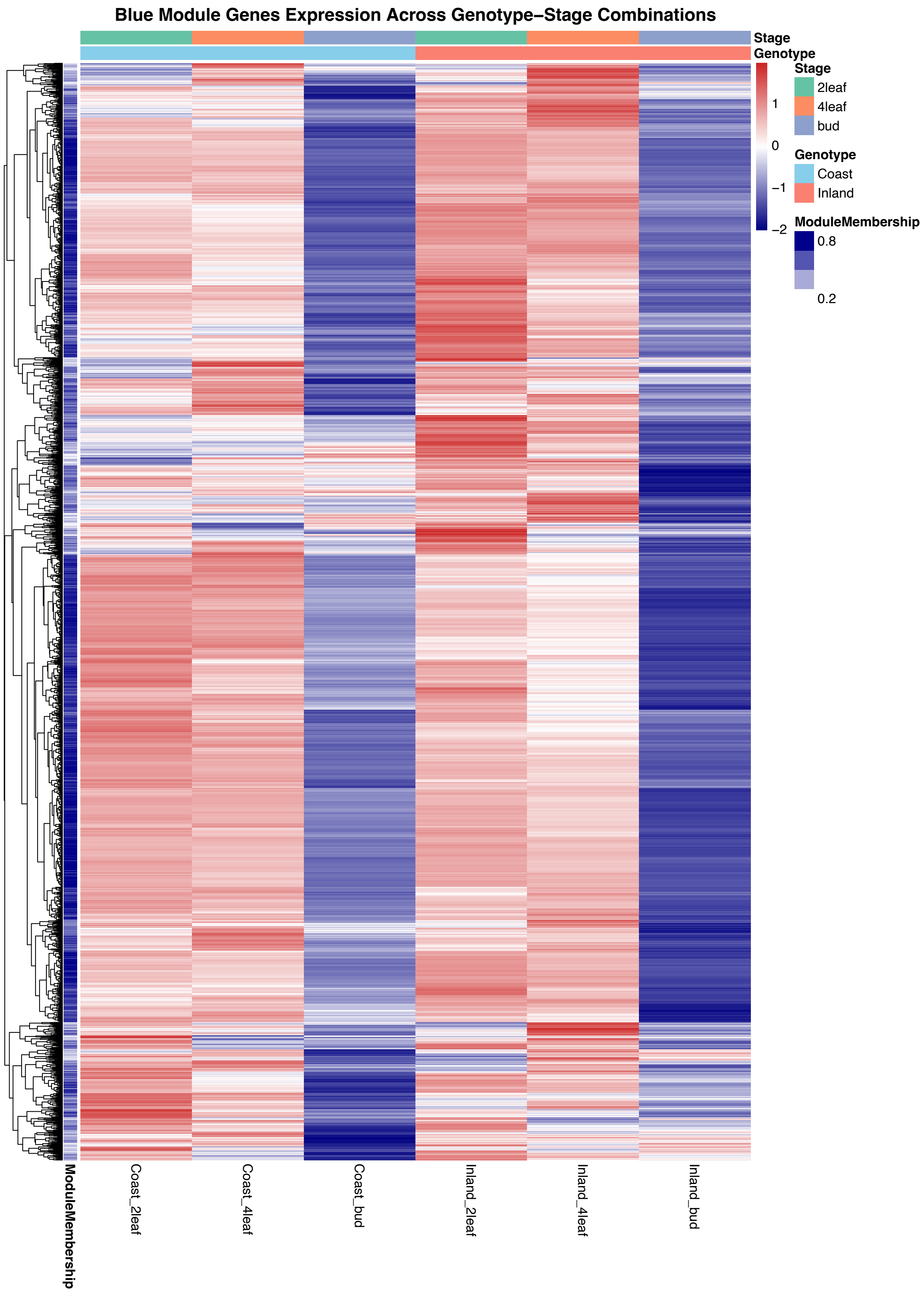


**Figure S26.** Heatmap of the expression of genes within the blue module across genotypes and developmental stages.

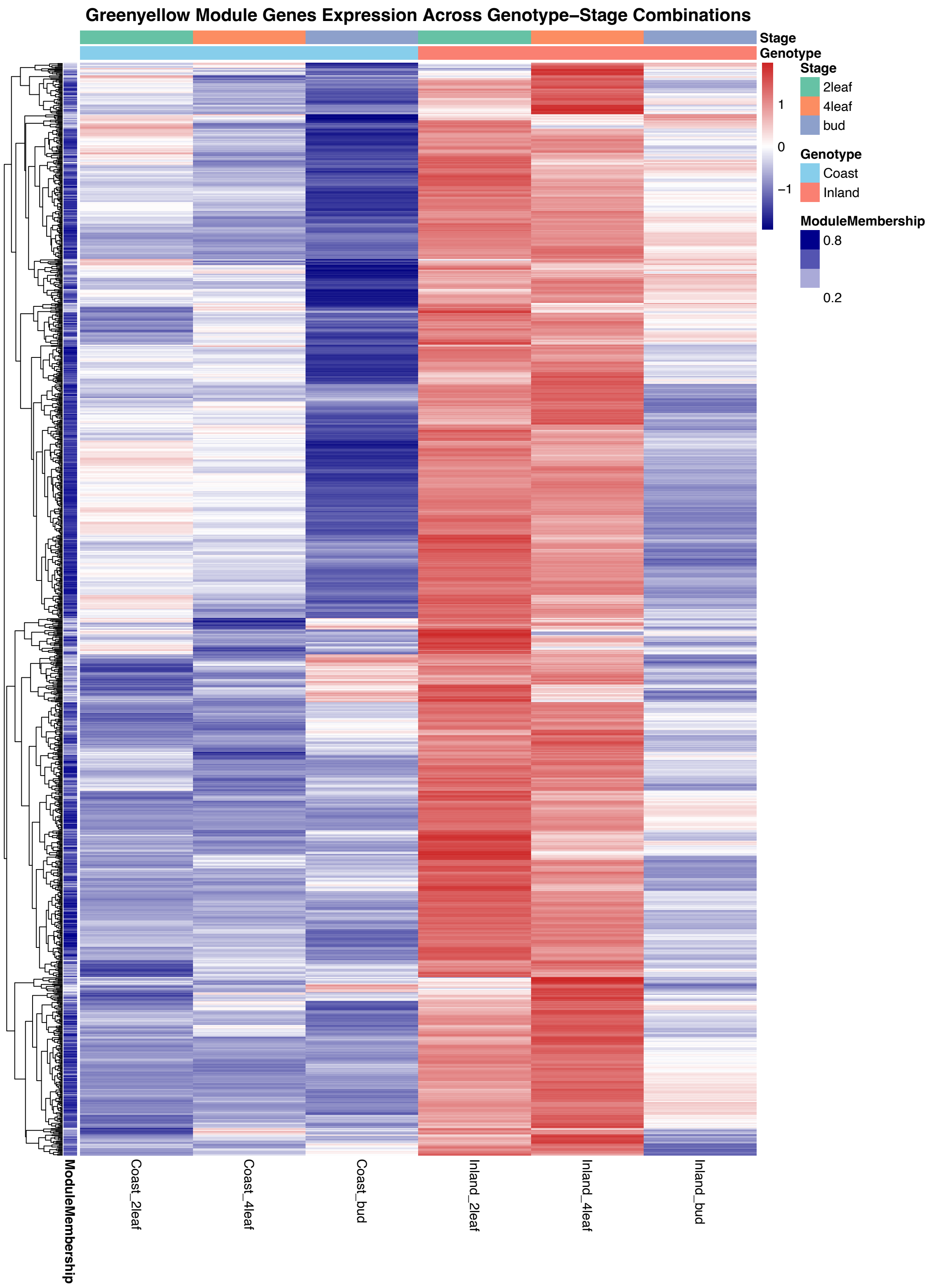


**Figure S27.** Heatmap of the expression of genes within the greenyellow module across genotypes and developmental stages.
